# TLR4 signaling drives tissue inflammation, Claudin-5 internalization, and vascular barrier breakdown in a mouse model of neonatal meningitis

**DOI:** 10.64898/2025.12.31.697193

**Authors:** Philip V. Seegren, Amir Rattner, Philip M. Smallwood, Yanshu Wang, Jeremy Nathans

**Author notes:** Address for editorial correspondence: Dr. Jeremy Nathans, 805 PCTB, 725 North Wolfe Street, Johns Hopkins University School of Medicine Baltimore, MD 21205.

## Abstract

Neonatal bacterial meningitis is a leading cause of infant morbidity and mortality, yet the molecular and cellular basis of the leptomeningeal response to infection remains poorly defined. Here, we study a mouse model of neonatal *E. coli* meningitis, combining conditional gene knockouts, leptomeningeal single-nucleus RNA sequencing, and endothelial cell culture to explore the role of Toll-like receptor 4 (TLR4) signaling in the host response to infection. Deletion of *Tlr4* in non-myeloid cells dramatically reduced the inflammatory response in all leptomeningeal cell types and abrogated the infection-associated increase in vascular permeability. In a brain endothelial cell line (bEnd.3 cells), exposure to *E. coli* triggered NF-κB activation, selective internalization of Claudin- 5, and increased monolayer permeability, responses that were eliminated by *Tlr4* knockout. RNA-seq showed that TLR4 controls an NF-κB–driven transcriptional program that orchestrates the endothelial response to *E. coli*. These findings reveal multiple TLR4-dependent host responses to neonatal Gram-negative bacterial meningitis.

## Introduction

Bacterial meningitis is a major global health challenge, causing ∼250,000 deaths annually (Brouwer, 2010; GBD 2016 Neurology Collaborators, 2019). Among neonates, Group B Streptococcus (*Streptococcus agalactiae*, GBS) and *Escherichia coli* K1 are the predominant causes of meningitis and are responsible for roughly 40% and 30% of cases, respectively (Ouchenir et al., 2017; Bundy et al., 2023). While vaccination has markedly reduced mortality in childhood meningitis, neonatal infections continue to carry high case-fatality rates, especially in low-income regions, and roughly half of the survivors experience lasting neurological deficits (Edmond et al., 2010; Guarnera et al., 2024). The high susceptibility of neonates likely reflects the immaturity of both the adaptive immune system and multiple CNS barriers, including the blood–brain barrier (BBB), the blood–cerebrospinal fluid (CSF) barrier, and the leptomeninges (Saunders et al., 2012; Yang et al., 2023). Understanding host–pathogen interactions in this developmental window is essential for designing strategies to protect the neonatal brain (Kim, 2003).

The leptomeninges form a surface barrier enveloping the brain and spinal cord. It contains a dense vascular network and large numbers of immune cells within the CSF-filled subarachnoid space. Within the leptomeninges, endothelial cells (ECs) possess BBB properties, including expression of Claudin-5 (Cldn5), Occludin, and transporters characteristic of CNS microvessels (Mastorakos et al., 2019). At present, the response of leptomeningeal ECs to bacterial infection, including alterations in barrier properties, is largely unexplored.

The present study focuses on the role of one innate immune receptor, Toll-like receptor 4 (TLR4), in regulating the leptomeningeal response to bacterial meningitis. TLR4 is a pattern-recognition receptor that detects lipopolysaccharide (LPS), the major outer membrane component of Gram-negative bacteria such as *E. coli* (Medzhitov, 2001; Kawai and Akira, 2007). Activation of TLR4 initiates MyD88- and TRIF-dependent signaling cascades, leading to the induction of proinflammatory cytokines, chemokines, and adhesion molecules that orchestrate leukocyte recruitment to infected tissues (Vallières and Rivest, 1997; Pålsson-McDermott and O’Neill, 2004; Nagyoszi et al. 2010; Casanova et al., 2011). In the context of bacterial meningitis, TLR4 has been shown to mediate both host-protective and pathological responses, including vascular inflammation and leukocyte infiltration (Van Well et al., 2013; Doran et al., 2016; Too et al., 2019; Wang et al., 2023). While these studies established TLR4 as a key driver of meningitis-associated inflammation, the molecular and cellular events associated with the TLR4-dependent host response during bacterial meningitis remain largely unexplored.

Claudin-5 (Cldn5) is a prominent tight-junction protein in brain ECs. Originally identified as an endothelial-specific component of tight junctions (Morita et al., 1999), Cldn5 was later shown to be indispensable for restricting paracellular permeability *in vivo*, as *Cldn5*-deficient mice exhibit size-selective leakage of small molecules across the BBB (Nitta et al., 2003). Under physiological conditions, Cldn5 is localized to endothelial cell–cell contacts, but its distribution and stability are dynamically regulated by endocytosis, recycling, and degradation pathways that respond to inflammatory stimuli (Engelhardt and Sorokin, 2009; Coisne and Engelhardt, 2011; Hashimoto et al., 2023). During CNS inflammation and infection, pro-inflammatory cytokines and innate immune receptor activation disrupt tight-junction continuity and alter endothelial barrier properties. This junctional remodeling includes Cldn5 internalization, but the upstream signaling pathways controlling Cldn5 trafficking remain incompletely understood.

A recent single-cell transcriptome study defined the genomic response of the leptomeninges in a mouse model of neonatal *E. coli* meningitis (Wang et al., 2023). Leptomeningeal ECs exhibited a clear inflammatory transcriptional signature, implicating the leptomeningeal vasculature as a locus of immune signaling during *E. coli* infection. Whether these signals directly control tight junction remodeling and vascular leak was not tested.

Here, we study a mouse neonatal meningitis model and cultured CNS ECs (bEnd.3 cells) to test whether non-myeloid TLR4 signaling orchestrates the multi-cellular response of the leptomeninges to infection and regulates Cldn5 trafficking and vascular barrier function. In the present work, we (i) used single nucleus (sn)RNA-seq to profile cell-type–specific responses in the leptomeninges from WT mice, endothelial and fibroblast (i.e., non-myeloid) *Tlr4* conditional knockout mice (*Tlr4^VEKO^*), and myeloid *Tlr4* conditional knockout mice (*Tlr4^MKO^*); (ii) quantified intravascular tracer leakage and immune pathway activation *in vivo*; (iii) tracked Cldn5 redistribution and NF-κB nuclear translocation in bEnd.3 cells exposed to *E. coli*; (iv) generated CRISPR *Tlr4^KO^* bEnd.3 lines to assess the TLR4 dependence of NF-κB activation, tight-junction dynamics, and barrier permeability; (v) quantified Cldn5 colocalization with junctional, plasma-membrane, and endosomal/lysosomal markers; and (vi) defined TLR4-dependent transcriptional programs in bEnd.3 cells exposed to *E. coli*. The results of these experiments establish a framework for understanding the pathophysiology of neonatal meningitis that establishes TLR4 signaling as a central regulator of both the multi-cellular leptomeningeal response to infection and Cldn5 trafficking and junctional integrity.

## Results

### Cell-type-specific transcriptional responses of the leptomeninges from WT, *Tlr4^VEKO^*, and *Tlr4^MKO^* mice to neonatal *E. coli* infection

Moving from brain to skull, the meninges consist of four distinct layers: (1) the pia, a thin, semi-permeable layer that mediates solute exchange between the CNS and CSF; (2) the subarachnoid space, a highly vascularized region that contains a high density of immune cells; (3) the arachnoid and its barrier layer, demarcating the boundary between CNS and non-CNS territories; and (4) the dura, a fibrous layer outside of the CNS with a high density of blood vessels, lymphatics, and immune cells (Figure 1A). The first three layers constitute the leptomeninges. Taking advantage of a natural cleavage plane between dura and leptomeninges, we have optimized whole mount immunostaining and confocal imaging of the leptomeninges together with the adjacent cortex, preserving the anatomical integrity of the pia and subarachnoid space (Figure 1B).

**Figure 1.**
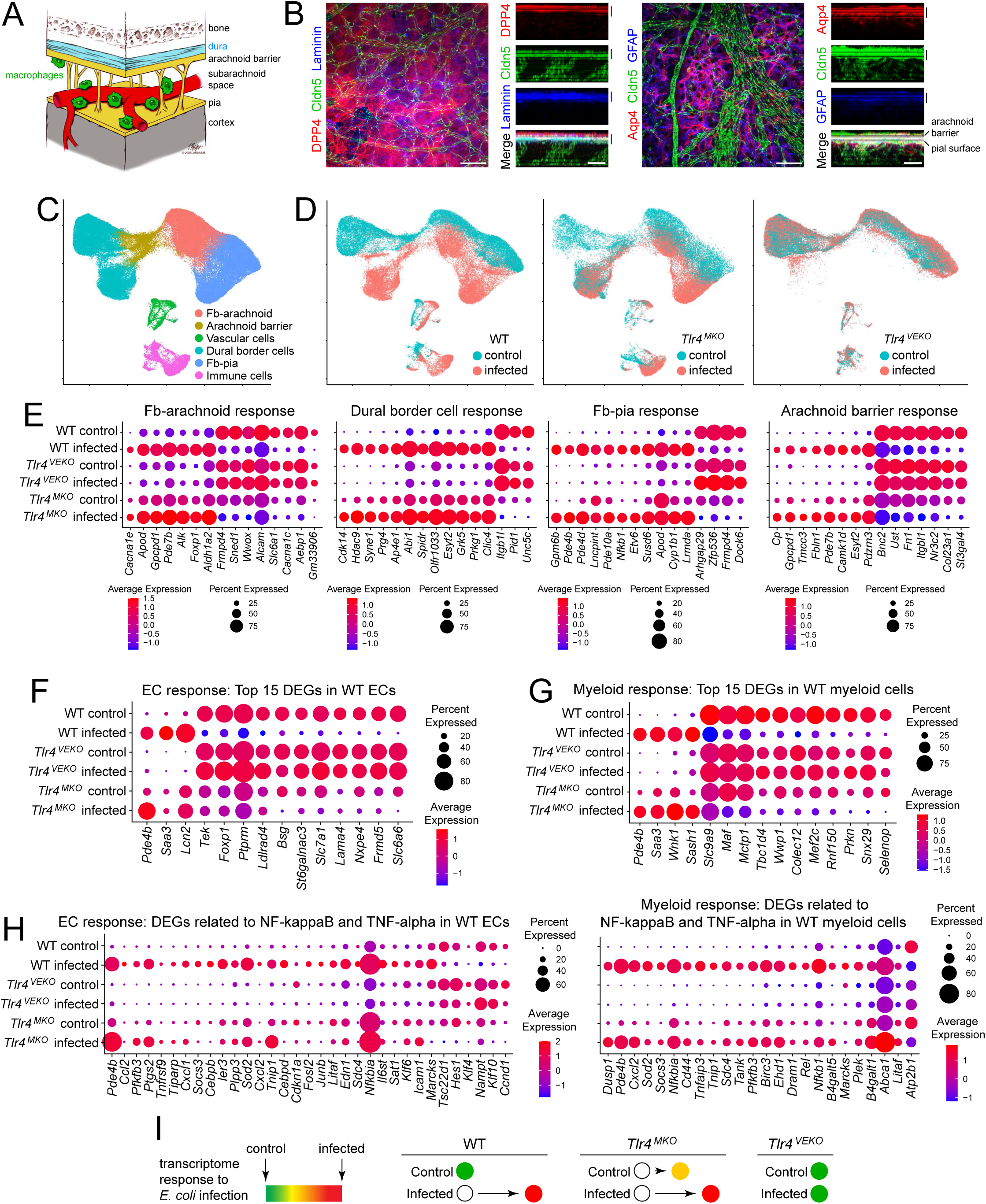
Cell-type-specific transcriptional responses of leptomeninges from WT, *Tlr4^VEKO^*, and *Tlr4^MKO^* mice to neonatal *E. coli* infection. (A) Schematic of the leptomeninges (pia, subarachnoid space, and arachnoid barrier) and adjacent structures. (B) Immunostaining of P6 mouse leptomeninges shown as a whole mount (left) or as a transverse section through a series of Z-planes (right). The vertical black bars indicate the extent of the leptomeninges. Scale bars, 20 µm. (C) UMAP plot of leptomeninges snRNA-seq from WT, *Tlr4^VEKO^*, and *Tlr4^MKO^* mice at P6, from uninfected and *E. coli*-infected mice, with cell clusters identified. N=126,235 nuclei. Fb, fibroblast. (D) UMAP comparison of P6 leptomeningeal snRNA-seq from control vs. 24-hour *E. coli*- infected mice of the indicated genotypes. (E) snRNA-seq from four leptomeninges cell clusters (Fibroblast-arachnoid, Dural border cells, Fibroblast-pia, and Arachnoid barrier cells) showing differentially expressed genes in control vs. *E. coli*-infected mice of the indicated genotypes at P6. For each cell cluster, DEGs were chosen based on a comparison of infected vs. uninfected WT cells. (F and G) As in panel (E) except for ECs (F) and myeloid cells (G). DEGs, differentially expressed genes. (H) As in panel (F) and (G) except DEGs were related to NF-κB and TNF-α signaling. (I) Summary of transcriptional responses of leptomeninges from WT, *Tlr4^VEKO^*, and *Tlr4^MKO^* mice to *E. coli* infection.

In the *E. coli* meningitis model used here, postnatal day (P)5 mice were given a subcutaneous injection of 20 µl of phosphate buffered saline (PBS) with 120,000 colony forming units of *E. coli* RS218 (O18:K1:H7), a clinical isolate from the CSF of a neonate with bacterial meningitis (Zhu et al., 2020; Wang et al., 2023) or, as a control, 20 µl of phosphate buffered saline (PBS). This *E. coli* strain was additionally transformed with a plasmid encoding a red fluorescent protein (RFP) to facilitate detection of bacteria in infected tissues. Infected and control mice were sacrificed at P6, 24 hours after inoculation. P5 mice were chosen to approximate the developmental stage of the human newborn BBB and immune system, as described in Wang et al. (2023). This model recapitulates the main temporal sequence of events in human neonates with bacterial meningitis: peripheral entry of bacteria (e.g., via skin abrasions during birth), hematogenous spread of bacteria (bacteremia), and bacterial seeding of the meninges.

To assess cell type-specific roles of TLR4 signaling, we used two conditional knockout lines: *Tlr4^VEKO^* (=*Cdh5-CreER;Tlr4^fl/−^*) and *Tlr4^MKO^* (=*Lyz2^Cre/+^;Tlr4^fl/−^*). *Cdh5* encodes Vascular Endothelial (VE) Cadherin, and the *Cdh5-CreER* line carries a *Cdh5* promotor-*CreER* transgene (Monvoisin et al., 2006). *Lyz2*, also referred to as *LysM*, encodes a myeloid-specific lysozyme, and the *Lyz2^Cre^* line is a *Cre* knock-in at the *Lyz2* locus (Clausen et al., 1999). The cell type specificities of *Cdh5-CreER* and *Lyz2^Cre^* were assessed using a *Rosa26 loxP-stop-loxP* reporter (*Rosa26-LSL-tdT-2A-nlsGFP*; Wang et al., 2018). The specificity of *Cdh5-CreER* was additionally assessed using a *Rosa26-LSL-SUN1-sfGFP* reporter (Mo et al., 2015).

Throughout most of the CNS, the *Cdh5-CreER* line directs Cre-mediated recombination specifically to ECs (e.g., Wang et al., 2025b). However, in the meninges, only ∼75% of the cells recombined by the *Cdh5-CreER* transgene are ECs and ∼25% are neither ECs nor myeloid cells (Figure 1 – figure supplement 1A, C, and D). Quantification of DAPI stained sections of leptomeninges shows that ECs (ERG+ cells) constitute 31+/-9% of leptomeningeal cells, and myeloid cells (PU.1+ cells) constitute 14+/-5% of leptomeningeal cells (Figure 1 – figure supplement 1B, C, and E). Therefore, dural border cells, arachnoid barrier cells, and fibroblasts together constitute the remaining ∼55% of leptomeningeal cells. Since the non-ECs and non-myeloid cells that express *Cdh5-CreER* are one-third as numerous as ECs, they constitute ∼10% of leptomeningeal cells, equivalent to ∼18% of the combined population of dural border cells, arachnoid barrier cells, and fibroblasts. These data imply that at P2, when the single dose of 4HT was administered, the *Cdh5-CreER* transgene produced Cre-mediated recombination in all or nearly all ECs, in a minority of dural border cells, arachnoid barrier cells, and fibroblasts, and not in myeloid cells, consistent with the pattern of *Cdh5* expression in leptomeninges described by others (Mapunda et al., 2023; Pietilä et al., 2023; Smyth et al., 2024). Extrapolating these data to *Tlr4*, *Tlr4^VEKO^* mice should be considered to have complete or nearly complete *Tlr4* knockout in ECs, with *Tlr4* additionally knocked out in a fraction of the other non-myeloid cell types. For simplicity, we will refer to the *Tlr4^VEKO^*KO pattern as “non-myeloid” KO of *Tlr4,* although the data in Figure 1 – figure supplement 1 imply that it is predominantly an EC KO.

Expression of the *Lyz2^Cre^* knock-in allele was restricted to myeloid cells within the leptomeninges as determined by immunostaining for three partially overlapping myeloid markers, CD206, ASC, and PU.1 (Figure 1 – figure supplement 2). *Lyz2^Cre^*directed Cre- recombination with 90-100% efficiency in CD206+ cells and with 50-70% efficiency in ASC+ and PU.1+ cells, the range depending on whether tdTomato or GFP colocalization was being scored (Figure 1 – figure supplement 2). In the text that follows, we will refer to *Lyz2^Cre^*- recombined cells simply as “myeloid cells”, although they should be understood as CD206+ myeloid cells. The low abundance of TLR4 and the limitations of commercial anti-TLR4 antibodies precluded a direct immunohistochemical assessment of TLR4 loss in *Tlr4^VEKO^* and *Tlr4^MKO^* mice.

To profile the transcriptional landscape of the leptomeninges in WT, *Tlr4^VEKO^*, and *Tlr4^MKO^*mice, with and without meningitis, we performed single-nucleus RNA sequencing (snRNA-seq) on dissected leptomeninges (Supplementary file 1). A uniform manifold approximation and projection (UMAP) analysis of the dataset revealed six major cell classes: Fb (fibroblast)-arachnoid1, Fb-pia, arachnoid barrier cells, dural border cells, vascular cells, and immune/myeloid cells (Figure 1C; Figure 1 – figure supplement 3A–C), consistent with prior studies (Pietilä et al., 2023; Wang et al., 2023; Smyth et al., 2024; we note that the cell cluster referred to in Wang et al., 2023 as “fibroblasts dura 3” corresponds to dural border cells). The same set of cell clusters was observed with both PIP-seq (Clark et al., 2023) and 10X Genomics platforms, which additionally confirmed expression of *Cdh5* in non-myeloid and non-EC leptomeningeal cell types and expression of *Lyz2* exclusively in myeloid cells (Figure 1 – figure supplement 4).

Comparison of infected and uninfected conditions showed that *Tlr4^VEKO^* mice exhibited a global attenuation of infection-induced transcriptional responses across all major leptomeningeal cell types, as judged by the positions of cell clusters in the UMAP (Figure 1D). Similar trends were observed in principal component analysis (PCA), where infected and uninfected WT samples were widely separated along the PC2 axis, whereas infected and uninfected *Tlr4^VEKO^* samples clustered closely together (Figure 1 – figure supplement 5). Uninfected *Tlr4^MKO^* transcriptomes localized roughly midway between infected and uninfected WT samples along PC2.

At the level of individual genes, all major leptomeningeal cell types displayed infection-associated transcriptional changes in which the most differentially expressed genes, based on a comparison between infected and uninfected WT mice, exhibited similarly responses in WT and *Tlr4^MKO^*mice but little or no response in *Tlr4^VEKO^* mice (Figure 1E-G). In WT and *Tlr4^MKO^* mice, infection upregulated NF-κB and TNF-α transcriptional responses, whereas *Tlr4^VEKO^*mice were largely devoid of these responses (Figure 1H and Figure 1 – figure supplement 3D). Similar trends were observed in dot plots showing transcriptional changes associated with JAK-STAT and Interferon (IFN)-ɣ signaling pathways (Figure 1 – figure supplement 5). Interestingly, leptomeningeal cells in uninfected *Tlr4^MKO^* mice exhibit a transcriptional state that partially mimics the infection response (Figure 1D-H and Figure 1 – figure supplements 5 and 6). We speculate that this *Tlr4^MKO^* phenotype could represent an increase in the baseline level of innate immune signaling in uninfected *Tlr4^MKO^* mice. Comparing snRNA-seq data sets for individual mice and individual genes confirms for ECs and myeloid cells the conclusions obtained from aggregated data (Figure 1 – figure supplement 7). Together, these findings define an *E. coli* meningitis-associated transcriptional program controlled by TLR4 signaling that drives inflammatory responses in all of the major leptomeningeal cell types (Figure 1I).

### TLR4 signaling drives ICAM1 induction, increased vascular permeability, and myeloid activation during neonatal *E. coli* infection

To examine the role of TLR4 signaling in meningitis-associated responses within the leptomeninges, we used whole mount imaging to assess markers of vascular permeability and immune activation. Intercellular adhesion molecule-1 (ICAM1) serves as a well-established marker of endothelial activation during infection and inflammation (Bui et al., 2020; Singh et al., 2023). Whole mount leptomeninges from P6 uninfected and *E. coli*-infected WT, *Tlr4^VEKO^*, and *Tlr4^MKO^* mice revealed robust ICAM1 upregulation in WT and *Tlr4^MKO^* mice, whereas *Tlr4^VEKO^* mice showed little change in ICAM1 levels (Figure 2A–B). These protein-level changes mirrored the genotype-specific transcriptional differences detected by snRNA-seq (Figure 2 – figure supplement 1A). TLR4-dependent ICAM1 upregulation was also observed in underlying cortical capillaries (Figure 2 – figure supplement 1B–C). For these and all other immunostaining experiments with mice or with cultured cells, the numbers of mice and the numbers of images used in the analyses are listed in Supplementary file 2.

**Figure 2.**
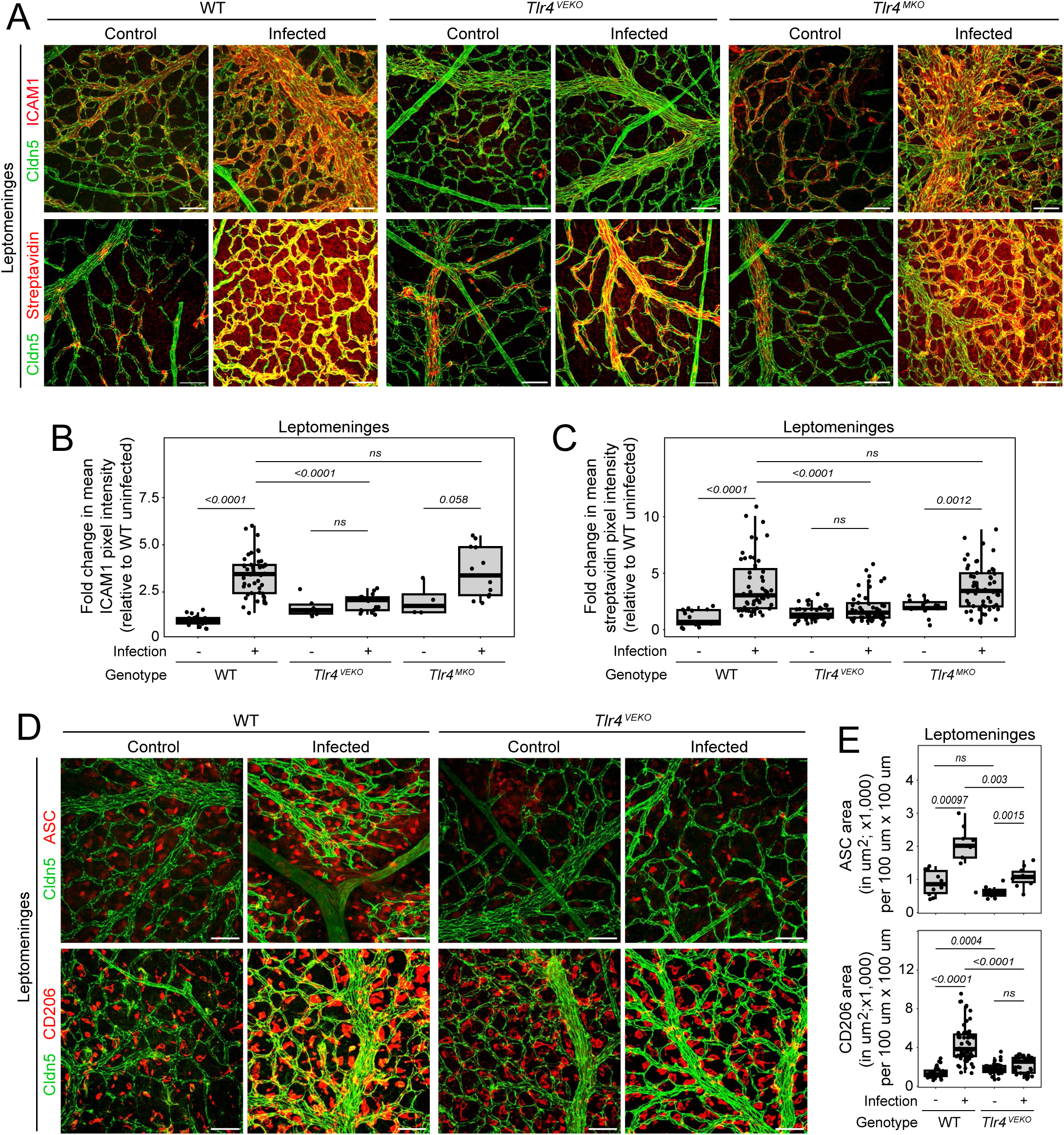
Non-myeloid TLR4 signaling drives ICAM-1 induction, increased vascular permeability, and myeloid cell activation during neonatal *E. coli* infection. (A) Whole mount leptomeninges from P6 WT, *Tlr4^VEKO^*, and *Tlr4^MKO^* mice, under control or *E. coli*-infected conditions, immunostained for Cldn5 and ICAM-1 (upper panels) or immunostained for Cldn5 and incubated with fluorescent streptavidin (lower panels) to visualize the fate of intravascular sulfo-NHS-biotin. Scale bars, 20 µm. (B) Quantification of ICAM1 immunostaining relative to WT controls, as shown in (A). (C) Quantification of extravascular streptavidin binding relative to WT controls, as shown in (A). (D) Whole mount leptomeninges from P6 WT and *Tlr4^VEKO^* mice, control or *E. coli*-infected, immunostained for Cldn5 and either ASC or CD206 to visualize myeloid cells. Scale bars, 20 µm. (E) Quantification of ASC+ area (upper) and CD206+ area (lower), as shown in (D). Box plots show the median, interquartile range, and all individual data points. Each data point represents one image from a single mouse, with two locations imaged per mouse. Statistical comparisons (p-values) were calculated with the Wilcoxon rank-sum test. ns, not significant (p > 0.05).

To assess BBB integrity within the leptomeninges, we performed intraperitoneal (IP) injections of sulfo-NHS-biotin, a low molecular weight intravascular tracer. In this assay, perivascular streptavidin staining serves as a readout of vascular leakage (Wang et al., 2019). WT and *Tlr4^MKO^* mice exhibited marked increases in sulfo-NHS-biotin extravasation upon *E. coli* infection, whereas *Tlr4^VEKO^* mice showed no change from the uninfected state (Figure 2A and C). The simplest interpretation of these data is that the site of sulfo-NHS biotin leakage is primarily vascular. However, we cannot exclude some contribution from increased arachnoid barrier permeability.

In addition to these endothelial responses, we examined alterations in myeloid cells within the leptomeninges and the underlying cortex. The ASC protein (Apoptosis-associated speck-like protein containing a CARD; ASC/PYCARD), an inflammasome adaptor that changes its oligomeric state and subcellular localization in response to inflammatory stimuli (Sester et al., 2016; Franklin et al., 2018), localizes to leptomeningeal myeloid cells and brain microglia (Figure 2D). Quantification of the area occupied by ASC+ cells revealed significant increases in infected WT mice within both the leptomeninges and cortex (Figure 2D and E; Figure 2 – figure supplement 1D–E), whereas infected *Tlr4^VEKO^* mice exhibited a more modest response. To further characterize myeloid morphology, we stained for CD206 (Figure 2D and E; Figure 2 – figure supplement 1F). Within the leptomeninges, and similar to ASC staining, the area occupied by CD206+ myeloid cells increased in infected WT mice but showed substantially smaller changes in infected *Tlr4^VEKO^* mice (Figure 2D–E). In the brain, CD206+ cells were too sparse to reliably quantify (Figure 2 – figure supplement 1F). The increase in area of ASC and CD206 immunostaining in the infected leptomeninges relative to controls largely reflects a shift of myeloid cells from a compact to a more extended morphology (Wang et al, 2023). These findings imply that non-myeloid TLR4 signaling promotes myeloid activation and associated morphologic changes during *E. coli* meningitis.

Immunostaining for cleaved Caspase-3+ cells in leptomeninges+cortex flat mounts revealed increased numbers of cleaved Caspase-3+ cortical cells within ∼30 um beneath the pial surface following *E. coli* infection in WT and *Tlr4^MKO^* mice, but not in *Tlr4^VEKO^*mice (Figure 2 – figure supplement 2A and B). Similar analyses in coronal sections extending ∼750 um into the cortex showed ∼2-fold greater numbers of cleaved Caspase-3+ cortical cells in WT mice compared to *Tlr4^VEKO^*mice (Figure 2 – figure supplement 2C and D). In coronal sections, the number of cleaved Caspase-3+ cells in the leptomeninges correlated with the number in the cortex (R=0.79) in both WT mice and *Tlr4^VEKO^* mice (Figure 2 – figure supplement 2E). Co-staining with neuronal (NeuN), glial (GFAP), EC (Cldn5), and myeloid (PU.1) markers revealed that cleaved Caspase-3 in the cortex localized to myeloid cells (Figure 2 – figure supplement 2F). Immunostaining for Iba1, a marker of microglial activation, showed increased numbers of cortical Iba1+ cells in both WT and *Tlr4^VEKO^* mice in response to infection, with a ∼2-fold greater number in WT mice compared to *Tlr4^VEKO^* mice (Figure 2 – figure supplement 3). The infection-dependent appearance of cleaved Caspase-3+ myeloid cells in the cortex could reflect the previously described non-apoptotic role of caspases in immune cell differentiation and activation (Sordet et al., 2002; Santambrogio et al., 2005). Together, these results indicate that non-myeloid TLR4 signaling drives both increased vascular permeability and myeloid cell activation during neonatal *E. coli* meningitis.

The loss of TLR4 signaling in *Tlr4^VEKO^* mice had little or no effect on the clinical course of the *E. coli* infection. The presence of *E. coli* in blood cultures and its dissemination to brain, liver, and lung occurred indistinguishably in infected WT and *Tlr4^VEKO^* mice (Figure 2 – figure supplement 4A-C). The most obvious clinical manifestation of *E. coli* infection is a failure to gain weight, an observation consistent across WT, *Tlr4^VEKO^*, *Tlr4^MKO^*, and *Tlr4^-/-^* mice (Figure 2 – figure supplement 4D; Wang et al., 2023).

### Claudin-5 redistribution in response to *E. coli* in leptomeningeal endothelial cells and in bEnd.3 cells

In our previous description of meningitis-associated responses in the leptomeninges, we observed a redistribution of Cldn5 independent of changes in total Cldn5 levels (Wang et al., 2023). Given the central role of Cldn5 in maintaining BBB integrity, we investigated whether TLR4-mediated inflammatory signaling contributed to these changes. Consistent with prior findings (Wang et al., 2023), both WT and *Tlr4^MKO^* mice showed an increase in the area occupied by Cldn5 in the leptomeninges following infection, likely referable to both increased vessel diameter and a redistribution of Cldn5 within ECs (Figure 3A-B; Figure 3 – figure supplement 1C). Similar to the effects of global *Tlr4* KO (Wang et al., 2023), *Tlr4^VEKO^*mice showed minimal redistribution of Cldn5 (Figure 3A-B). The area occupied by Zonula Occludens-1 (ZO-1), a tight junction scaffold protein, showed a modest but statistically significant increase in WT leptomeningeal vessels but no significant change in *Tlr4^VEKO^* leptomeningeal vessels during infection (Figure 3 – figure supplement 1A and B). In sum, these data suggests that non-myeloid TLR4 signaling regulates EC tight-junction organization.

**Figure 3.**
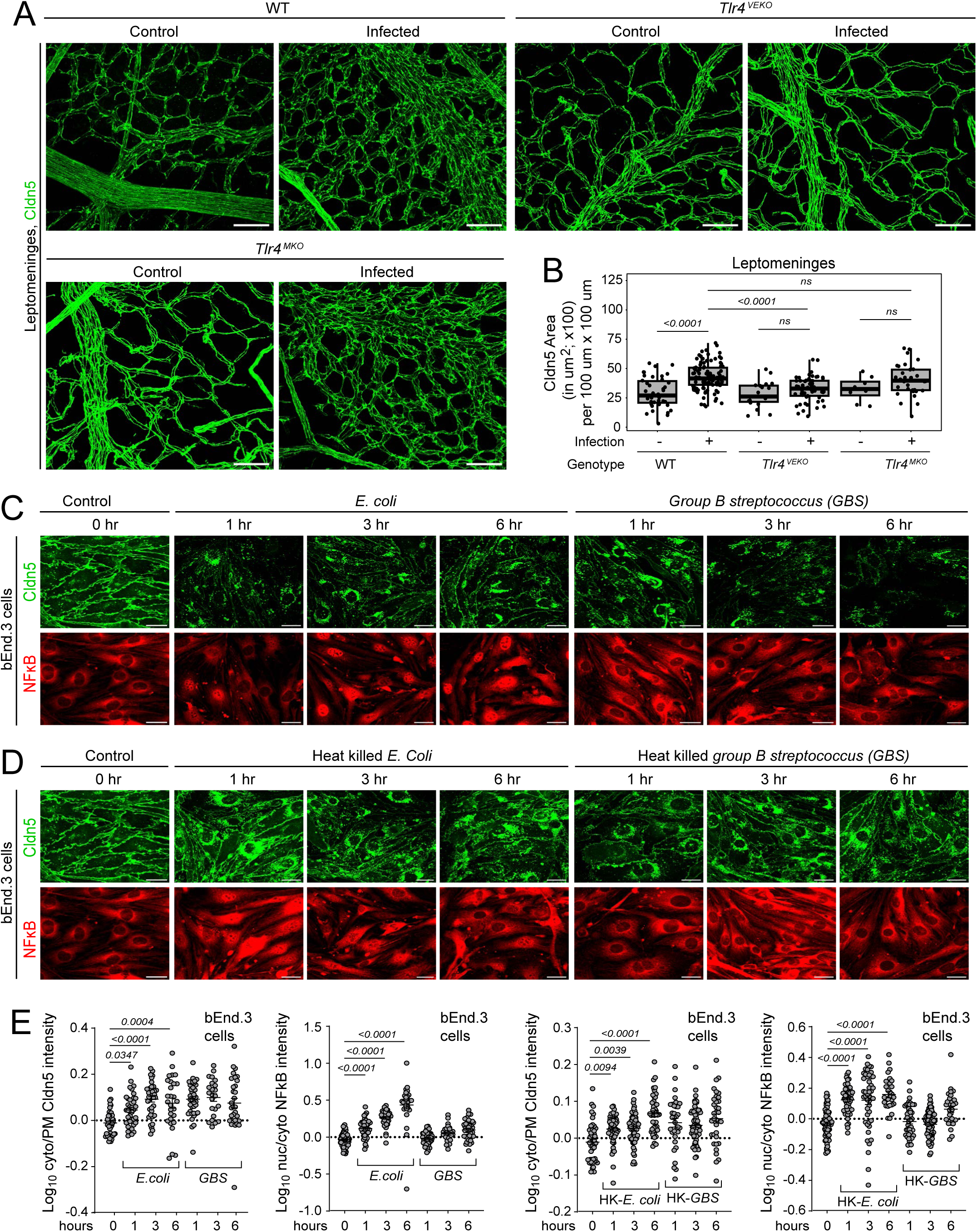
Claudin-5 redistribution in response to *E.coli* in leptomeningeal endothelial cells and in bEnd.3 cells. (A) Whole mount leptomeninges from P6 WT, *Tlr4^VEKO^*, and *Tlr4^MKO^*mice, control or *E. coli*- infected, immunostained for Cldn5. Scale bars, 20 µm. (B) Quantification of Cldn5+ area in the leptomeninges, as shown in (A). (C) Control bEnd.3 cells and bEnd.3 cells exposed to live *E. coli* or live *group B Streptococcus* (GBS) for 1–6 hours were immunostained for Cldn5 and NF-κB. Scale bars, 20 µm. (D) Control bEnd.3 cells and bEnd.3 cells exposed to heat-killed *E. coli* or heat-killed *group B Streptococcus* (GBS) for 1–6 hours were immunostained for Cldn5 and NF-κB. Scale bars, 20 µm. (E) Quantification of experiments shown in (C) and (D). First plot, quantification of log_10_- transformed cytoplasmic-to-plasma membrane (Cyto/PM) intensity ratio for Cldn5 in bEnd.3 cells exposed to live *E. coli* for the indicated time in hours. Second plot, quantification of log_10_- transformed nuclear-to-cytoplasmic (Nuc/Cyto) intensity ratio for NF-κB in bEnd.3 cells exposed to live GBS for the indicated time in hours. The third and fourth plots are analogous to the first and second plots, except that bEnd.3 cells were exposed to heat-killed (HK) *E. coli* or GBS. Box plot in (B) shows median, interquartile range, and all individual data points. Each *in vivo* data point represents one image from a single mouse, with two locations imaged per mouse. For plots in (E), each data point represents a single bEnd.3 cell analyzed from representative images in three biological replicates. Statistical comparisons (p-values) were calculated with the Wilcoxon rank-sum test. ns, not significant (p > 0.05).

To directly test the link between inflammatory signaling and Cldn5 redistribution, we turned to cultured mouse brain ECs (bEnd.3), which express both TLR4 and Cldn5 at baseline (Williams et al., 1988; Montesano et al., 1990; Sikorski et al., 1993). Although they are brain-derived, bEnd.3 cells lack many BBB-specific attributes and, therefore, they likely exhibit generalized endothelial responses rather than brain-specific responses to bacterial exposure. Time-course imaging of bEnd.3 cells exposed to living or heat-killed *E. coli* or *Group B Streptococcus* (GBS) revealed rapid redistribution of Cldn5 from the plasma membrane to intracellular compartments within one hour of bacterial exposure under all four conditions (Figure 3C, 3D, and 3E first and third plots). Nuclear translocation of NF-κB was greatest following exposure to live *E. coli*, beginning within 1 hour and progressively increasing over the 6-hour duration of the experiment, consistent with strong TLR4 activation (Figure 3C left half, and 3E second plot). Heat-killed *E. coli* produced a more modest nuclear translocation of NF- κB, peaking at 3–6 hours (Figure 3D left half, and 3E fourth plot). Despite the robust internalization of Cldn5 in response to either live or heat-killed GBS, the NF-κB response to GBS was minimal, suggesting that TLR2-mediated NF-κB signaling is weaker than TLR4- mediated signaling in bEnd.3 cells (Figure 3C and D right halves, and 3E second and fourth plots). Together, these findings indicate that Cldn5 redistribution is a rapid endothelial response to bacterial exposure that can occur in the absence of detectable NF-κB nuclear translocation, suggesting partial or complete independence from NF-κB–driven transcriptional responses.

### TLR4 signaling controls NF-κB activation, tight junction dynamics, and barrier integrity in bEnd.3 cells exposed to *E. coli*

To directly test the role of TLR4 signaling in bEnd.3 cells exposed to *E. coli*, we generated a CRISPR *Tlr4* knockout line (Figure 4). Tlr4 knockout lines were generated using Cas9 and sgTrack-mCherry-mediated sgRNA delivery (see Methods; Hulton et al., 2020). Exons 1 and 3 of the mouse *Tlr4* locus were targeted with sgRNAs and clonal lines were isolated by fluorescence-activated cell sorting. Functional loss of TLR4 was confirmed by quantifying NF-κB nuclear translocation following bacterial exposure (Figure 4 – figure supplement 1A). Sanger sequencing confirmed frameshift mutations in all *Tlr4* alleles in several exon 3-targeted clones, one of which (*Tlr4^KO^*-m1) was selected for further study (Figure 4 – figure supplement 1B and C). Immunostaining for Cldn5 and NF-κB in WT and *Tlr4^KO^* bEnd.3 cells showed that deletion of TLR4 prevented both NF-κB nuclear translocation and Cldn5 internalization in response to *E. coli* (Figure 4A–D).

**Figure 4.**
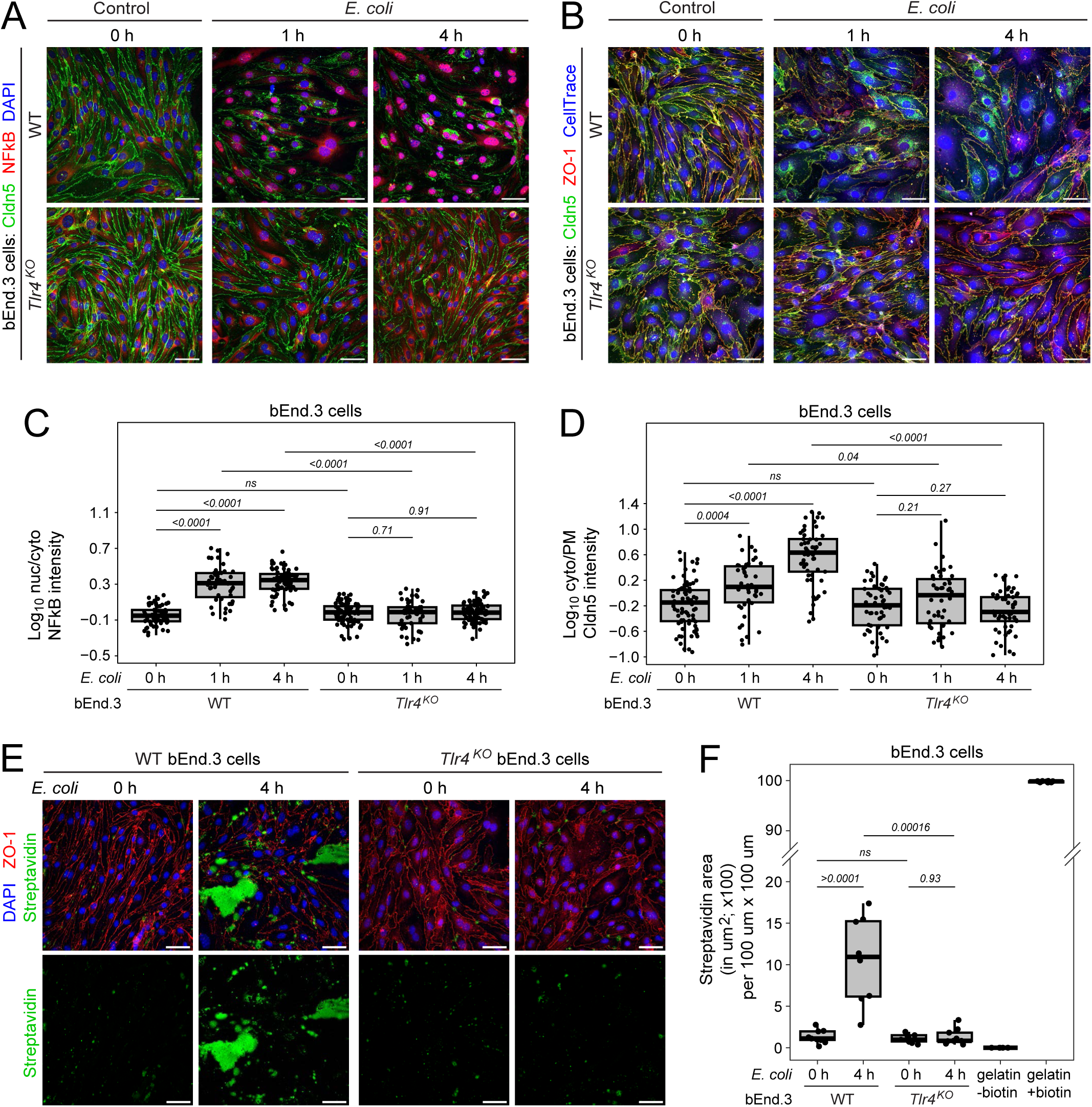
TLR4 signaling controls NF-κB activation, tight junction dynamics, and barrier integrity in bEnd.3 cells exposed to *E. coli*. (A) WT and *Tlr4^KO^* bEnd.3 cells, either not exposed to *E. coli* (control) or exposed to *E. coli* for 1 hour or 4 hours, were fixed and immunostained for Cldn5 and NF-κB and stained with DAPI. Scale bars, 20 µm. (B) WT and *Tlr4^KO^* bEnd.3 cells, either not exposed to *E. coli* (control) or exposed to *E. coli* for 1 hour or 4 hours, were fixed and immunostained for Cldn5 and ZO-1 and stained with CellTrace. Scale bars: 20 µm. (C) Quantification of log_10_-transformed nuclear-to-cytoplasmic (Nuc/Cyto) NF-κB intensity ratio in WT and *Tlr4^KO^* bEnd.3 cells, with or without exposure to *E. coli*, as shown in (A). (D) Quantification of log_10_-transformed cytoplasmic-to-plasma membrane (Cyto/PM) Cldn5 intensity ratio in WT and *Tlr4^KO^* bEnd.3 cells, with or without exposure to *E. coli*, using ZO-1 localization to define the plasma membrane region, as shown in (B). (E) WT and *Tlr4^KO^*bEnd.3 cells were grown to confluence on biotinylated gelatin-coated coverslips, with or without *E. coli* exposure for 4 hours, were then incubated with fluorescent streptavidin, and were finally fixed and immunostained for ZO-1. Representative images are shown with merged (upper) and streptavidin-only (lower) channels. Scale bars, 20 µm. (F) Quantification of streptavidin+ area in WT and *Tlr4^KO^* bEnd.3 cells with or without exposure to *E. coli*, as shown in (E). Wells coated with gelatin alone or biotinylated gelatin (without cultured cells) served as negative and positive controls, respectively. Box plots show median, interquartile range, and all individual data points. Each data point represents a single cell (C and D) or one image field (F) from representative images in three biological replicates. Statistical comparisons (p-values) were calculated with the Wilcoxon rank- sum test. ns, not significant (p > 0.05).

To test whether TLR4 signaling directly regulates endothelial barrier permeability, we used a cell culture assay to quantify barrier permeability across confluent bEnd.3 monolayers (Dubrovskyi et al., 2013). In this assay, bEnd.3 cells were grown to confluence on biotinylated gelatin and breaks in the monolayer of cells were visualized by adding fluorescent streptavidin to the medium. Cldn5-containing tight junctions were observed only in confluent bEnd.3 monolayers (Figure 4 – figure supplement 2A). Control experiments showed that the streptavidin signal depended on coating the plate with biotinylated gelatin and on gaps in the cell monolayer (Figure 4 – figure supplement 2B). When bEnd.3 cells were grown to confluence, the monolayer effectively restricted streptavidin access to the underlying biotinylated gelatin substrate (Figure 4 – figure supplement 2B). Application of this assay to WT and *Tlr4^KO^* bEnd.3 monolayers that were exposed to *E. coli* showed that loss of TLR4 signaling prevented streptavidin access to the underlying biotinylated gelatin substrate and preserved tight-junction integrity (Figure 4E and F). Together, these data indicate that TLR4 signaling in cultured brain ECs drives Cldn5 internalization and increases monolayer permeability *in vitro*, mirroring TLR4’s role *in vivo*.

### Comparisons of Cldn5 localization with junctional, plasma membrane, and trafficking markers in WT and *Tlr4^KO^* bEnd.3 cells with or without *E. coli* exposure

Next, we examined the subcellular localization and specificity of Cldn5 redistribution in WT and *Tlr4^KO^*bEnd.3 cells. We examined several tight junction-associated and integral membrane proteins expressed in bEnd.3 cells – ZO-1, β-catenin, glucose transporter 1 (GLUT1), and platelet endothelial cell adhesion molecule-1 (PECAM1) – to assess changes in their colocalization with Cldn5 during *E. coli* exposure (Figure 5A–5C). This analysis was conducted using confocal microscopy with a 40X objective; thus, the designation of “colocalization” is at the resolution conferred by this objective. Cellular colocalization was quantified using percent-overlap analysis in ImageJ (see Methods). At baseline, ZO-1 and β- catenin showed an average of ∼75% overlap with Cldn5, consistent with their colocalization at endothelial tight junctions (Figure 5D, Figure 5 – figure supplement 1A). Both ZO-1 and β- catenin remained plasma membrane-localized after 4 hours of *E. coli* exposure, while Cldn5 was redistributed to cytoplasmic compartments in WT but not *Tlr4^KO^*cells (Figure 5A, 5D, Figure 5 – figure supplement 1A).

**Figure 5.**
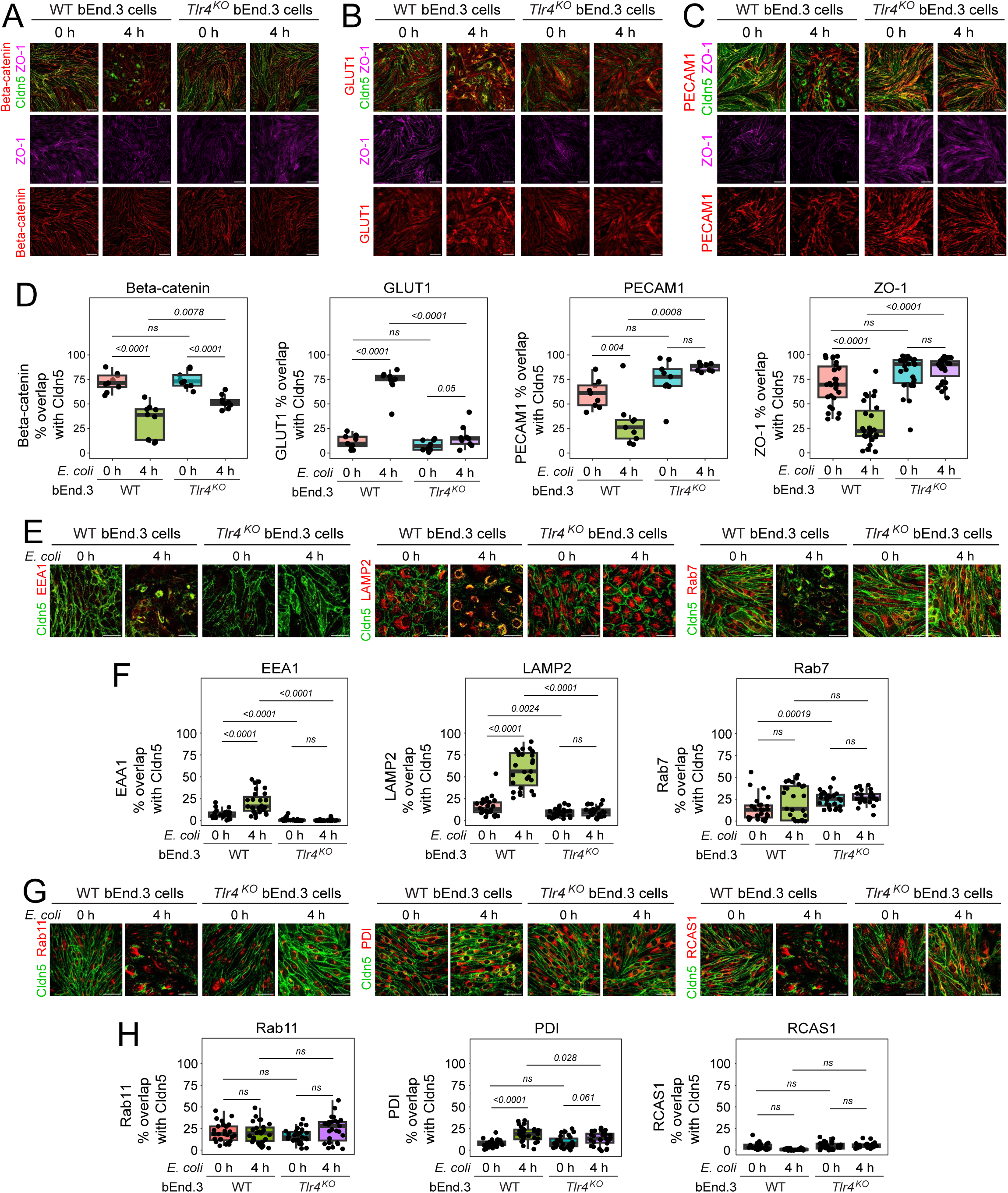
Comparisons of Cldn5 localization with junctional, plasma membrane, and trafficking markers in WT and *Tlr4^KO^* bEnd.3 cells with or without *E. coli* exposure. (A-C) WT and *Tlr4^KO^* bEnd.3 cells, either not exposed to *E. coli* (control) or exposed to *E. coli* for 4 hours, were fixed and immunostained for Cldn5 and the indicated markers. Scale bars, 20 µm. (D) Quantification of overlap between Cldn5 and β-catenin, GLUT1, PECAM1, and ZO-1, with each data point representing a 100 µm x 100 µm region of interest (ROI). (E) WT and *Tlr4^KO^* bEnd.3 cells, either not exposed to *E. coli* (control) or exposed to *E. coli* for 4 hours, were fixed and immunostained for Cldn5 and the indicated markers. Scale bars, 20 µm. (F) Quantification of overlap between Cldn5 and EEA1, LAMP2, and Rab7, as in (D). (G) WT and *Tlr4^KO^* bEnd.3 cells, either not exposed to *E. coli* (control) or exposed to *E. coli* for 4 hours, were fixed and immunostained for Cldn5 and the indicated markers. Scale bars, 20 µm. (H) Quantification of overlap between Cldn5 and Rab11, PDI, and RCAS1, as in (D). Box plots show median, interquartile range, and all individual data points. Each data point represents one image from representative images in three biological replicates. Statistical comparisons (p-values) were calculated with the Wilcoxon rank-sum test. ns, not significant (p > 0.05).

We next analyzed non-tight junction membrane proteins. The glucose transporter GLUT1 showed relatively little overlap with Cldn5 at baseline and localized primarily to the plasma membrane (Figure 5B). In these images, GLUT1 immunostaining was present across the cell but it did not reveal a silhouette of the nucleus, implying that it was localized to the plasma membrane rather than the cytoplasm. During *E. coli* exposure, GLUT1 was internalized alongside Cldn5, showing ∼75% overlap within cytoplasmic compartments and revealing the silhouette of the nucleus (Figure 5D, Figure 5 – figure supplement 1A). In contrast, PECAM1, another non-tight junction surface marker, retained its plasma-membrane localization during the 4-hour exposure to *E. coli* in both WT and *Tlr4^KO^*cells (Figure 5C, 5D, Figure 5 – figure supplement 1A). Taken together, these findings reveal distinct specificities of internalization among membrane and membrane-associated proteins dependent on TLR4 signaling in cultured brain endothelial cells.

To identify the intracellular compartments involved in Cldn5 trafficking, we co-stained WT and *Tlr4^KO^* cells for Early Endosome Antigen 1 (EEA1), Lysosome-associated Membrane Protein 2 (LAMP2), Rab7, and Rab11 to label endosomal and lysosomal vesicles, and for Protein Disulfide Isomerase (PDI) and Receptor-binding Cancer Antigen Expressed on SiSo Cells (RCAS1) to mark the endoplasmic reticulum and Golgi apparatus, respectively (Figure 5E-H) (Stamatovic et al., 2009, 2017; Scott et al., 2014; Shearer and Peterson, 2019; MacDonald et al., 2020). After 4 hours of exposure to *E. coli*, Cldn5 colocalization with LAMP2⁺ vesicles increased to ∼50% overlap in a TLR4-dependent manner (Figure 5E and 5F). *E. coli* exposure also produced small increases in Cldn5 colocalization with EEA1 and PDI, to ∼20% overlap, which was similarly dependent on TLR4 (Figure 5E-H). *E. coli* exposure produced little change in the low-level colocalization of Cldn5 with Rab7, Rab11, or RCAS1, and there was little or no effect of TLR4 loss (Figure 5E-H). We interpret the TLR4-dependent increase in Cldn5-EEA1 and Cdn5-LAMP2 overlap as representing the endocytic internalization of Cldn5 and its delivery to lysosomes. We interpret the modest increase in Cldn5-PDI as likely arising from the close proximity of endosomes and lysosomes to the ER, which is present throughout the cytoplasm.

Further insights into the mechanism of Cldn5 endocytosis came from visualizing the internalization of transferrin, cholera toxin subunit B (CTB), and 10-kDa dextran, well-established tracers for clathrin-mediated endocytosis, caveolin-mediated endocytosis, and macropinocytosis, respectively (Figure 6A; Johnson and Spence, 2010). Prior to *E. coli* exposure, these tracers and Cldn5 exhibited ∼80% overlap at the plasma membrane (Figure 6B). Following *E. coli* exposure, reduced overlap with transferrin and CTB, together with persistent overlap with 10-kDa dextran, suggested a macropinocytosis-like mechanism of Cldn5 internalization (Figure 6B). The persistence of intracellular Cldn5 immunostaining with up to at least 6 hours of *E. coli* exposure, together with washout experiments showing restoration of plasma membrane Cldn5 localization within 3 hours of *E. coli* removal, suggest little or no degradation of internalized Cldn5 during this time frame (Figure 6C and D). Together, these results indicate that TLR4 signaling triggers a highly specific and reversible internalization of Cldn5, providing a potential mechanistic link between inflammatory activation and increased vascular permeability.

**Figure 6.**
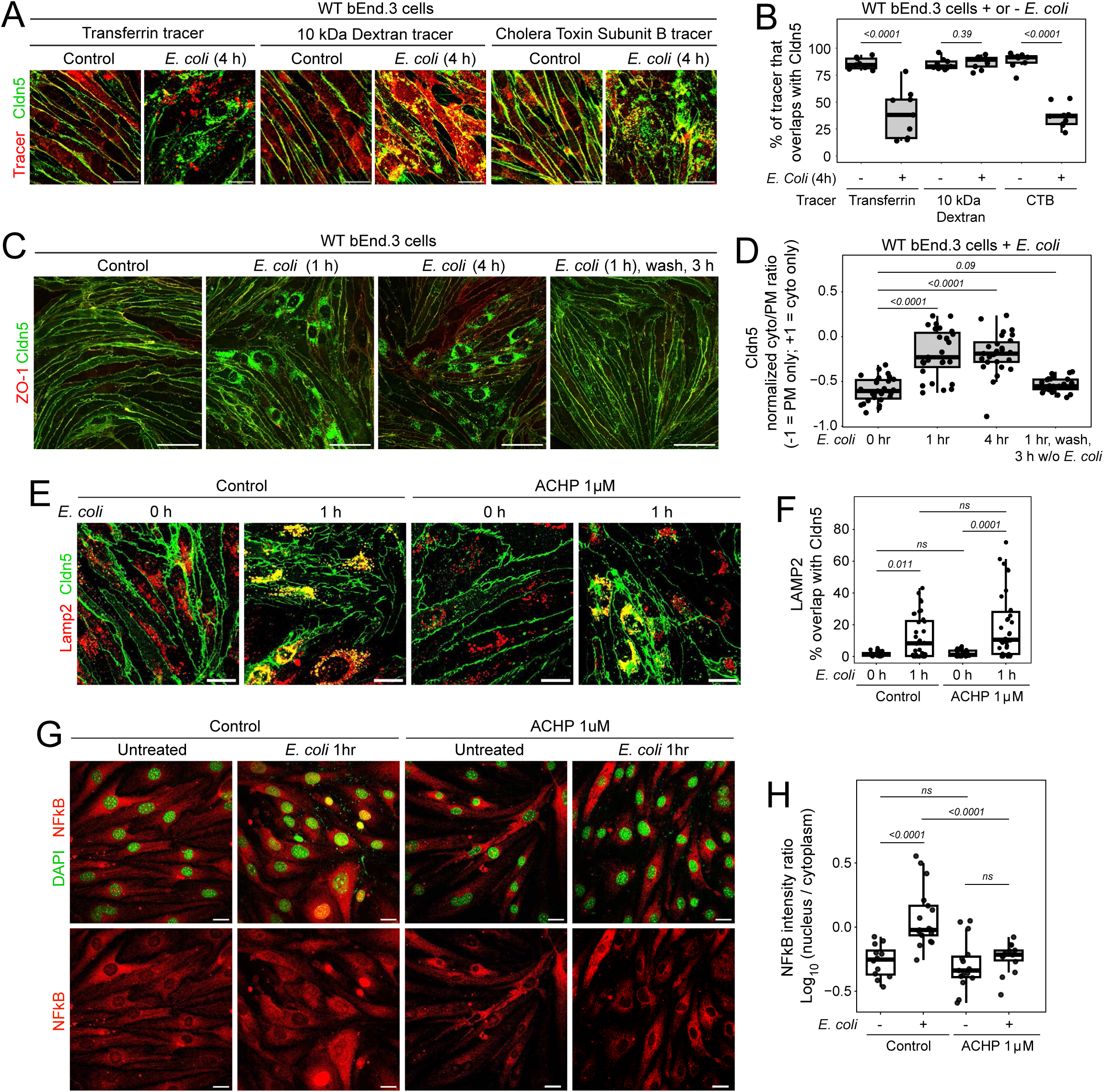
In bEnd.3 cells, pharmacologic inhibition of NF-κB nuclear translocation and signaling does not impede rapid internalization of Cldn5 in response to *E. coli* exposure. (A) bEnd.3 cells were pre-incubated for 30 minutes with the indicated fluorescent tracer, then either not exposed to *E. coli* (control) or exposed to *E. coli* for 4 hours and then fixed and immunostained for Cldn5. Scale bars, 20 µm. (B) Quantification of % overlap between tracer and Cldn5 from the experiment shown in (B). (C) Cldn5 internalization and recovery during 1 hour of *E. coli* exposure, followed by washout of *E. coli*, and then an additional 3 hours of incubation in medium without bacteria. Scale bars, 50 µm. (D) Quantification of the experiment shown in (D). The metric on the y-axis is (cytoplasmic signal – plasma membrane signal)/(cytoplasmic signal + plasma membrane signal). 100% cytoplasmic localization corresponds to 1.0. 100% plasma membrane localization corresponds to -1.0. 50% cytoplasmic localization and 50% plasma membrane localization corresponds to 0.0. (E) The subcellular localization of Lamp2 and Cldn5 were determined in bEnd3 cells in the presence or absence of 1 µM of the IKK inhibitor ACHP before *E. coli* exposure or after 1 hour of *E. coli* exposure. Scale bars, 20 µm. (F) Quantification of the experiment shown in (A). Each symbol represents a 100 µm x 100 µm region of interest (ROI). (G) NF-κB translocation from cytoplasm to nucleus was visualized in the presence or absence of 1 µM of ACHP before *E. coli* exposure or after 1 hour of *E. coli* exposure. Scale bars, 20 µm. (H) Quantification of the experiment shown in (C). Each symbol represents an individual cell. Statistical comparisons (p-values) were calculated with the Wilcoxon rank-sum test. ns, not significant (p > 0.05).

To test the possible role of NF-κB signaling in Cldn5 internalization, we measured Cldn5 co-localization with Lamp2 in bEnd.3 cells after one hour of exposure to *E. coli* with or without the IκB kinase inhibitor ACHP (2-Amino-6-[2-(cyclopropylmethoxy)-6-hydroxyphenyl]-4-(4- piperidinyl)-3 pyridinecarbonitrile) at 1 µM (Figure 6E and F). Cldn5 internalization and co- localization with Lamp2 were insensitive to ACHP-mediated inhibition of NF-κB signaling under conditions that inhibited NF-κB nuclear translocation (Figure 6G and H). These data imply that the TLR4-dependent internalization of Cldn5 operates via an arm of the TLR4 pathway that is both rapid and distinct from NF-κB mediated changes in gene expression.

### Role of TLR4 signaling in the transcriptional response to *E. coli* in bEnd.3 cells

To define the TLR4-dependent transcriptional response to *E. coli* exposure in ECs, we performed bulk RNA-seq on WT and *Tlr4^KO^*bEnd.3 cells with or without 3 hours of *E. coli* exposure (Figure 7). Principal component analysis revealed separation of datasets based on both bacterial exposure and genotype (WT vs. *Tlr4^KO^*) (Figure 7 – figure supplement 1A and B). Interestingly, WT and *Tlr4^KO^* bEnd.3 cells that were not exposed to *E. coli* clustered separately, suggesting a basal level of TLR signaling even in the absence of *E. coli* exposure. Differential expression analysis in WT bEnd.3 cells showed robust induction of proinflammatory transcripts upon *E. coli* exposure, including *Il6*, *Cxcl2*, *Tnf*, *Icam1*, and *Ccl2* (Figure 7A), whereas this response was largely absent in *Tlr4^KO^* bEnd.3 cells (Figure 7B; note the change in axis scales between Figures 7A and 7B). Pathway analysis showed that inflammatory signaling was highly TLR4-dependent: TNF-α signaling via NF-κB, IL6–JAK–STAT signaling, and general inflammatory response pathways showed strong transcriptional activation in WT but not in *Tlr4^KO^* bEnd.3 cells (Figure 7C–F).

**Figure 7.**
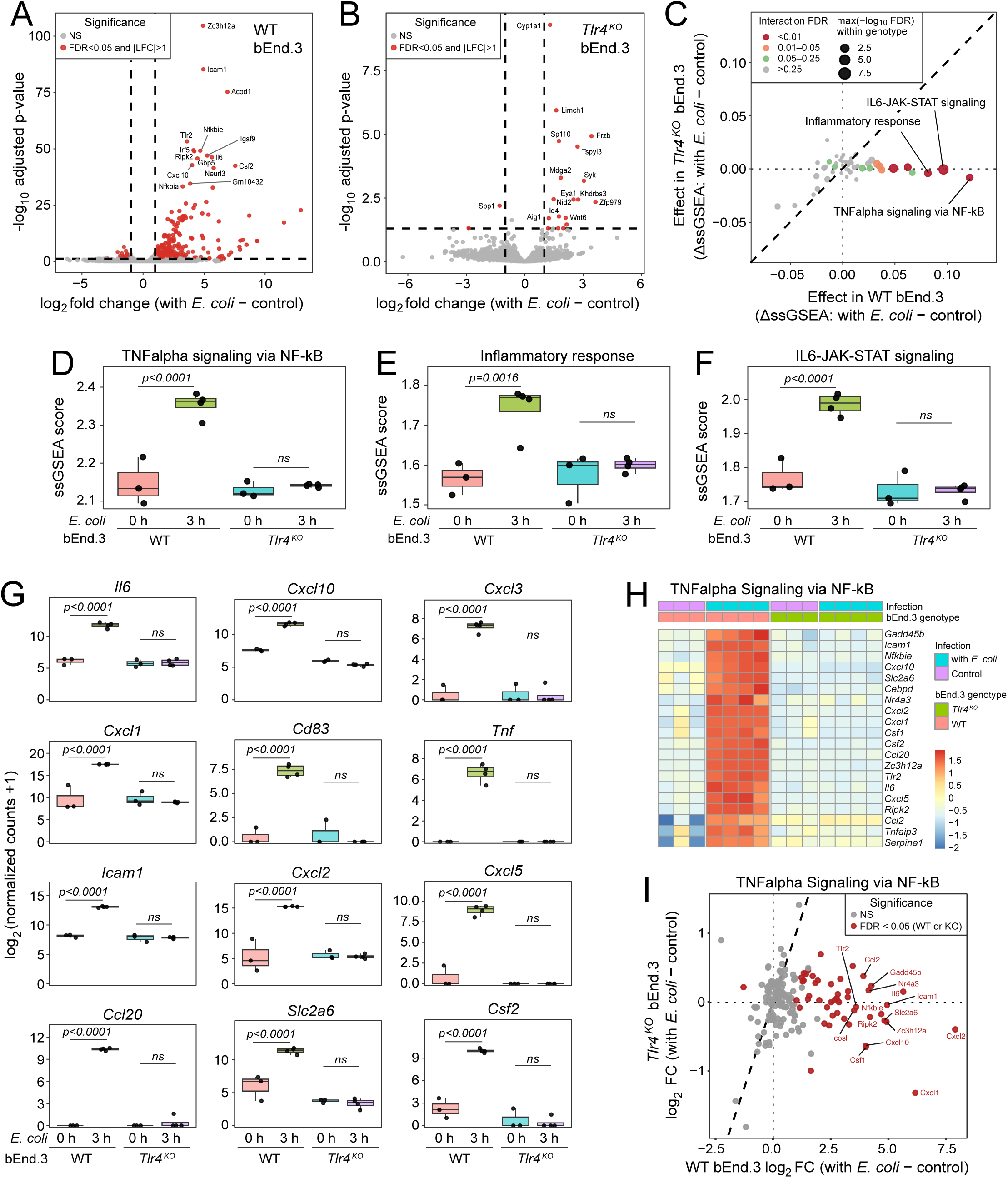
In bEnd.3 cells, TLR4 signaling plays a central role in the transcriptional response to *E. coli* exposure. (A) Volcano plot of differential transcript abundances in WT bEnd.3 cells, with or without exposure to *E. coli* for 3 hours. The plot shows transcript-level log_2_ fold change (LFC) on the horizontal axis versus −log_10_(false discovery rate; FDR) on the vertical axis. Red points mark significantly changed transcripts (FDR < 0.05 and |LFC| ≥ 1); grey points are not significant. (B) Volcano plot of differential transcript abundances in *Tlr4^KO^* bEnd.3 cells, with or without exposure to *E. coli* for 3 hours, plotted as in (A). Note the change of scale of the vertical and horizontal axes between (A) and (B). (C) Pathway effect map from ssGSEA Hallmark scores. Each symbol is a pathway. The horizontal axis is the effect of *E. coli* exposure for 3 hours on WT bEnd.3 cells (ΔssGSEA = *E. coli* exposed – not exposed) and the vertical axis is the effect of *E. coli* exposure for 3 hours on *Tlr4^KO^* bEnd.3 cells. Symbol size encodes the within-genotype significance [max(−log_10_ FDR) across both genotypes], and the color code indicates the genotype x *E. coli* exposure interaction FDR. The 45-degree diagonal line represents equal effects for the two genotypes. (D–F) ssGSEA pathway activity based on genotype and *E. coli* exposure for three Hallmark immune response pathways: TNF-α signaling via NF-κB (D), Inflammatory response (E), and IL6–JAK–STAT signaling (F). Symbols represent ssGSEA scores for individual samples. The vertical axis shows the interaction FDR score from a linear model for genotype x *E. coli* exposure. (G) Transcript abundance changes for 12 immune system genes in WT and *Tlr4^KO^* bEnd.3 cells, with or without exposure to *E. coli* for 3 hours. (H) Heatmap of transcript abundance changes in WT and *Tlr4^KO^* bEnd.3 cells, with or without exposure to *E. coli* for 3 hours, for TNF-α signaling via NF-κB. Rows correspond to transcripts with the greatest changes; values are variance-stabilized transformation (VST) z-scores derived from DESeq2-normalized counts. Columns are arranged by genotype and condition. (I) Scatterplot showing transcript abundance changes in WT and *Tlr4^KO^* bEnd.3 cells, with or without exposure to *E. coli* for 3 hours for TNF-α signaling via the NF-κB pathway. Each symbol represents one gene. Red points indicate genes significant for either genotype (FDR < 0.05). The top 15 genes by effect size are labeled. Note the different scales for the horizontal and vertical axes. The diagonal line represents equal effects for the two genotypes.

We next quantified transcripts for representative cytokines, chemokines, and adhesion molecules that define endothelial activation. *Il6*, *Cxcl10*, *Ccl2*, *Cxcl5*, *Cd83*, *Ccl20*, and *Icam1* transcripts were all significantly upregulated in WT bEnd.3 cells following *E. coli* exposure but remained unchanged in *Tlr4^KO^* bEnd.3 cells (Figure 7G). Heatmap visualization of the Hallmark “TNF-α signaling via NF-κB” gene set confirmed that NF-κB–responsive genes, including *Nfkbia*, *Ccl2*, *Il6*, and *Tnfaip3*, were selectively induced in WT cells by *E. coli* exposure (Figure 7H). Direct fold change comparisons between genotypes showed that induction of NF-κB target genes was strongly attenuated in *Tlr4^KO^* cells (Figure 7I).

To assess TLR4-dependent pathway activation following *E. coli* exposure, we performed single-sample gene set enrichment analysis (ssGSEA) across 50 Hallmark pathways. The pathways most strongly affected by genotype x exposure interactions included IL6–JAK–STAT3 signaling, allograft rejection, inflammatory response, TNF-α signaling via NF-κB, and the interferon-ɣ response (Figure 7 – figure supplement 1C). Heatmaps of Leading-Edge genes in these pathways further highlighted the broad suppression of cytokine and cell adhesion transcript induction in *Tlr4^KO^* bEnd.3 cells compared to WT (Figure 7 – figure supplement 1D–F).

Together, the *in vivo* and bEnd.3 transcriptional analyses identify TLR4 as a dominant regulator of inflammatory gene expression in response to *E. coli* exposure.

## Discussion

This study demonstrates that, in a mouse model of neonatal *E. coli* meningitis, non-myeloid TLR4 signaling is a key determinant of the inflammatory response in all major leptomeningeal cell types and of the infection-associated increase in vascular permeability. Using conditional *Tlr4* knockout models, leptomeningeal single-nucleus RNA sequencing, and *Tlr4* knockout in bEnd.3 cells, we show that TLR4 activation directs molecular, cellular, and transcriptional reprogramming of the vascular barrier. Our quantitative analysis of *Cdh5- CreER*-mediated recombination, showing recombination in all leptomeningeal ECs but in only a minority of other nonmyeloid cells, implies that the blunted infection response in the *Tlr4^VEKO^* leptomeninges is likely referable to loss of TLR4 signaling in ECs. In cell culture, endothelial TLR4 activation triggers rapid, selective, and reversible Cldn5 internalization, coincident with increased endothelial permeability. At the transcriptional level, endothelial TLR4 controls NF-κB and IL6/JAK/STAT inflammatory programs and cytokine gene expression. These findings establish TLR4 as the central leptomeningeal sensor of gram-negative bacteria, linking innate immune recognition to barrier dysfunction and to the tissue-wide inflammatory response.

### The leptomeningeal response to bacterial infection

Our snRNA-seq analyses revealed a dramatic transcriptome response to neonatal *E. coli* infection across all major leptomeningeal cell types that was broadly attenuated with non-myeloid deletion of *Tlr4*. Non-myeloid TLR4 signaling was required for the induction of canonical immune activation markers, including ICAM1 and leukocyte-recruiting chemokines. In contrast, myeloid-specific inactivation of *Tlr4* with *Lyz2-Cre* had minimal effect in the mouse meningitis model. Taken together, these data point to innate immune sensing as the gatekeeper of immune escalation in the leptomeninges. It is possible that myeloid TLR4 signaling in the leptomeninges may augment non-myeloid signaling or become relevant later in the course of the infection. We note that these experiments do not distinguish between local vs. distal anatomic sources of LPS or downstream effector molecules, such as cytokines, as activators of the leptomeningeal inflammatory response (Huang et al., 2021). Histologic observations of RFP-expressing *E. coli* in the brain, liver, and lungs, together with positive blood cultures, indicate substantial systemic dissemination of *E. coli* in this model. Thus, the inflammatory responses of leptomeningeal cells likely reflect exposure to bacterial products and inflammatory mediators derived from both local and distal sources.

In bEnd.3 cells, TLR4 activation triggers a rapid (≤1 hour) and reversible internalization of Cldn5, and *Tlr4^KO^*in bEnd.3 cells prevents both NF-κB translocation from cytoplasm to nucleus and Cldn5 redistribution. With TLR4 activation, ZO-1 and β-catenin remain junctional, PECAM1 remains at the plasma membrane, and GLUT1 co-internalizes with Cldn5, implying protein-specific re-localization. Percent-overlap and tracer analyses support endocytic routing of Cldn5, with prominent LAMP2+ (i.e., endosomal/lysosomal) localization, in a pattern that argues against canonical clathrin/caveolin pathways and is most consistent with macropinocytosis-like uptake. Rapid Cldn5 internalization occurred independently of NF-κB nuclear translocation, suggesting regulation through a non-transcriptional arm of TLR4 signaling. Strikingly, washout of *E. coli* restores Cldn5 to cell-cell junctions within three hours. These observations are broadly aligned with prior work showing cytokine/chemokine-evoked endocytosis of tight-junction proteins (including claudins and occludin) during EC barrier opening, via Rho/ROCK or caveolae-dependent routes (Stamatovic et al., 2006, 2009, 2012). Functionally, Cldn5 internalization was accompanied by increased barrier permeability, as measured by streptavidin passage across bEnd.3 monolayers or by Sulfo-NHS-biotin leakage *in vivo*. These findings reveal an EC-intrinsic process linking TLR4 activation to tight-junction remodeling and increased vascular permeability. In classic BBB tight-junction biology, Cldn5 is considered a gatekeeper of size-selective permeability (Morita et al., 1999). These findings extend that picture by identifying Cldn5 internalization as a disease-induced process that is rapid, selective, and reversible.

### Meningitis and vascular permeability

Mouse models of neonatal *E. coli* meningitis have shown conflicting effects of global TLR signaling on the clinical course of infection. Krishnan et al. (2013) reported that global loss of TLR4 resulted in a more rapid downhill course, whereas Jia et al. (2025) reported that global loss of TLR4 conferred substantial protection. Interestingly, Krishnan et al. (2013) reported that global loss of TLR2 conferred substantial protection in the same early postnatal *E. coli* meningitis model. The observations reported here are intermediate between those of Krishnan et al. (2013) and Jia et al. (2025), as we observed no obvious differences in two measures of disease progression (bacterial burden and body weight) between infected WT and *Tlr4^VEKO^* mice at 24 hours post-infection, despite substantial blunting of infection-associated gene expression and permeability changes in the leptomeninges of *Tlr4^VEKO^* mice. These divergent findings may reflect differences in experimental models and/or the fact that global TLR deletion affects multiple compartments simultaneously, including immune cell pathogen sensing, systemic inflammatory responses, and vascular barrier regulation.

Multiple pathways and mechanisms have been proposed to explain the increase in vascular permeability in the leptomeninges following bacterial infection. In an EC monolayer model of *E. coli* infection, activation of protein kinase C-alpha (PKC-alpha) led to disassembly of VE-cadherin mediated cell-cell junctions and a concomitant increase in monolayer permeability (Sukumaran and Prasadarao, 2003). In a similar EC monolayer model of *E. coli* infection, NOD-like receptor family pyrin domain containing 6 (NLRP6), a regulator of caspase, NF-κB, and inflammasome signaling, was found to reduce *E. coli*-induced tight junction disruption while also increasing *E. coli*-induced cell death (Jia et al., 2026). Wang, et al. (2025a) found that RIPK1, a kinase and scaffolding protein that integrates signals from inflammatory and cell death pathways, plays a central role in endothelial cell death and BBB breakdown during *E. coli* meningitis. Huang et al. (2024) showed that Wnt/beta-catenin signaling, which controls BBB-specific gene expression and is essential for maintaining BBB integrity, is inhibited by NF-κB signaling; augmenting Wnt/beta-catenin signaling blunted LPS-induced vascular leakage while reducing Wnt/beta-catenin signaling had the opposite effect. Yang et al. (2024) observed IL22-mediated degradation of occludin, ZO-1, and Cldn5 in cultured ECs, and showed that IL22 acts via a VEGF-STAT3 pathway to increase vascular permeability; in a mouse model of *E. coli* meningitis, this effect was largely abrogated by knocking out *Il22.* Finally, models of LPS-induced and *Klebsiella pneumoniae*-induced BBB breakdown implicate a Caspase-11 pathway that leads to the induction of Gasdermin D pores in CNS EC plasma membranes, leading to BBB breakdown and eventual pyroptotic cell death of ECs (Wei et al., 2024). Thus, the emerging picture is one of multiple inflammatory and cell death pathways converging on endothelial hyper-permeability.

We speculate that increased endothelial permeability in the leptomeninges could represent a double-edged response, deleterious when sustained and/or excessive, yet potentially protective when transient, facilitating the regulated passage of immune mediators and immune cells that are needed to combat infection. More generally, increases in vascular permeability may represent a general response to CNS stress or injury. For instance, others have observed reduced BBB integrity in brain regions remote from a focal optic-nerve injury, following transient global ischemia, in ischemic stroke, and in LPS-driven systemic inflammation (Banks et al., 2015; Ju et al., 2018; Smith et al., 2018; Cottarelli et al., 2025).

Activation of TLR4 signaling and the resulting increase in vascular permeability has direct clinical implications for neonatal meningitis. In particular, the data are consistent with a model in which endothelial TLR4 signaling contributes causally to CNS injury by promoting loss of barrier integrity, thereby permitting inflammatory cytokines and bacterial products to diffuse into the CNS parenchyma. These findings align with clinical observations in infants with bacterial meningitis, where elevated CSF cytokines (e.g., IL-6, TNF-α) and albumin level correlate with poor neurologic outcomes (Garges et al., 2006; Xu et al., 2019; Gao and Hu, 2024). By demonstrating the centrality of endothelial-intrinsic TLR4 signaling, our data highlight the endothelium not merely as a passive target but as a primary effector of neuropathology. These observations raise the possibility that selective modulation of endothelial TLR4 or more proximate regulators of Cldn5 trafficking could preserve vascular integrity, offering a potential avenue for neuroprotection in neonatal meningitis.

### Limitations of the study

Several limitations of the present study should be noted. While bEnd.3 cells provide a useful endothelial model, they do not exhibit the distinctive structural and molecular complexity of the blood–brain barrier (Garcia-Gallardo and Campbell, 2025). In particular, the bEnd.3 culture system lacks pericytes and astrocytes. Additionally, studying fixed cells and tissues does not permit detailed analyses of molecular and cellular dynamics. Going forward, live cell imaging of CNS ECs in culture (cell lines and/or acutely isolated cells) and *in vivo* two-photon imaging of the leptomeninges would complement the present studies by revealing dynamic responses in real time. Finally, the present study focused on *E. coli* K1, the dominant Gram-negative neonatal pathogen. Future work could assess TLR signaling in response to other bacterial pathogens, such as Group B *Streptococcus*.

### Concluding remarks

In sum, this study shows that TLR4 signaling couples innate immune recognition to inflammatory responses in multiple leptomeningeal cell types, inducing endothelial Cldn-5 internalization and increasing vascular permeability. The rapid and reversible nature of the Cldn5 internalization response implies that tight-junction remodeling is a dynamically regulated process. These results highlight potential opportunities to preserve vascular barrier properties by reducing endothelial TLR4 signaling, stabilizing Cldn5 in tight junctions, or dampening IL6– JAK–STAT and NF-κB signaling in endothelial cells (Freitas et al., 2018; Gadina et al., 2018; Zaffaroni and Peri, 2018; Hashimoto et al., 2021).

## Materials and Methods

### Key Resources Table

| Reagent type or resources | Designation | Source or reference | Additional information |
| --- | --- | --- | --- |
| Genetic reagent (*M. musculus*) | *Tlr4*<sup>-/-</sup> | JAX, stock #029015 |  |
| Genetic reagent (*M. musculus*) | *Tlr4*<sup>fl/fl</sup> | JAX, stock #024872 |  |
| Genetic reagent (*M. musculus*) | *Lyz2*<sup>Cre</sup> | JAX, stock #004781 |  |
| Genetic reagent (*M. musculus*) | *Cdh5*<sup>CreER</sup> | Monvoisin et al., 2006 |  |
| Genetic reagent (*M. musculus*) | *TLR4*<sup>MKO</sup> | *Tlr4*<sup>fl/fl</sup> X *Lyz2*<sup>Cre/+</sup>;<br>*Tlr4*<sup>-/-</sup> | *Lyz2*<sup>Cre/+</sup>; *Tlr4*<sup>fl/-</sup> |
| Genetic reagent (*M. musculus*) | *TLR4*<sup>VEKO</sup> | *Tlr4*<sup>fl/fl</sup> X<br>*Cdh5*<sup>CreER</sup>; *Tlr4*<sup>-/-</sup> | *Cdh5*<sup>CreER/+</sup>; *Tlr4*<sup>fl/-</sup> |
| Genetic reagent (*M. musculus*) | *Rosa26-LSL-tdT-2A-nlsGFP* | JAX, stock #030867 |  |
| Genetic reagent (*M. musculus*) | *Rosa26-LSL-SUN1-sfGFP* | JAX, stock #021039 |  |
| Genetic reagent (*E. coli*) | *E. coli* RS218 (O18:K1:H7) – RFP plasmid | Zhu et al., 2020 |  |
| Genetic reagent (recombinant DNA reagent) | sgTrack-mCherry | Addgene #114013 |  |
| Genetic reagent (recombinant DNA reagent) | Lenti-Cas9-Blast | Addgene #52962 |  |
| Cell Line (*M. musculus*) | bEnd.3 | ATCC CRL-2299 |  |
| Antibody | Mouse anti-Claudin-5 | ThermoFisher #352588 | (1:400) |
| Antibody | Mouse anti-ZO-1 | ThermoFisher #339194 | (1:400) |
| Antibody | Rabbit anti-ICAM-1 | ThermoFisher #14-0542-85 | (1:400) |
| Antibody | Rat anti-CD206 | Bio-Rad MCA2235 | (1:400) |
| Antibody | Rabbit anti-ASC/TMS1 | Cell Signaling Technology #67824S | (1:400) |
| Antibody | Rat anti-PECAM-1 | BD Pharmingen #553370 | (1:400) |
| Antibody | Rabbit anti-GLUT1 | ThermoFisher MA5-31960 | (1:400) |
| Antibody | Rabbit anti-cleaved Caspase-3 | Cell Signaling Technology, #9664 | (1:400) |
| Antibody | Rabbit anti-Iba1/AIF-1 | Cell Signaling Technology #17198 | (1:400) |
| Antibody | Rabbit anti-NFkB | Cell Signaling Technology #8242 | (1:400) |
| Antibody | Mouse anti-PU.1 | Cell Signaling Technology #39988 | (1:400) |
| Antibody | Rabbit anti-Rab7 | Cell Signaling Technology #9367 | (1:400) |
| Antibody | Rabbit anti-ERG | Cell Signaling Technology #97249 | (1:400) |
| Antibody | Rabbit anti-Rab11 | Cell Signaling Technology #5589 | (1:400) |
| Antibody | Rabbit anti-Lamp2 | Cell Signaling Technology #34141 | (1:400) |
| Antibody | Rabbit anti-PDI | Cell Signaling Technology #3501 | (1:400) |
| Antibody | Rabbit anti-EEA1 | Cell Signaling Technology #48453 | (1:400) |
| Antibody | Rabbit anti-RCAS1 | Cell Signaling Technology #12290 | (1:400) |
| Antibody | Rat anti-DPPIV/CD26 | R&D Systems #AF954 | (1:400) |
| Antibody | Rabbit anti-Aqp4 | Cell Signaling Technology #59678S | (1:400) |
| Antibody | Chicken anti-GFAP | Aves #GFAP | (1:400) |

### Mouse Models and *E. coli* Infection

Mice were maintained on a C57BL/6J background. Non-myeloid *Tlr4* conditional knockout mice (referred to as *Tlr4^VEKO^*) were generated by crossing *Tlr4^fl/fl^* (JAX, stock #024872) females with *Cdh5-CreER/+*;*Tlr4^−/−^* males, yielding *Cdh5-CreER/+*;*Tlr4^fl/–^*offspring. Myeloid-specific conditional knockout mice (*Tlr4^MKO^*), were generated by crossing *Tlr4^fl/fl^* females with *Lyz2^Cre/+^;Tlr4^−/−^*males, yielding *Lyz2^Cre/+^;Tlr4^fl/–^* offspring. We note that for both conditional knockout mouse lines, only a single Cre-mediated recombination event was required to convert *Tlr4^fl/-^* heterozygous cells to *Tlr4^-/-^* homozygous cells. For both sets of crosses, littermates lacking Cre were used as controls. The *Cdh5-CreER* line (Monvoisin et al., 2006) was the same line used in Wang. et al. (2025a). The *Lyz2^Cre^* line (JAX, stock #004781) was described in Clausen et al. (1999). [For the present study, the *Lyz2^Cre^* line was chosen based on the observations of Abram et al (2014) that it gives the most efficient myeloid-specific Cre-recombination among twelve Cre lines tested. We note that McKinsey et al (2020) have found that *PF4^Cre^* also recombines LoxP targets efficiently in leptomeningeal myeloid cells.] The *Rosa26-LSL-tdT-2A-nlsGFP* reporter line (JAX, stock #030867) was described in Wang et al. (2018). The Rosa26-LSL-SUN1-sfGFP reporter line (JAX, stock #021039) was described in Mo et al. (2015). Neonatal mice were genotyped by PCR and used for meningitis experiments without sex selection. Meninges snRNA-seq experiments and histological studies used postnatal day 6 (P6) mice with age-matched controls. To activate Cre-mediated recombination in progeny from the cross with *Cdh5-CreER*, 4-hydroxytamoxifen (4HT; Sigma-Aldrich H7904) was dissolved at a final concentration of 2 mg/ml in 1:10 vol:vol ethanol:sunflower seed oil (Sigma-Aldrich S5007), stored in aliquots at -80°C, and administered by intraperitoneal (IP) injection of 40-50 µl at P2. Mice were housed and handled according to the approved Institutional Animal Care and Use Committee protocol of the Johns Hopkins Medical Institutions (protocol MO22M375).

### Neonatal *E. coli* K1 infection model

To model neonatal meningitis, postnatal day 5 (P5) mice were used, approximating the developmental stage of the human newborn blood–brain barrier and immune system as described in Wang et al. (2023). Litters were randomized to receive a subcutaneous inoculation in the dorsal skin of 1.2 × 10⁵ colony-forming units (CFU) of *E. coli* RS218 (O18:K1:H7) with an RFP-expression plasmid (Zhu et al., 2020) in 20 µL PBS or 20 µL PBS alone. Twenty-four hours later (at P6), mice were euthanized for tissue collection. Brains, meninges, and relevant tissues were harvested, fixed, and processed for immunofluorescence and tracer leakage assays.

### Antibodies for tissue staining

Leptomeningeal and cortical whole mounts were immunostained with the following primary antibodies: mouse anti-Claudin-5 (Alexa Fluor™ 488, ThermoFisher #352588), mouse anti-ZO- 1 (Alexa Fluor™ 594, ThermoFisher #339194), rabbit anti-ICAM-1 (CD54, eBioKAT-1, ThermoFisher #14-0542-85), rat anti-CD206 (Bio-Rad MCA2235), rabbit anti-ASC/TMS1 (Cell Signaling Technology #67824S), rat anti-PECAM-1 (BD Pharmingen #553370), rabbit anti- GLUT1 (ThermoFisher MA5-31960), rabbit anti-cleaved Caspase-3 (Cell Signaling Technology, #9664), rabbit anti-ERG (Cell Signaling Technology, #97249). Alexa dye-conjugated secondary antibodies were from ThermoFisher.

### Sulfo-NHS-biotin leakage

For analysis of vascular leakage, mice were injected intraperitoneally with Sulfo-NHS-biotin (30 µl of 20 mg/mL Sulfo-NHS-biotin in PBS per mouse; ThermoFisher 21217) 15 minutes before sacrifice. After IP injection, the tracer rapidly equilibrates into the systemic circulation.

### Immunostaining of leptomeninges whole mounts

For whole mount imaging of the leptomeninges and adjacent cortex, a preparation was developed to preserve these structures in their native configuration. After euthanasia, brains (with leptomeninges attached) were fixed by immersion in 1% paraformaldehyde in PBS overnight at 4°C. The forebrain was then dissected and hemisected, and a thin slice of tissue from the surface of the brain was obtained by making a 300 µm vibratome section parallel to the surface of the cortex. Tissue slices were permeabilized in 1% Triton X-100/PBS before immunostaining, and blocked with 5% goat or donkey serum, depending on the secondary antibody host species. Primary antibodies were applied at 1:400 dilution, followed by fluorophore-conjugated secondary antibodies. Vascular leakage in mice injected with Sulfo-NHS-biotin was visualized by staining with streptavidin–Alexa Fluor 488 (Thermo Fisher, #S11223).

### Confocal microscopy and image acquisition

All images were acquired on a Zeiss LSM 780 laser-scanning confocal microscope controlled by Zeiss Zen software using 20x, 40x, or 63x/1.4 NA oil-immersion objectives. Fluorophores were excited with 405 nm, 488 nm, 561 nm, and 633 nm laser lines, and channels were collected sequentially to minimize spectral overlap. Z-stacks were acquired with step sizes optimized for each staining set; thinner stacks were used for cultured cell monolayers and deeper stacks for leptomeningeal whole mounts. The pinhole was set to 1 Airy unit. Laser power, gain, and offset settings were maintained at constant values within each experimental cohort (e.g. comparisons across genotypes). For leptomeninges imaging, two fields were imaged per hemisphere. Maximum intensity projections or single optical sections were exported from Zen for quantification and figure preparation.

### Blood bacterial burden (CFU assay)

Neonatal mice (P6; 24 hours post-infection) were euthanized on ice and whole blood was collected by cardiac puncture into tubes containing EDTA to prevent coagulation. Samples were kept on ice, and 20 µL of whole blood was diluted in 100 µL of LB broth, briefly vortexed, and plated onto LB agar plates. Plates were incubated overnight at 37°C, and colonies (which were visibly red due to RFP expression) were counted the following day. Bacterial burden was reported as colony-forming units (CFUs) per 20 µL of blood. Plates with colony numbers exceeding the reliable counting range were assigned a value of 2,000 CFUs.

### Quantifying RFP-expressing *E. coli* in brain sections

Sections were stained with DAPI to label nuclei and imaged for RFP-expressing *E. coli*. Fluorescence images were acquired as maximum intensity projections under identical imaging settings across experimental groups. For brain analyses, four anatomically matched territories from coronal sections of cortex were imaged and quantified per mouse. Quantification was performed using fixed threshold-based image analysis in Fiji, where the RFP signal was segmented to generate binary masks representing bacterial area. To define the corresponding cortical tissue area, DAPI images were thresholded and morphologically processed (including smoothing and hole filling) to generate continuous masks of the imaged cortical region. RFP signal was normalized to this DAPI-defined cortical area and reported as the ratio of RFP-positive area to DAPI-defined cortical area.

### bEnd.3 cell culture and maintenance

The mouse brain endothelial cell line bEnd.3 (ATCC CRL-2299) was cultured according to ATCC recommendations. Cells were maintained in Dulbecco’s Modified Eagle Medium (DMEM; high glucose, 4.5 g/L) supplemented with 10% fetal bovine serum (FBS). Cultures were kept at 37°C in a humidified 5% CO₂ incubator. Cells were passaged at ∼80–90% confluence using 0.05% trypsin–EDTA and reseeded at a split ratio of 1:4–1:6.

### CRISPR-Cas9 generation of bEnd.3 *Tlr4^KO^* cell lines

A bEnd.3-Cas9 monoclonal line was first established by transducing bEnd.3 cells with a lentivirus carrying *Streptococcus pyogenes* Cas9 (Addgene #52962) (Sanjana et al., 2014). Transduced cells were serially diluted to generate monoclonal populations, and Cas9 expression was validated by western blot. Using this Cas9-expressing line, *Tlr4* knockouts were generated by transduction with sgRNAs targeting exon 3 of *Tlr4* delivered with the sgTrack-mCherry lentivirus (Addgene #114013) (Hulton et al., 2020). Transduced cells were sorted by flow cytometry for mCherry expression, seeded in 96-well trays at one cell per well, and individual colonies were expanded. Genomic DNA was analyzed by PCR amplification across the mutated region in *Tlr4* exon 3, followed by subcloning of PCR products into a plasmid vector and sequencing of individual plasmids following colony PCR. A cloned bEnd.3 derivative cell line was considered a candidate *Tlr4^KO^* if none of its cloned PCR products showed the WT sequence and all cloned PCR products showed frameshift mutations in *Tlr4* exon 3. Functional loss of TLR4 was confirmed by quantifying NF-κB nuclear translocation after *E. coli* exposure compared to parental controls.

### bEnd.3 immunofluorescence

For all experiments involving responses to bacterial exposure, bEnd.3 cells were seeded onto glass coverslips at ∼80% confluency two days prior to the start of the experiment to allow them to form a confluent monolayer with tight junctions. After exposure to *E. coli* or GBS or mock exposure, cells were fixed in 1% paraformaldehyde in PBS for 15 min at room temperature, permeabilized in 0.1% Triton X-100/PBS, and blocked with 5% goat or donkey serum depending on the secondary antibody host species. Primary antibodies were applied at 1:400 dilution overnight at 4°C, followed by fluorophore-conjugated secondary antibodies. Coverslips were mounted in antifade medium and imaged by confocal microscopy.

### Antibodies for bEnd.3 cell immunostaining

Cultured bEnd.3 cells were immunostained with the following primary antibodies: mouse anti-Claudin-5 (Alexa Fluor™ 488, Thermo Fisher #352588), mouse anti-ZO-1 (Alexa Fluor™ 594, Thermo Fisher #339194), rabbit anti-NF-κB p65 (Cell Signaling Technology #8242S), rabbit anti-β-catenin (Cell Signaling Technology #8480), rat anti-PECAM-1 (Thermo Fisher #63-0311- 82), rabbit anti-GLUT1 (Thermo Fisher MA5-31960), mouse anti-EEA1 (Cell Signaling Technology #48453T), rabbit anti-LAMP2 (Cell Signaling Technology #34141T), rabbit anti- Rab7 (Cell Signaling Technology #9367T), rabbit anti-Rab11 (Cell Signaling Technology #5589T), rabbit anti-PDI (Cell Signaling Technology #3501T), and rabbit anti-RCAS1 (Cell Signaling Technology #12290T). Primary antibodies were diluted 1:400 in PBS with 1% Triton X-100 and applied overnight at 4 °C, followed by fluorophore-conjugated secondary antibodies (Thermo Fisher). Coverslips were mounted and imaged by confocal microscopy.

### bEnd.3 streptavidin leak assay

Barrier permeability was assayed using biotin-conjugated gelatin substrates as described by Dubrovskyi et al. (2013). Bovine gelatin (Sigma, G1393) was diluted to 1% in D-PBS, prewarmed to 37°C, and dissolved with stirring in a 70°C water bath. EZ-Link NHS-LC-Biotin (Thermo Fisher, Cat. No. 21336) dissolved in DMSO (5 mg/mL) was added to the gelatin at a final concentration of 0.5 mg/mL and reacted for 1 hour at room temperature with constant stirring. The solution was clarified by centrifugation (10,000 x g, 5 minutes), aliquoted, and stored at –20°C. For coating, biotinylated gelatin was thawed, diluted in 0.1 M bicarbonate buffer (pH 8.3) to 0.25 mg/mL, sterilized by 0.22 µm filtration, and applied to culture substrates (e.g., 50 µl per well of a 96-well plate, 2 ml per 35 mm dish). Adsorption was performed overnight at 4°C, followed by PBS washing. Unconjugated gelatin was used as a control.

Incorporation of biotin into gelatin was validated using the Pierce Biotin Quantitation Kit (Thermo Fisher, Cat. No. 28005) according to the manufacturer’s instructions. bEnd.3 monolayers were grown to confluence on coated coverslips, and control experiments included (i) gelatin without cells, (ii) biotinylated gelatin without cells, (iii) low-density cells on gelatin, and (iv) low- or high-density cells on biotinylated gelatin. To quantify the integrity of the bEnd.3 monolayer, streptavidin–FITC tracer (Thermo Fisher, #11-4317-87) at 2 µg/mL in DMEM was added to the cell culture medium for 5 min, followed by extensive PBS washing and fixation in 1% paraformaldehyde. Samples were immunostained for ZO-1, nuclei were visualized with DAPI, and the coverslips were mounted in antifade solution and imaged by confocal microscopy. Quantification of streptavidin signal was measured as area fraction per field of view. Each condition was repeated in three independent biological replicates, with multiple image fields analyzed per replicate. To ensure comparability across conditions, all images were collected using identical confocal acquisition settings, and exposure times and intensity scaling were held constant within each experiment.

### bEnd.3 and *E. coli* washout experiments

bEnd.3 cells were exposed to *E. coli* K1 (at a 5:1 ratio of *E.coli*:bEnd.3 cells) in serum-containing DMEM for 1 hour at 37°C, 5% CO₂. For washout, the medium was removed, and cells were washed extensively with pre-warmed PBS, then returned to fresh complete medium for a 3-hour recovery period. Parallel groups were fixed at 1 hour (no washout) or after the 3- hour washout. Cells were fixed in 1% paraformaldehyde, permeabilized, blocked, and immunostained (e.g., for Cldn5 and ZO-1) as described above, then mounted for confocal imaging. Acquisition settings (laser power, gain, offset, and channel order) were held constant across conditions within each experiment. Quantification of Claudin-5 cytoplasm/plasma membrane ratios followed the CellProfiler pipeline (Stirling et al., 2021), and results were analyzed as log_10_-transformed single-cell values with statistical significance assessed using the Wilcoxon rank-sum test. Three independent biological replicates were performed, with multiple fields analyzed per replicate. For inhibition of NF-κB signaling, ACHP (2-Amino-6-[2- (cyclopropylmethoxy)-6-hydroxyphenyl]-4-(4-piperidinyl)-3 pyridinecarbonitrile; Tocris #4547) was added to the cell culture medium at a final concentration of 1 µM.

### bEnd.3 tracer experiments

This procedure was performed as described (Stamatovic et al., 2006, 2017). Briefly, bEnd.3 monolayers on glass coverslips were preloaded with plasma-membrane–binding/uptake tracers prior to bacterial exposure: Cholera Toxin Subunit B, Alexa Fluor™ 647 conjugate (CTB-AF647; Thermo Fisher, C34778), Transferrin from human serum, Alexa Fluor™ 647 conjugate (Tf- AF647; Thermo Fisher, T23366), or Fixable Dextran, 10,000 MW, Alexa Fluor™ 647 (Dextran- AF647; Thermo Fisher, D22914). Tracers were added in ice-cold complete medium for 30 minutes at 4°C to allow accumulation at the plasma membrane. Excess tracer was removed by gentle washes with PBS, and cells were returned to tracer-free complete medium at 37°C. Cultures were then exposed to *E. coli* K1 (at a 5:1 ratio of *E.coli*:bEnd.3 cells) for the indicated intervals (e.g., 4 hours) or left unexposed as controls. Cells were fixed in 1% paraformaldehyde, permeabilized, blocked, and immunostained (e.g., for Cldn5 and ZO-1) as described above. Colocalization/overlap with Cldn5 was quantified in Fiji using fixed 100 x 100 µm ROIs and AND-masking of binarized channels (see “Percent overlap quantification”), with all acquisition parameters (laser power, gain, offset, channel order) held constant within each experiment. Three independent biological replicates were performed, with multiple fields analyzed per replicate.

### *E. coli* and GBS preparation and exposure of bEnd.3 cells

*Escherichia coli* K1 (RS218) and its RFP-expressing derivative (ampicillin-resistant) were cultured in LB broth and on LB agar plates with or without ampicillin, depending on the strain. *Streptococcus agalactiae* (GBS, ATCC 13813) was cultured in brain heart infusion (BHI) broth and on BHI agar plates. For tissue culture experiments, a single colony was grown overnight in the appropriate broth, and a 200 µl aliquot of the overnight culture was diluted into fresh medium and grown for 1 hour at 37°C. Bacterial density was estimated by optical density at 600 nm, and bacteria were added to cultured bEnd.3 cells (at a 5:1 ratio of bacteria to bEnd.3 cells for all experiments). Heat-killed bacteria were prepared in parallel by heating suspensions to 95°C for 30 min, and bacterial killing was confirmed by plating aliquots to check for the absence of growth. For bacterial exposure experiments, bEnd.3 cells were cultured in the absence of antibiotics in serum-containing medium, and bacteria remained in the medium for the duration of the experiment, except in designated washout assays.

### Image Analysis: Iba1 quantification

Immunostained Iba1 image quantification was performed as described above for quantifying RFP-expressing *E. coli* in brain sections, except that for Iba1 the superficial cortical territory beneath the leptomeninges was manually delineated in coronal sections based on the pattern of Cldn5 immunostaining. Iba1-positive signal was quantified using fixed threshold-based segmentation and normalized to the defined cortical area.

### Image Analysis: nuclear-to-cytoplasmic (Nuc/Cyto) intensity ratio

NF-κB and Ki-67 nuclear translocation was quantified using a custom CellProfiler pipeline (Stirling et al., 2021). Nuclei were segmented by DAPI, cytoplasmic regions were defined as the cell body excluding the nucleus, and mean intensities were measured in both compartments. Ratios of nuclear-to-cytoplasmic (Nuc/Cyto) intensity were computed per cell and log_10_-transformed for analysis. Output spreadsheets from CellProfiler were imported into R (v4.3) and annotated by parsing file names to assign genotype, clone, treatment, and timepoint metadata. Data from replicate experiments were combined, and box plots were generated to show distributions of single-cell values. Statistical comparisons between conditions were performed using the Wilcoxon rank-sum test. For validation, replicate-level plots were generated, outliers were defined by interquartile range (1.5 x IQR) and excluded from simplified representative plots, and a subset of clones and timepoints (e.g., 0, 1, and 3 hours) were displayed for final figures.

### Image Analysis: cytoplasmic-to-plasma membrane (Cyto/PM) intensity ratio

Claudin-5 redistribution between the plasma membrane and cytoplasm was quantified in CellProfiler (Stirling et al., 2021) using a custom pipeline. Plasma membrane regions were segmented using ZO-1 as a boundary, while cytoplasmic regions were defined as the cell body excluding the nucleus and plasma membrane masks. Integrated mean intensities of Claudin-5 were measured in both regions, and cytoplasmic-to-plasma membrane (Cyto/PM) ratios were calculated for each cell and log₁₀-transformed for analysis. Output spreadsheets from CellProfiler were imported into R (v4.3) for downstream analysis. File names were parsed to assign genotype, treatment, timepoint, and replicate metadata, and data from replicate experiments were combined. Box plots were generated to display distributions of single-cell values, with statistical comparisons performed using the Wilcoxon rank-sum test. For representative figures, a single replicate per group was shown, and outliers were removed based on the interquartile range (1.5 x IQR).

### Image Analysis: percent overlap quantification (ImageJ)

Colocalization between Cldn5 and trafficking/endothelial markers (e.g., EEA1, Rab7, Rab11, LAMP2, ZO-1) or tracers was quantified in Fiji (Schindelin et al., 2012) using a scripted workflow. For each field, maximum-intensity–projected channels were processed as follows: images were Gaussian-blurred (σ = 2), converted to 8-bit, and binarized using fixed thresholds chosen per channel and experiment. Binary masks were generated with “BlackBackground” enabled and saved for Cldn5 and the comparison marker. Overlap masks were created with Image Calculator (logical AND) between the Cldn5 mask and the marker mask (and, when needed, between Cldn5 and ZO-1). An ROI set comprising nine 100 x 100 µm squares was loaded via the Region of Interest (ROI) Manager and applied uniformly to all images. For each ROI, we recorded raw mean intensities (non-binary images) and, on the binary masks, measured area and mean intensity. “Signal area” was computed as (ROI area x mask mean)/255 for Cldn5 and for the overlap image, providing an area-weighted signal metric. Percent overlap with Cldn5 was then calculated per ROI as 100 x (overlap signal area ÷ Cldn5 signal area). Identical steps were repeated for each endosomal marker or tracer by substituting the corresponding channel and threshold. Outputs were written to CSV and pooled across images/replicates for plotting; statistical comparisons were performed using Wilcoxon rank-sum tests as described in the Statistics section. All acquisition and analysis parameters (thresholds, ROIs, and export settings) were held constant within a given experiment.

### Image Analysis: statistics

All image quantifications were performed on raw data exported directly from Fiji or CellProfiler without manual exclusion of individual cells or ROIs, unless otherwise noted for representative plots. For Cyto/PM and Nuc/Cyto ratios, values were log₁₀-transformed prior to statistical testing. Percent overlap measurements were expressed as the fraction of Cldn5-positive signal area colocalizing with the comparison marker within fixed 100 x 100 µm ROIs. For extravascular tracer (Sulfo-NHS-biotin) and ICAM-1 analyses, fluorescence intensity or area fractions were normalized to the ROI area or expressed as fold change relative to WT uninfected controls. For all datasets, each dot represents a single measurement (one cell *in vitro* or one image field *in vivo*), and biological replicates correspond to independent mice or independent cell culture experiments. Statistical comparisons between two groups were performed using the two-sided Wilcoxon rank-sum test (Mann–Whitney U). Where multiple groups were compared, pairwise Wilcoxon rank-sum tests were performed, and p-values are reported in figure panels. Box plots display the median and interquartile range, with all individual data points overlaid. All statistical analyses were conducted in R (v4.3) using ggplot2 and ggpubr packages, with significance set at p < 0.05.

### snRNA-seq libraries and sequencing

For single-nucleus RNA sequencing, leptomeninges were collected as previously described (Wang et al., 2023). In brief, P6 mice were euthanized 24 hours after infection, brains were rapidly removed, and the leptomeninges were dissected into ice-cold Hibernate-A medium. The leptomeninges were frozen at -80°C.

For the PIP-seq protocol, single nucleus (sn)RNA-seq libraries were prepared using the PIP- seq T20 3’ Single Cell RNA Kit v4.0 PLUS (Fluent BioSciences; Clark et al., 2023). Frozen tissue was suspended in homogenization buffer (0.25 M sucrose, 25 mM KCl, 5 mM MgCl_2_, 20 mM Tricine-KOH,pH 7.8) supplemented with 1 mM DTT, 0.15 mM spermine, 0.5 mM spermidine, EDTA free protease inhibitor (Roche 11836 170 001), 0.5% IGEPAL-630, and 40 U/mL Protector RNase Inhibitor (Sigma, 03335402001). Samples were homogenized in a 2 mL Dounce homogenizer, using 15 strokes with a loose-fitting pestle followed by 30 strokes with a tight-fitting pestle. The sample was filtered through a 10 μm filter (CellTrix, Sysmex, 04-004- 2324), and nuclei were pelleted in Low Retention 1.5 mL microcentrifuge tubes (Thermo Scientific, 3451) for 5 min at 500 x g at 4 °C. Nuclei were washed twice with Nuclei Suspension Buffer (supplied in the Fluent BioSciences kit) supplemented with 1% BSA and 40 U/mL Protector RNase Inhibitor. Nuclei were counted, and ∼40,000 nuclei (for the T20 kit) or ∼20,000 nuclei (for the T10 kit) were used for library production following the Fluent BioSiences protocol. The resulting snRNA-seq libraries were sequenced on an Illumina NovaSeq X Plus sequencer (Supplementary file 1).

For the 10X genomics protocol, snRNA-seq libraries were constructed from suspended nuclei (prepared as described above) using the 10x Genomics Chromium single-cell 3’ v3 kit following the manufacturer’s protocol. Libraries were sequenced on an Illumina NovaSeq 6000.

### Analysis of snRNA-seq data

Reads were aligned with the PIPseeker program (Fluent BioSiences, version 3.3.0) using a STAR index based on the GRCm38 mouse genome with addition of Cre sequences (STAR version 2.7.9a). The CellBender program was used to detect empty droplets and remove background (version 0.3.2; Fleming et al., 2023). The SOLO program was used to remove doublets (scVI-tools 1.2.2-post2) (Bernstein et al., 2020). Manual curation was used to further remove potentially mixed (doublet) nuclei. Clusters representing various brain cells were removed from the final data set. The data were normalized using a regularized negative binomial regression algorithm implemented with the SCTransform function (Hafemeister and Satija, 2019). UMAP dimensional reduction was performed using the R uwot package (https://github.com/jlmelville/uwot) integrated into the Seurat R package (Melville, 2022). Data for the various scatter plots were extracted using the Seurat AverageExpression function, and differential gene expression was analyzed using the Seurat FindMarkers function. The Wilcoxon rank sum test was used to calculate p-values. The p-values were adjusted with a Bonferroni correction using all genes in the dataset. Data exploration, analysis, and plotting were performed using RStudio (RStudio Team, 2020), the tidyverse collection of R packages (Wickham, 2017), and ggplot2 (Wickham, 2009).

Dotplots were generated with the default settings, including default normalization. For Figure 1E-H and Figure 1 – figure supplement 5, differential expression between infected and control samples was calculated using Seurat FindMarkers function using only the WT data. The results were filtered for statistically significant difference per cell type. Differentially expressed genes from the indicated Hallmark gene sets are presented in the dot plots for individual cell clusters.

### GSEA analysis

For Gene Set Enrichment Analysis (GSEA; Subramanian et al., 2005), genes were ranked by the fold expression change between control and mutant datasets. The ranked gene list was used to detect enriched gene sets within the Broad Institute Hallmark Gene Sets using the fgsea R package (https://github.com/ctlab/fgsea; Korotkevich et al., 2019).

### RNA-seq on WT and *Tlr4^KO^* bEnd.3 cells: libraries and sequencing

bEnd.3 mouse brain endothelial cells were cultured in 10-cm tissue culture dishes and grown to confluency under standard conditions (37°C, 5% CO₂) and either exposed or not exposed to *E. coli* for 3 hours. Total RNA was isolated using the Zymo Research RNA purification kit according to the manufacturer’s instructions. Briefly, culture medium was aspirated, cells were rinsed once with ice-cold PBS, and 600 µL of the provided lysis buffer with β-mercapto-ethanol was added directly to the dish. The lysate was collected by scraping and transferred to a microcentrifuge tube, briefly mixed, and applied to the Zymo spin column. After binding, the column was washed with the supplied wash buffers, including an on-column DNase I treatment step to remove contaminating genomic DNA, and RNA was eluted in 50 µL of DNase/RNase-free water. RNA concentration and purity were assessed by spectrophotometry (A₂₆₀/A₂₈₀), and samples were stored at −80°C. Library construction was performed with the Illumina mRNA stranded kit following the manufacturer’s protocol, and sequencing was performed on an Illumina NovaSeqX with a read length of 300 nucleotides.

### RNA-seq on WT and *Tlr4^KO^* bEnd.3 cells: RNA-seq processing and normalization

Gene-level count matrices from bEnd.3 cells were analyzed in R (v4.5) with Bioconductor (v3.21). The study included two genotypes (WT and *Tlr4^KO^*) under two conditions (control and exposed to *E. coli* for 3 hours), with biological replication per group (WT: 3 control samples and 4 exposed to *E. coli* samples; *Tlr4^KO^*: 3 control samples and 4 exposed to *E. coli* samples). Counts were normalized using DESeq2’s median-of-ratios size-factor method to control for library size/composition effects. A variance-stabilizing transform (VST) was calculated for quality control and visualization only (e.g., PCA, sample-distance heatmaps). All samples were retained for analysis after QC review.

### RNA-seq on WT and *Tlr4^KO^* bEnd.3 cells: Differential expression

To quantify infection responses within each genotype, DESeq2’s negative-binomial generalized linear model was fit separately to WT and in *Tlr4^KO^* data and then control vs. exposed to *E. coli* data were compared. The reported log_2_ fold changes are the control vs. exposed to *E. coli* estimates from these genotype-specific analyses. P-values were adjusted for multiple testing with the Benjamini–Hochberg procedure. False-discovery rates (FDR or q-values) are cited throughout. Gene-level WT vs. *Tlr4^KO^* effect comparisons were visualized by plotting the two genotype-specific fold changes against one another.

### RNA-seq on WT and *Tlr4^KO^* bEnd.3 cells: Pathway activity scoring and statistics

Pathway analyses used the MSigDB Hallmark collection for mouse, retrieved via msigdbr. Single-sample gene-set enrichment (ssGSEA) was performed with GSVA to obtain an activity score for each pathway in each sample. Scores were computed from log_2_-scaled normalized counts (log_2_ (normalized counts + 1)), using only genes present in both the RNA-seq data and the gene set. For statistical inference, we modeled pathway scores with a linear model that included genotype, infection status, and their interaction. From this model we reported three quantities per pathway: the *E. coli* exposure effect in WT, the *E. coli* exposure effect in *Tlr4^KO^*, and the genotype x exposure interaction (difference-in-response between genotypes). P-values were adjusted across pathways with the Benjamini–Hochberg procedure. Adjusted values are reported as q-values (FDR).

### RNA-seq on WT and *Tlr4^KO^* bEnd.3 cells: Visualization

Heatmaps were drawn from VST values after centering each gene by its mean (raw z-scores). Within each Hallmark pathway, genes were ranked by the magnitude of the *E. coli* exposure effect (difference of condition means within each genotype), and the top genes were displayed to show the pattern and directionality across groups. Boxplots summarize ssGSEA scores by group. Per-facet annotations show FDRs for WT and *Tlr4^KO^*infection effects and the interaction term. Figure 7C plots the pathway-level infection responses in WT (x-axis) versus *Tlr4^KO^*(y- axis), with point size reflecting within-genotype significance and point color indicating interaction FDR. All figures were exported as vector PDFs.

### RNA-seq on WT and *Tlr4^KO^* bEnd.3 cells: Software

Analyses used DESeq2, limma (for pathway-score modeling), SummarizedExperiment, GSVA, msigdbr, pheatmap, ggplot2, and ggrepel within the R/Bioconductor ecosystem.

## Data availability

All RNA sequencing data were deposited in GEO (accession number GSE319556).

## Acknowledgements.

Supported by the Howard Hughes Medical Institute. The authors thank Timothy Phelps (Department of Art as Applied to Medicine, Johns Hopkins Medical School) for the drawing in Figure 1A, the members of the Johns Hopkins Single Cell and Transcriptomics Core Facility for preparing and sequencing RNAseq libraries from bEnd.3 cells, Ying Wang for assistance dissecting the leptomeninges, Britta Engelhardt (Theodor Kocher Institute) for advice, and Zhongming Li for helpful comments on the manuscript.

## Supplemental Files

**Supplementary file 1. Overview of snRNA-seq on leptomeninges.**

**Supplementary file 2. Numbers of mice and cells used in various experiments.**

## Supplemental Figure Legends

**Figure 1 – figure supplement 1.**
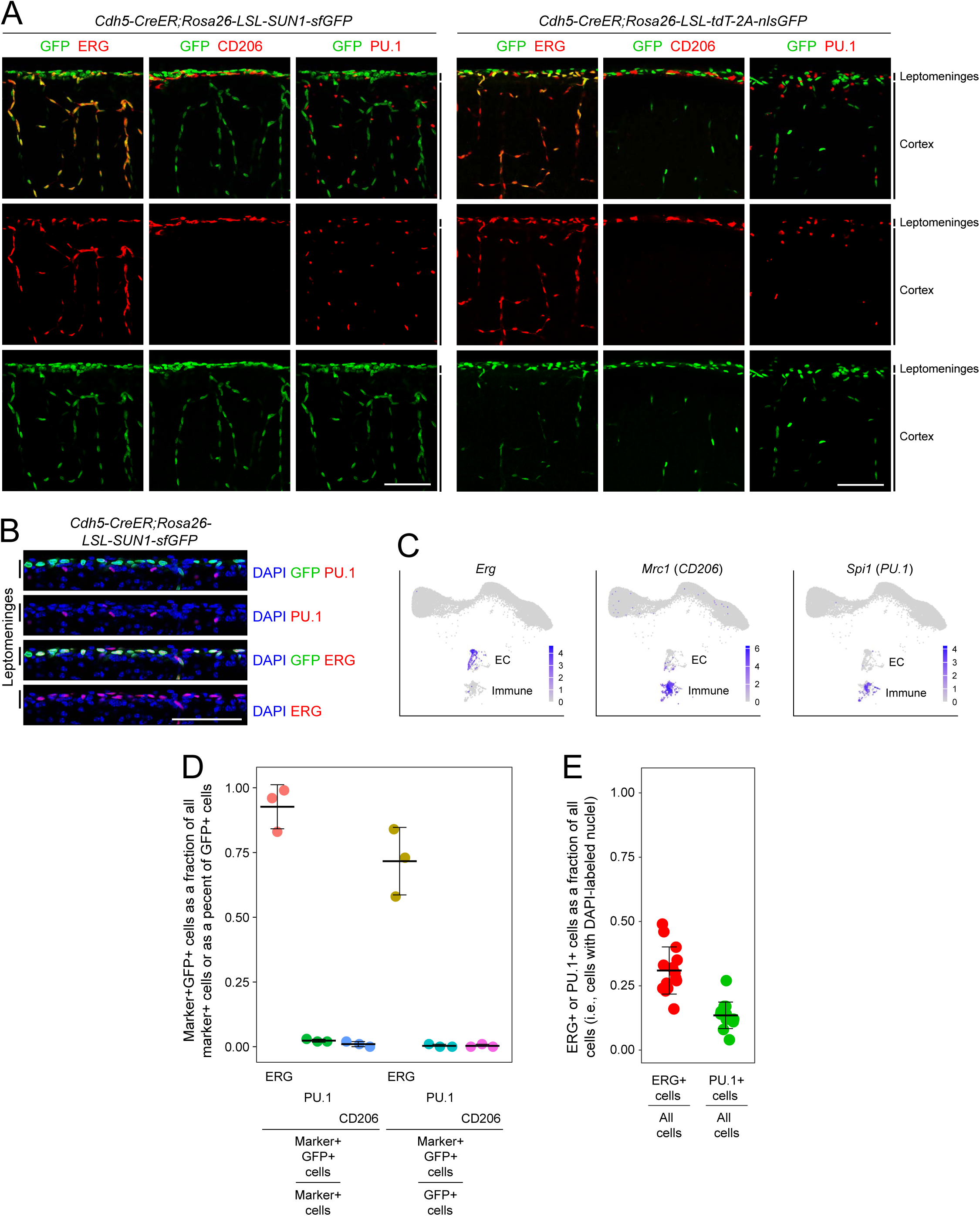
Specificity of *Cdh5-Cre*ER in leptomeninges. (A) *Cdh5Cre*-mediated recombination of the *loxP-stop-loxP* (*LSL*) reporters *Rosa26-LSL-SUN1- sfGFP* (left) and *Rosa26-LSL-tdT-2A-nlsGFP* (right). Immunostaining for markers ERG, CD206, and PU.1 was performed on coronal sections of P6 brains following 4HT administration at P2. Nuclear-localized GFP is observed in nearly all ECs, as visualized with ERG immunostaining, but not in myeloid cells, as visualized with PU.1 and CD206 immunostaining. The region corresponding to the leptomeninges is labeled on the right. Scale bars, 100 µm. (B) Immunostaining for markers ERG and PU.1 was performed on coronal sections of P6 brains from *Cdh5Cre*;*Rosa26-LSL-SUN1-sfGFP* mice as described for (A). The leptomeninges are demarcated by the black bars at left. Scale bar, 100 µm. (C) WT uninfected leptomeninges UMAPs showing the expression of the genes encoding the markers used in (A) and (B): *Erg* (ECs), *Mrc1* (CD206; myeloid cells); *Spi1* (PU.1; myeloid cells). (D) Quantification of the co-localization of nuclear GFP from *Rosa26-LSL-tdT-2A-nlsGFP* with cell-type-specific markers in the leptomeninges, based on images like the ones in (A). The left set of three data points show the fraction of marker+ cells that express GFP. The right set of three data points show the fraction of GFP+ cells that express each of the three cell-type specific markers. Approximately 75% of GFP+ cells are ECs and approximately 0% are myeloid cells. Therefore, the remaining 25% of GFP+ cells correspond to some combination of other leptomeningeal cell types. Each data point represents one mouse. A mean of 158 cells was scored per datapoint. Bars, mean +/- S.D. (E) Fraction of all leptomeningeal cells, quantified by counting DAPI-stained nuclei, that have ERG+ nuclei (left) or PU.1+ nuclei (right) based on images like the ones in (B). Three mice were analyzed. Five images were analyzed per mouse, and each data point represents a single image. A mean of 60 cells was analyzed per image. Bars, mean +/- S.D.

**Figure 1 – figure supplement 2.**
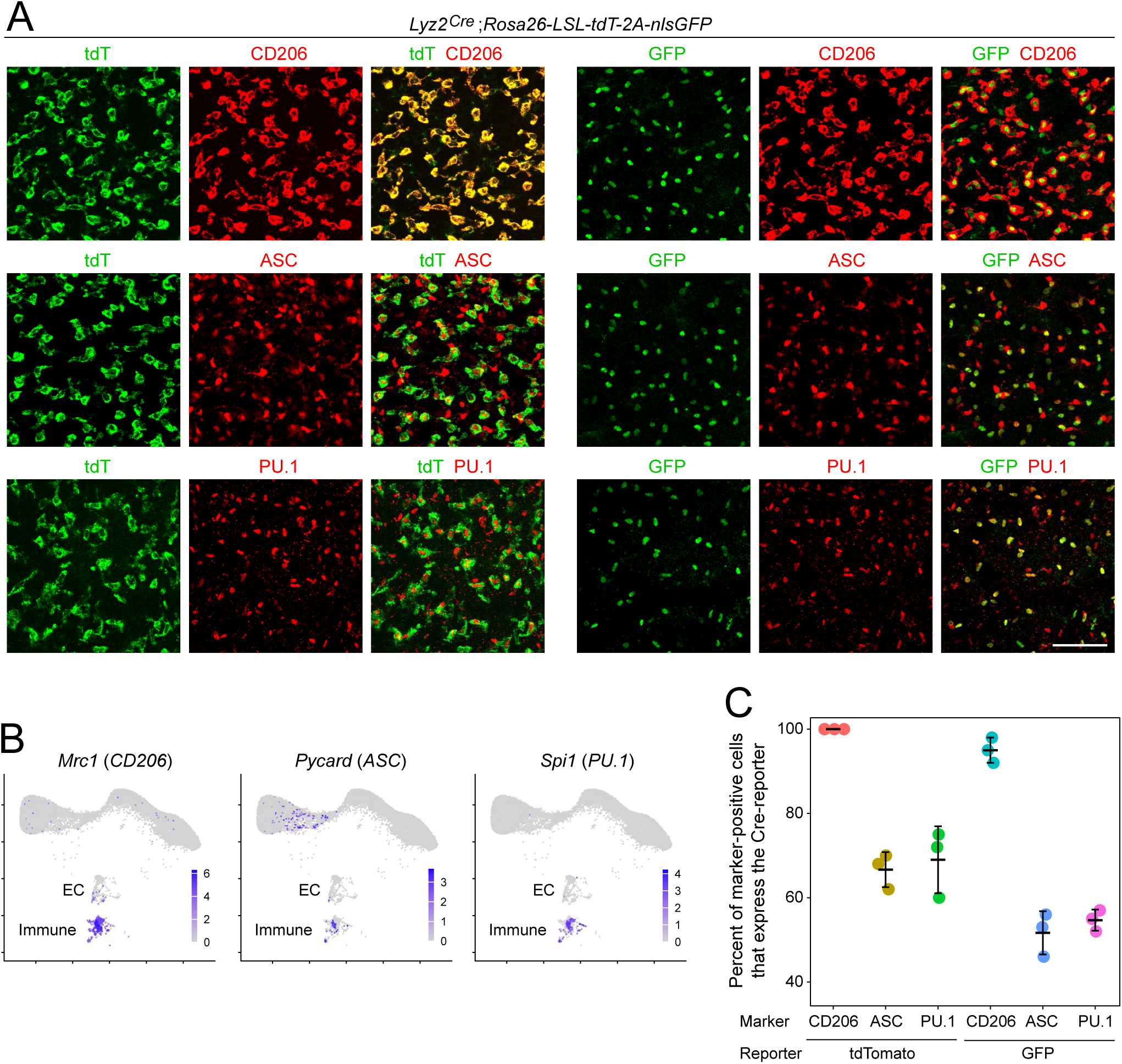
Specificity of *Lyz2^Cre^* in leptomeninges. (A) *Lyz2Cre*-mediated recombination of the *loxP-stop-loxP* (*LSL*) reporter *Rosa26-LSL-tdT-2A- nlsGFP*, with the tdTomato reporter (false-colored green) shown in the left set of nine panels and the nuclear-localized GFP reporter (green) shown in the right set of nine panels. Immunostaining for markers CD206, PU.1, and ASC was performed on P6 leptomeninges+cortex whole mounts, as described for Figure 1B. The leptomeninges region is shown. Scale bar, 200 µm. (B) WT uninfected leptomeninges UMAPs showing the expression of the genes encoding the markers used in (A): *Mrc1* (CD206; myeloid cells), Pycard (ASC; myeloid plus arachnoid barrier cells), and *Spi1* (PU.1; myeloid cells). (C) Quantification of co-localization data, as shown in (A). Each data point represents one mouse. A mean of 136 cells was scored per datapoint. The Cre reporters, tdTomato and GFP, are expressed in 90-100% of CD206+ myeloid cells, in 40-70% of ASC+ cells, and in 50-80% of PU.1+ cells.

**Figure 1 – figure supplement 3.**
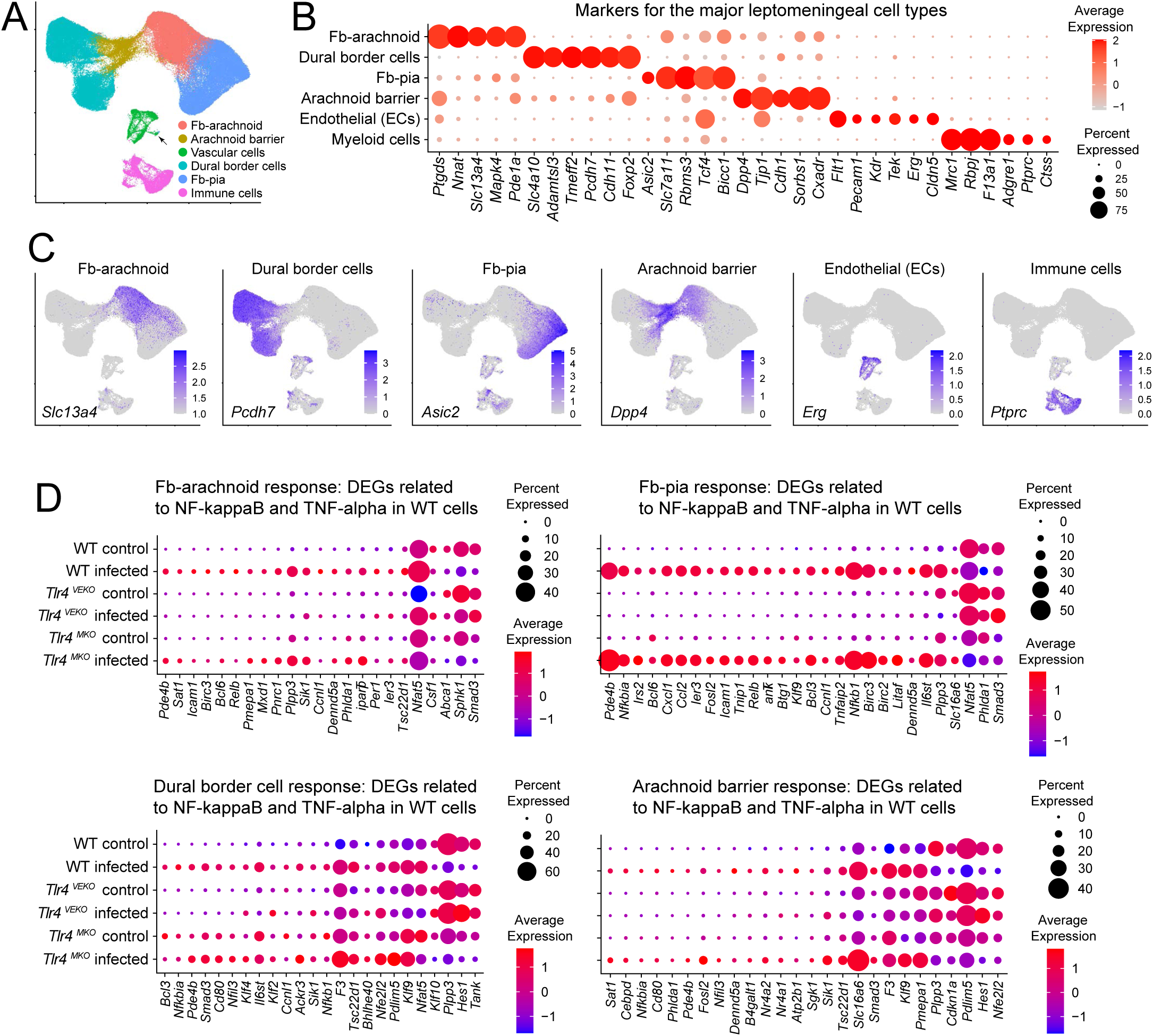
snRNA-seq cell cluster assignments and analysis of differentially expressed genes related to NK-κB and TNF-α responses. (A) UMAP plot of leptomeninges snRNA-seq data from WT, *Tlr4^VEKO^*, and *Tlr4^MKO^* mice at P6, either uninfected control or with *E. coli* infection, with cell clusters identified. N=126,235 nuclei. Fb, fibroblast. (B) snRNA-seq transcript abundances among the six major leptomeninges cell clusters for a set of 33 genes, for which the transcript abundances distinguish these clusters. We note that the dural border cell cluster was originally referred to as “fibroblasts dura3” in Wang et al. (2023). (C) UMAP plots of leptomeninges snRNA-seq showing the abundances of transcripts that are highly enriched in each of the six major leptomeningeal cell clusters. (D) snRNA-seq from four leptomeningeal cell clusters showing differentially expressed genes related to NK-κB and TNF-α responses in control vs. *E. coli*-infected mice of the indicated genotypes at P6. The corresponding dot plots for ECs and myeloid cells are shown in Figure 1H.

**Figure 1 – figure supplement 4.**
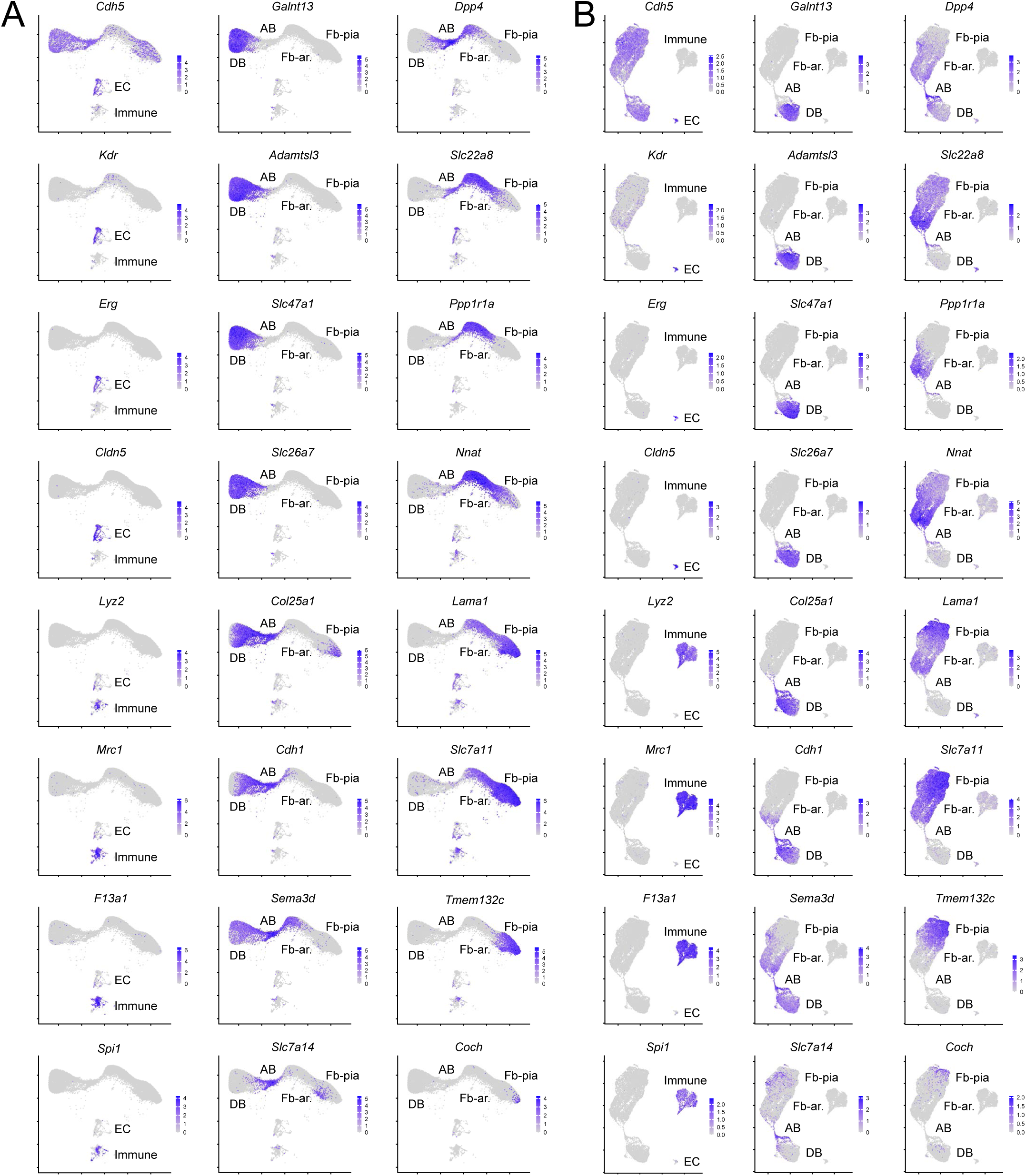
Comparison of WT P6 leptomeninges snRNA-seq using the PIP-seq and 10X Genomics platforms. (A) WT P6 leptomeninges snRNA-seq determined with the PIP-seq platform. Expression of 24 genes is illustrated to highlight: *Cdh5* (upper left), EC and immune cell genes (left column), and, the four principal cell clusters (right two columns; from left to right within each UMAP these are Dural border cells (DB), Arachnoid barrier cells (AB), Fibroblast arachnoid (Fb-ar.), and Fibroblast-pia (Fb-pia). (B) As for (A), except that the WT P6 leptomeninges snRNA-seq was determined with the 10X Genomics platform. Note that the yields of immune cells and ECs differ between the two datasets.

**Figure 1 – figure supplement 5.**
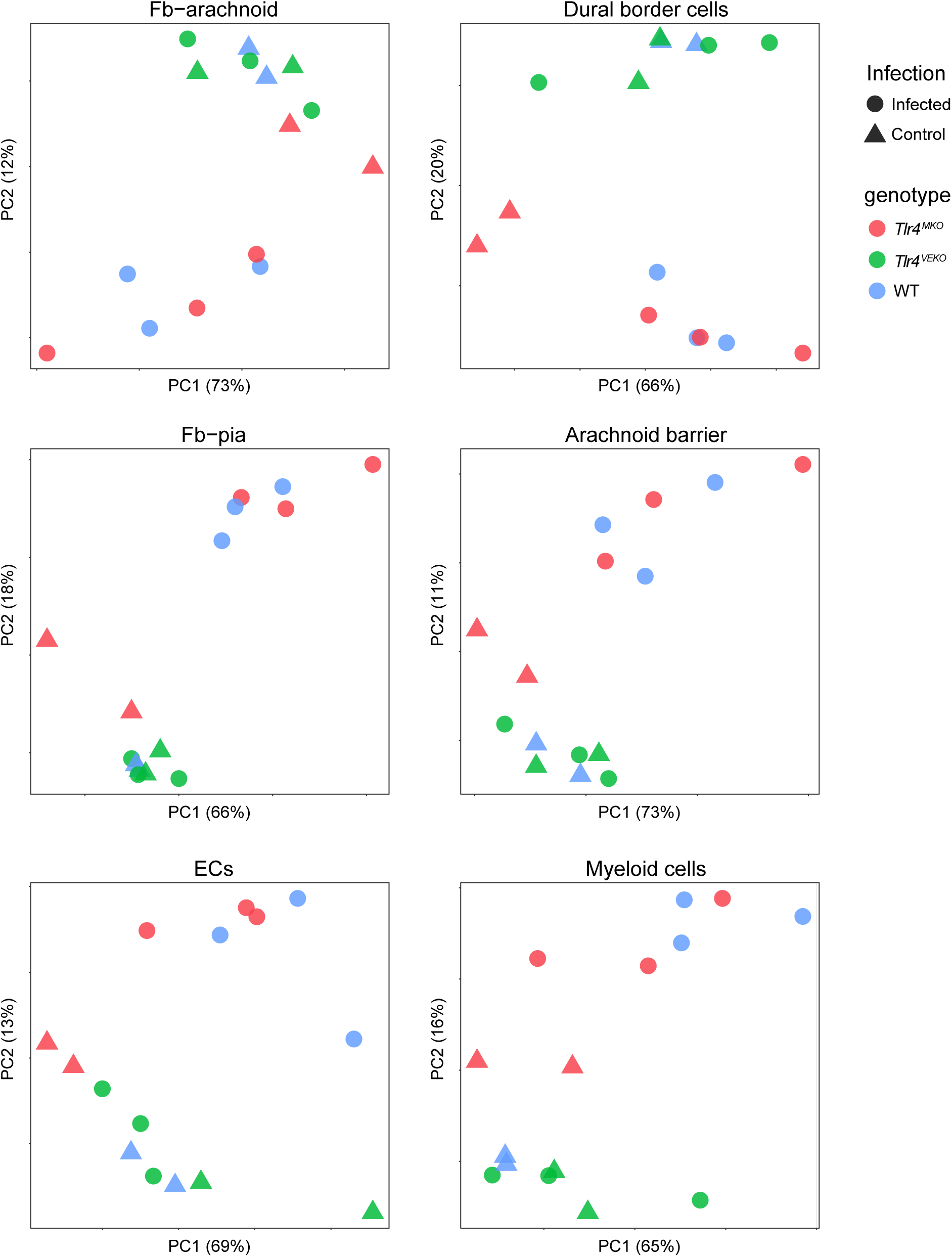
Principal component analysis of snRNA-seq-derived transcriptomes for each of the six major leptomeningeal cell clusters divided by genotype and infection status. The two experimental conditions (control vs. infected) and the three genotypes (WT, *Tlr4^VEKO^*, and *Tlr4^MKO^*) are indicated by the colors and symbols in the upper right. The datapoints represent the same leptomeningeal snRNA-seq datasets analyzed in Figure 1C-H, from control vs. *E. coli*-infected mice at P6.

**Figure 1 – figure supplement 6.**
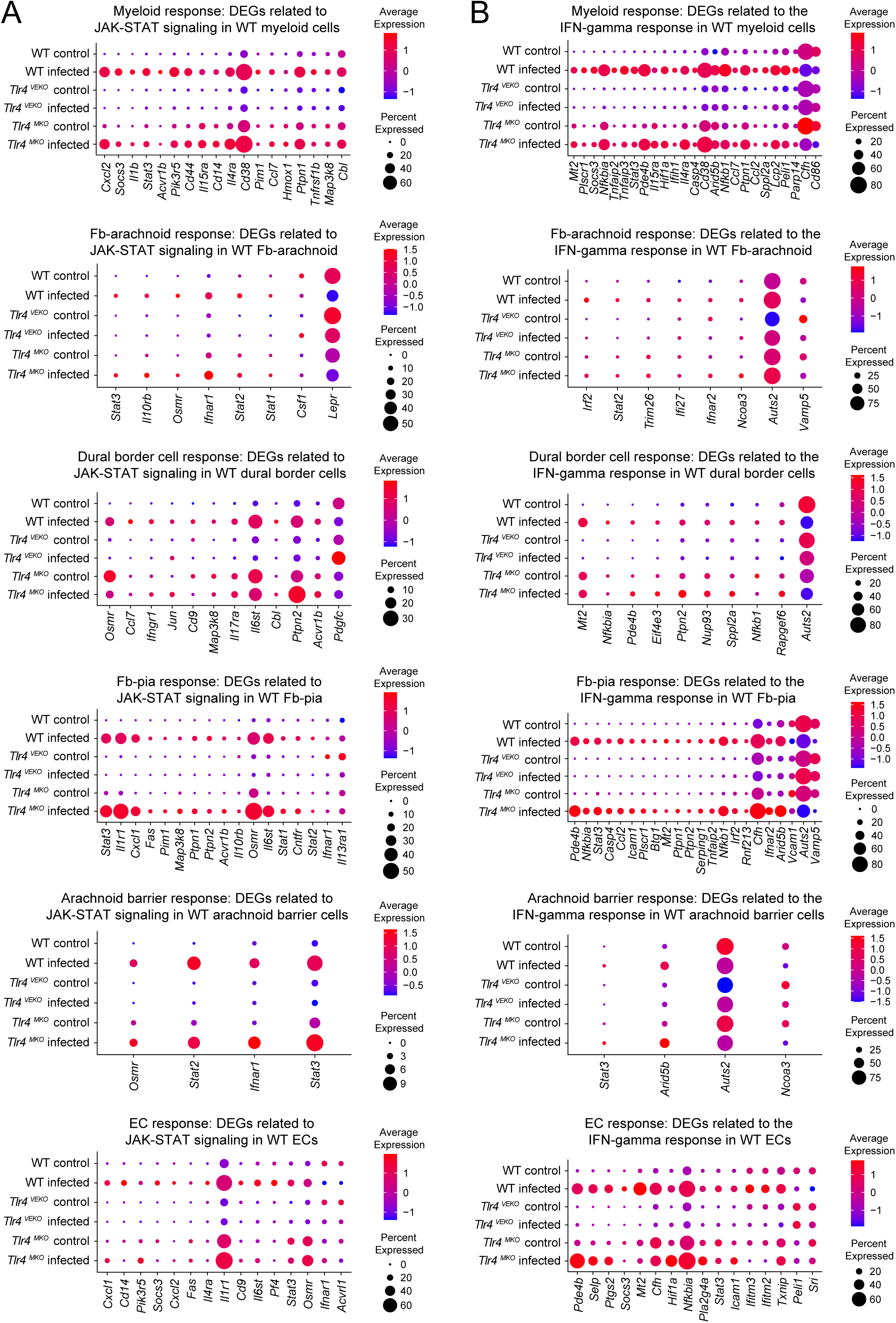
Transcriptome changes by cell cluster for the JAK-STAT and IFN-ɣ pathways. (A and B) snRNA-seq from each of the major leptomeninges cell clusters showing differentially expressed genes that are related to the JAK-STAT pathway (A) or the IFN-ɣ pathway (B). Dot plots are based on the same leptomeningeal snRNA-seq datasets analyzed in Figure 1C-H, from control vs. *E. coli*-infected mice at P6.

**Figure 1 – figure supplement 7.**
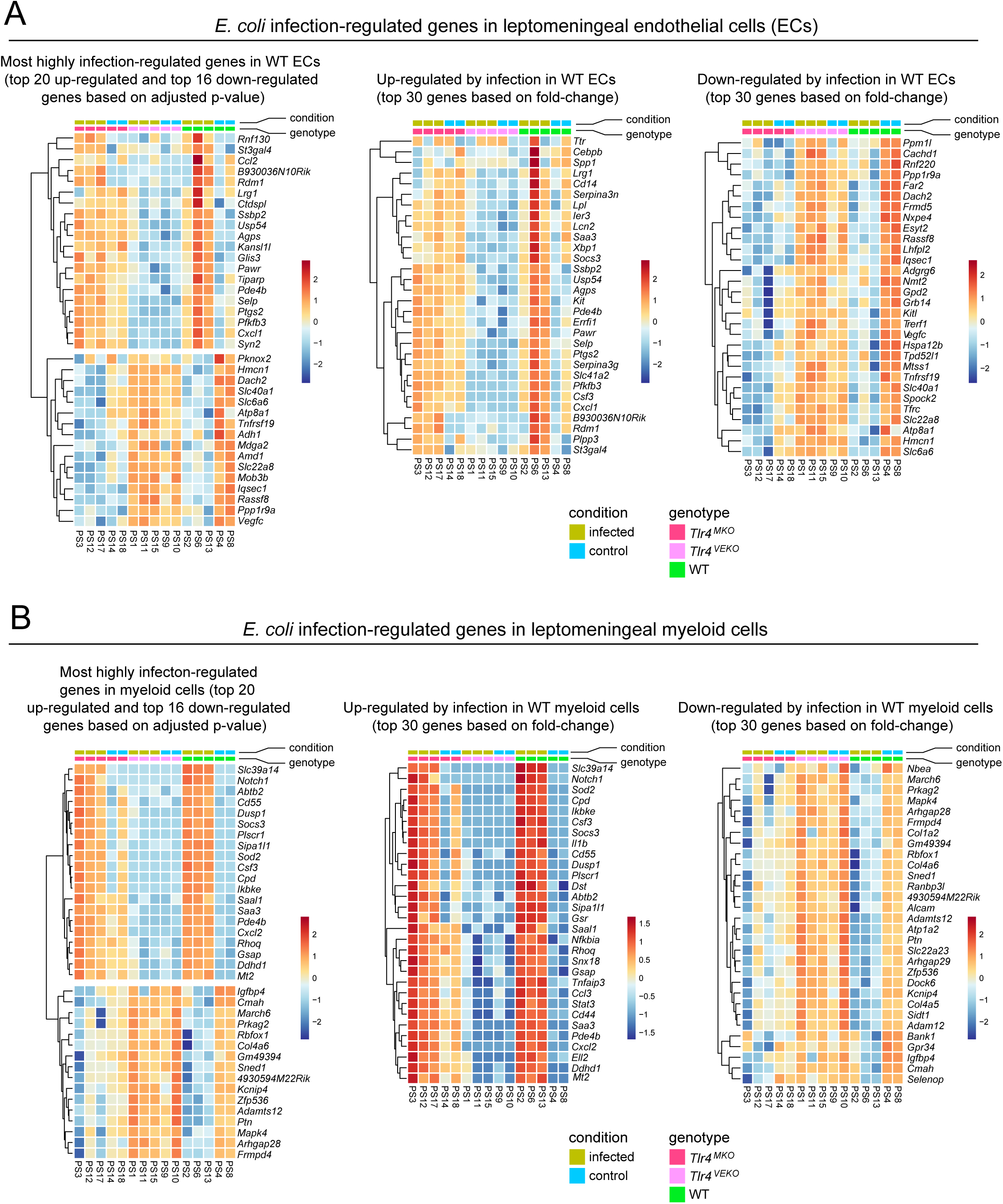
Genes regulated by *E. coli* infection in leptomeningeal endothelial cells and myeloid cells analyzed at the level of individual mice and individual genes. (A) Genes regulated by *E. coli* infection in leptomeningeal ECs. Left panel, most differentially regulated genes based on adjusted p-value. Right two panels, most differentially regulated genes based on fold change. (B) Genes regulated by *E. coli* infection in leptomeningeal myeloid cells. Left panel, most differentially regulated genes based on adjusted p-value. Right two panels, most differentially regulated genes based on fold change. The data are derived from the same leptomeningeal snRNA-seq datasets analyzed in Figure 1C-H, from control vs. *E. coli*-infected mice at P6.

**Figure 2 – figure supplement 1.**
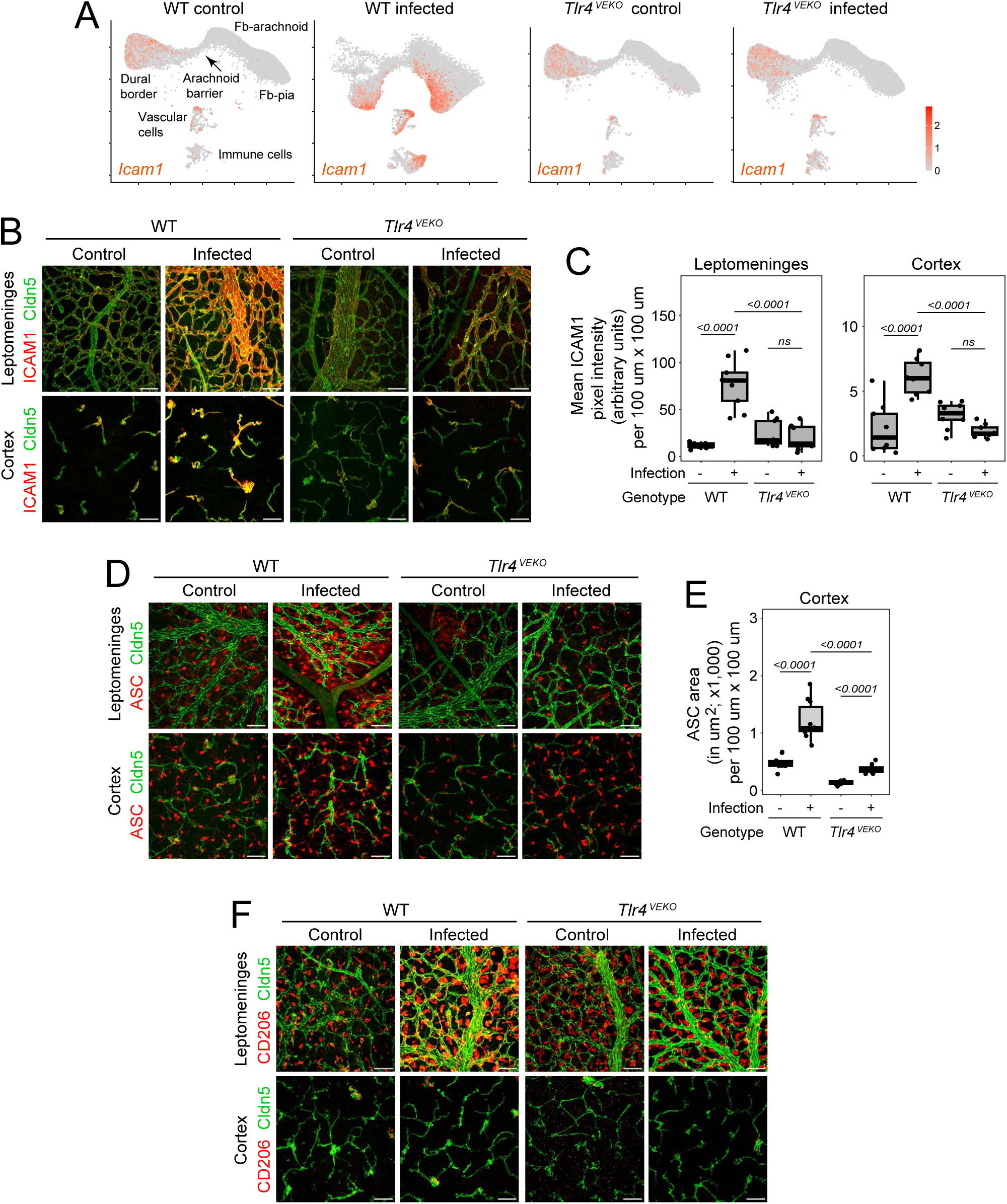
Infection responses in the leptomeninges and adjacent cerebral cortex in WT and *Tlr4^VEKO^* mice. (A) UMAP plots of leptomeninges snRNA-seq data from WT and *Tlr4^VEKO^* mice at P6, either uninfected control or with *E. coli* infection, showing *Icam1* transcripts. (B) Whole mount cortical surface, with attached leptomeninges, from P6 WT and *Tlr4^VEKO^* mice, control or *E. coli*-infected, immunostained for Cldn5 and ICAM1. Upper panels, stacked Z- planes at the level of the leptomeninges; lower panels, stacked Z-planes at the level of the adjacent cerebral cortex. Scale bars, 20 µm. (C) Quantification of ICAM1 immunostaining in the leptomeninges and adjacent cerebral cortex, as shown in (B). Note the different scales for the vertical axis in the two plots. (D) As in (B), except immunostained for Cldn5 and ASC. These are the same four leptomeninges+cortex flat mounts as shown in Figure 2D: the upper row of four images shown here reproduces the upper row of four images shown in Figure 2D, and the lower row of four images shown here correspond to the regions in the top row but were captured at a greater depth to visualize the underlying cortex. (E) Quantification of ASC immunostaining in the cerebral cortex, as shown in the lower row of images in (D). (F) As in (B), except immunostained for Cldn5 and CD206. These are the same four leptomeninges+cortex flat mounts as shown in Figure 2D: the upper row of four images shown here reproduces the lower row of four images shown in Figure 2D, and the lower row of four images shown here correspond to the regions in the top row but were captured at greater depth to visualize the underlying cortex. These data are not quantified because CD206 cells were extremely rare in the cortex in all samples.

**Figure 2 – figure supplement 2.**
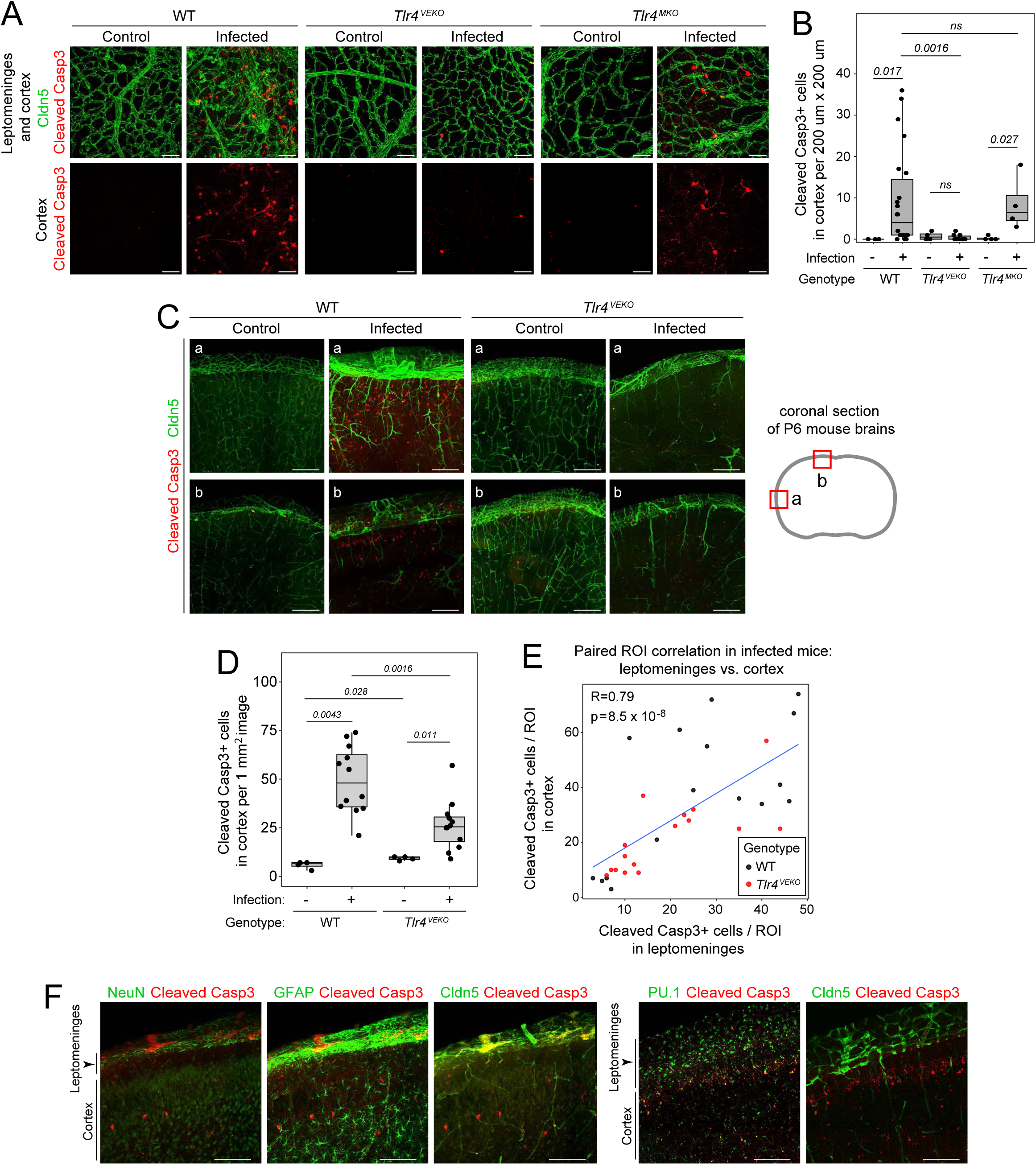
Cleaved Caspase-3+ cells accumulate in the cortex in *E. coli*-infected WT and *Tlr4^MKO^* mice but not in *Tlr4^VEKO^* P6 mice. (A) Flat mount images showing the meninges and adjacent cortex for the indicated genotypes and conditions. Scale bar, 50 µm. (B) Quantification of cleaved Caspase-3+ cells from flat mount images like the ones shown in (A). The region of cortex analyzed is adjacent to the leptomeninges. (C) Coronal sections showing meninges (upper) and cortex (lower). For quantification, two regions were imaged per hemisphere, as shown in the schematic on the right and labeled ‘a’ and ‘b’. Examples of ‘a’ and ‘b’ images are shown at left. Scale bar, 200 µm. (D) The number of cleaved Caspase-3+ cells in leptomeninges and cortex were quantified from 1 mm x 1 mm images like those shown in (C). The different planes of sectioning and the different cortical depths in the images quantified in (B) and (D) account for the different numbers of cleaved caspase3+ cells. (E) Correlation between cleaved Caspase-3+ cells in the leptomeninges and cortex for individual coronal images as shown in (C). (F) Cleaved Caspase-3+ cells in cortex are NeuN-, GFAP-, Cldn5-, and PU.1+, implying a myeloid identity. Scale bar, 100 µm.

**Figure 2 – figure supplement 3.**
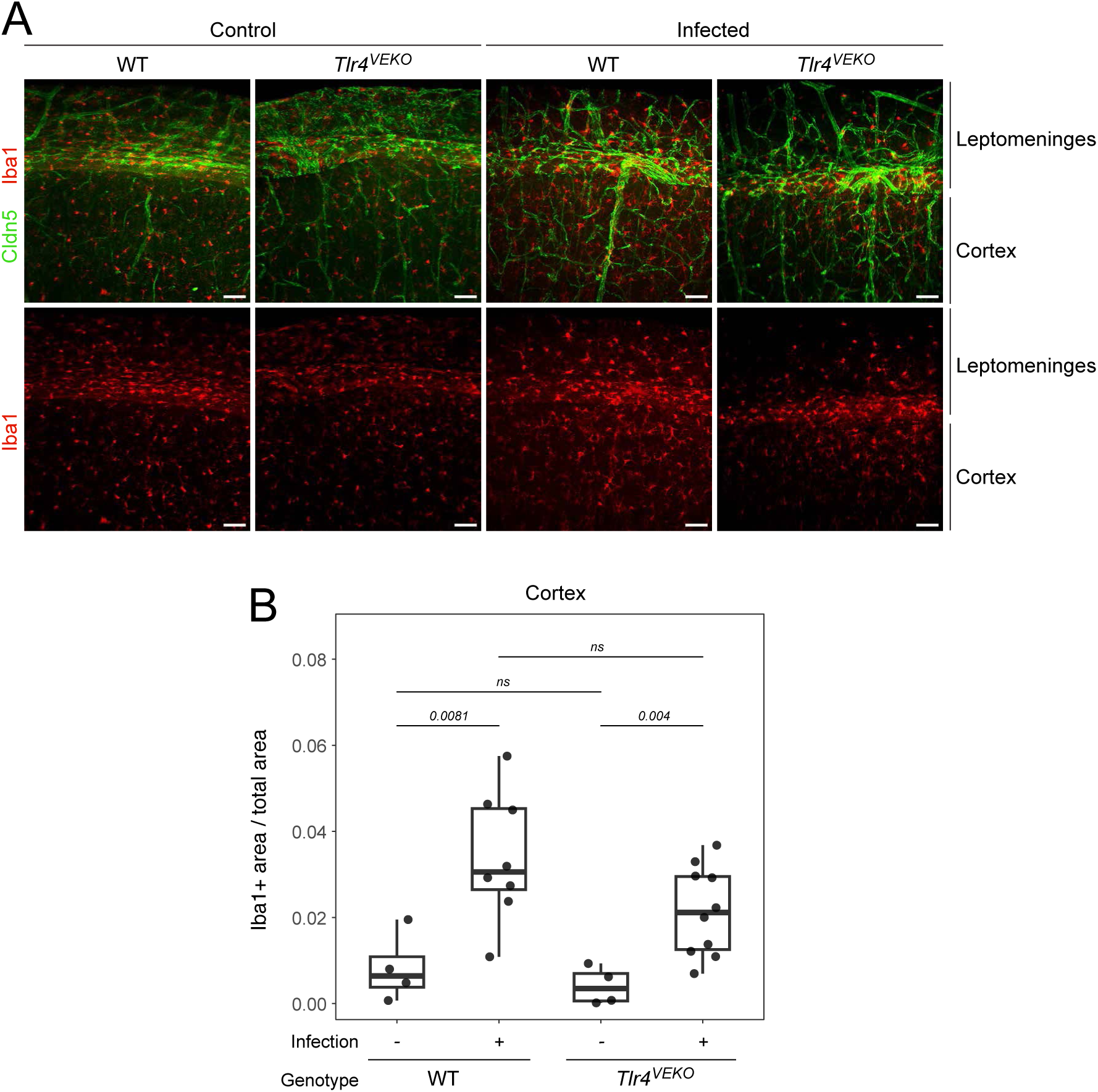
An increase in cortical Iba1+ immunostaining with *E. coli* infection. (A) Coronal sections of P6 cortex and leptomeninges showing Iba1 immunostaining with or without *E. coli* infection in P6 WT and *Tlr4^VEKO^* mice. Scale bars, 50 µm. (B) Quantification of the relative area occupied by Iba1+ pixels in the cortex of WT and *Tlr4^VEKO^* mice with or without *E. coli* infection at P6, as shown in (A).

**Figure 2 – figure supplement 4.**
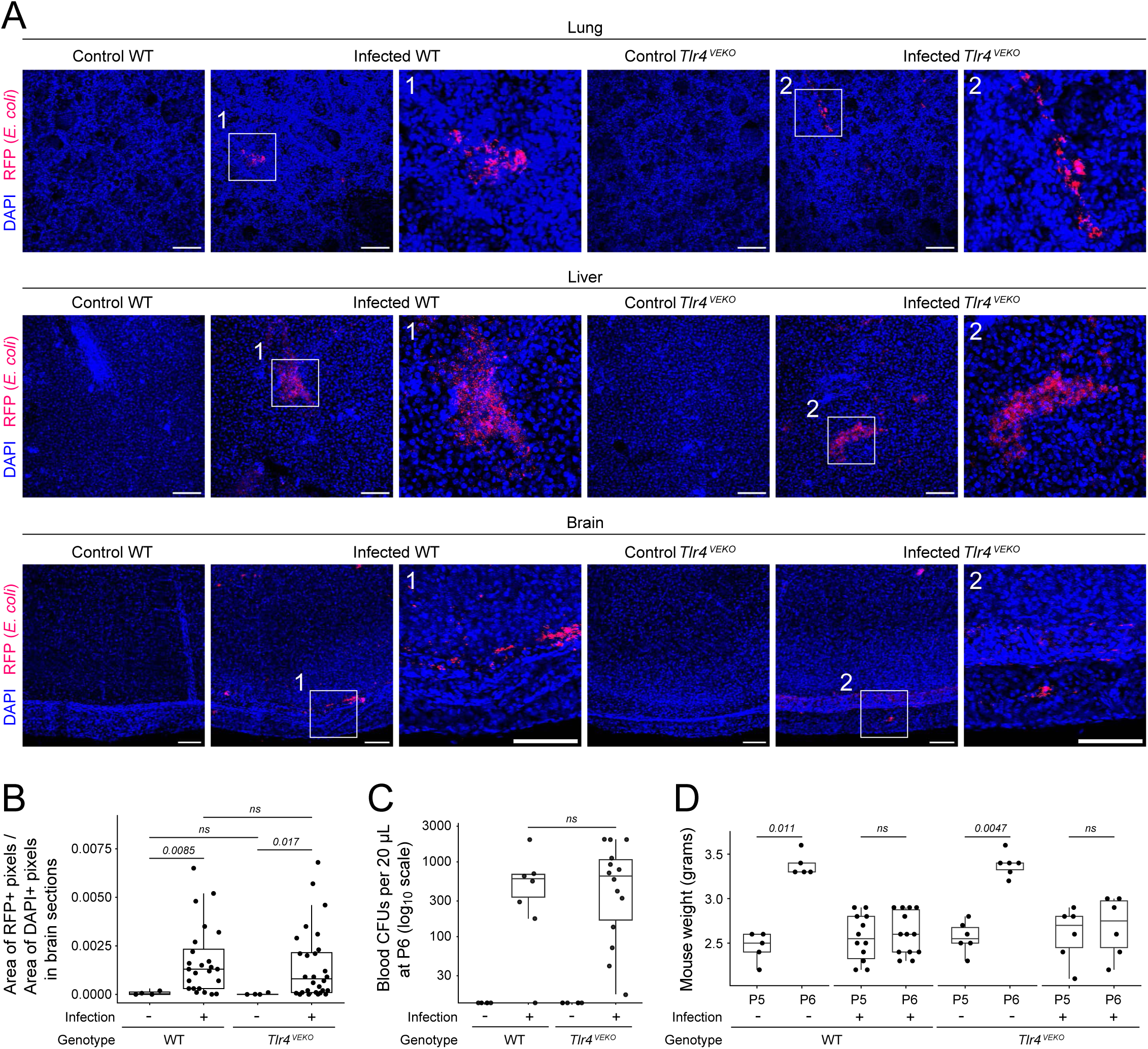
Assessment of clinical progression at 24 hours in WT and *Tlr4^VEKO^* mice following *E. coli* infection. (A) Representative maximum intensity projection images of RFP-expressing *E. coli* (red) and DAPI (blue) in lung, liver, and brain from control (uninfected) and *E. coli*-infected WT and *Tlr4^VEKO^*mice at 24 hours post-infection. Boxed regions (marked 1 and 2) indicate areas of bacterial accumulation. All scale bars, 100 µm. (B) Quantification of bacterial burden in the brain normalized to tissue area, expressed as the ratio of RFP-positive area to DAPI-positive area for each image. Each point represents an individual field of view. Boxes indicate median and interquartile range (IQR); whiskers represent 1.5 x IQR. (C) Bacterial titers in blood measured as colony-forming units (CFUs) per 20 µL of blood at P6 (log₁₀ scale). Each point represents an individual animal. Plates with colony numbers exceeding the reliable counting range were assigned a value of 2,000 CFUs. (D) Body weight measurements at P5 (pre-infection) and P6 (24 hours post-infection). Each point represents an individual animal. For (B-D), statistical comparisons (p-values) were calculated with the Wilcoxon rank-sum test. ns, not significant.

**Figure 3 – figure supplement 1.**
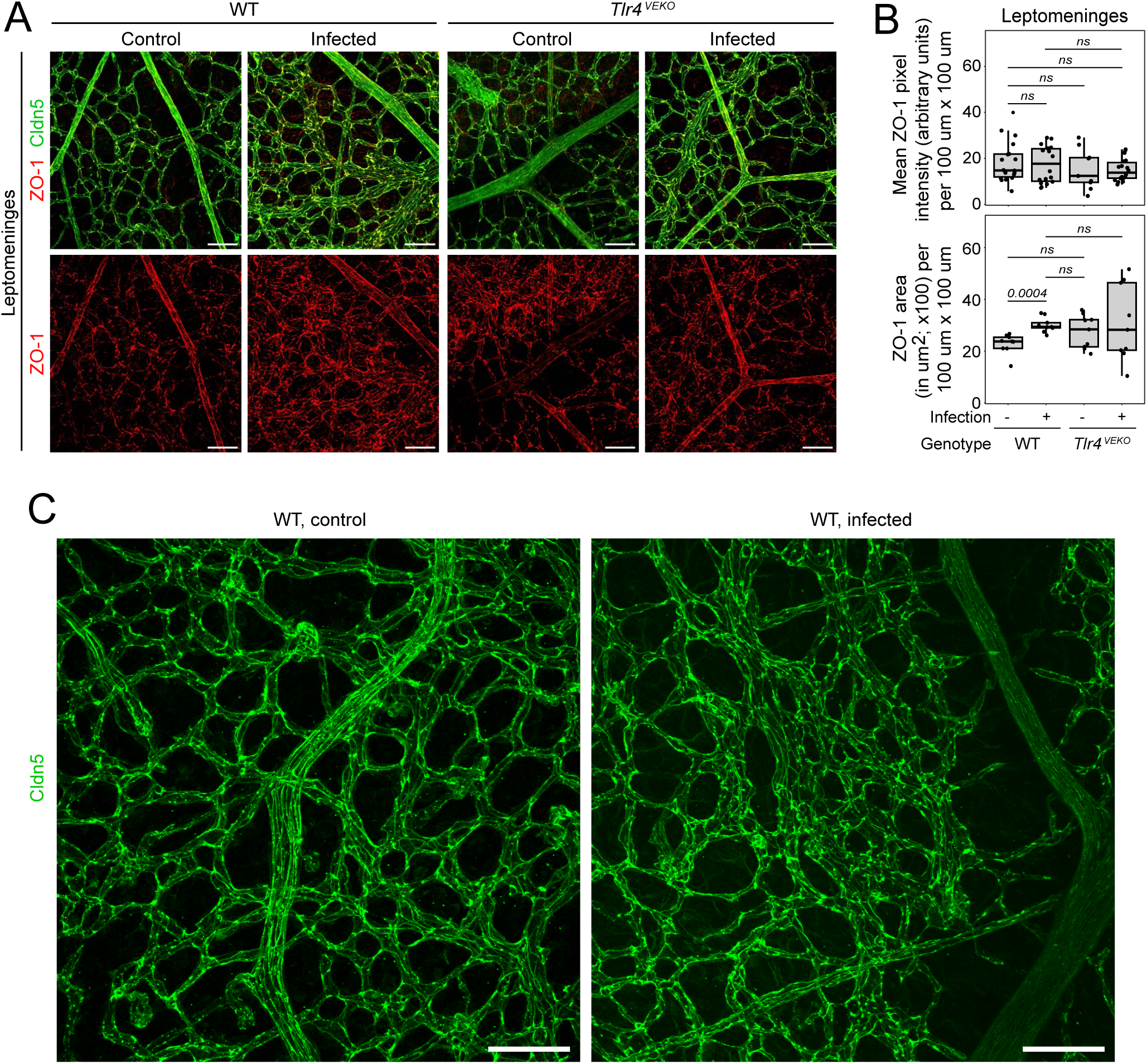
ZO-1 in leptomeningeal endothelial cells with or without *E. coli* infection. (A) Whole mount leptomeninges from P6 WT and *Tlr4^VEKO^*mice, control or *E. coli*-infected, immunostained for Cldn5 and ZO-1. Scale bars, 20 µm. (B) Quantification of ZO-1 immunostaining in the leptomeninges, as shown in (A). (C) Enlarged images of whole mount leptomeningeal vasculature immunostained for Cldn5. Visual inspection reveals a modest elevation in non-junctional Cldn5 in *E. coli*-infected (right) compared to control (left). Scale bar, 50 µm.

**Figure 4 – figure supplement 1.**
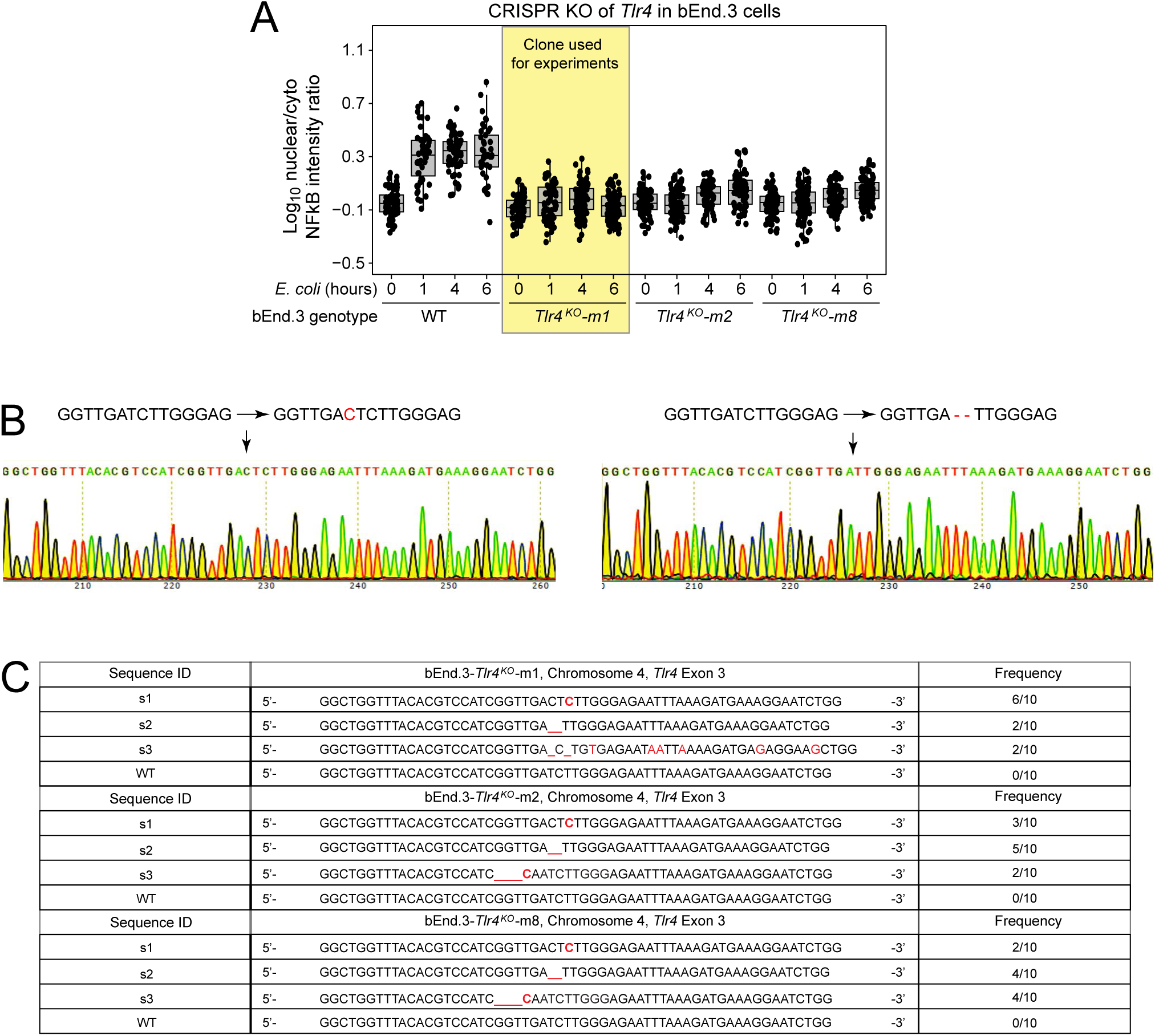
CRISPR knockout of *Tlr4* in bEnd.3 cells. (A) Functional testing of three bEnd.3 clones following CRISPR KO of *Tlr4.* For each clone, the TLR4-mediated NF-κB response to *E. coli* exposure (migration of NF-κB from cytoplasm to nucleus) was quantified by immunostaining, as shown in Figure 3C-E. (B) Sanger sequencing from two plasmid subclones carrying genomic PCR products encompassing the region targeted by the *Tlr4* guide RNA from bEnd.3-*Tlr4^KO^*-m1 cells. Each sequencing run shows a distinctive frameshift mutation. The sequences show the WT sequence on the left and the mutant sequence on the right (with the altered nucleotides in red). Left, a single nucleotide insertion. Right, a two-nucleotide deletion. (C) Compendium of ten plasmid subclone sequences from each of the three bEnd.3 clones shown in (A). Each clone shows three distinct alleles, all of which differ from WT and all of which produce a frameshift. No WT sequences were observed in any subclone. These data imply the *Tlr4* gene is present at three copies in bEnd.3 cells, and that each subclone is likely null for *Tlr4* function.

**Figure 4 – figure supplement 2.**
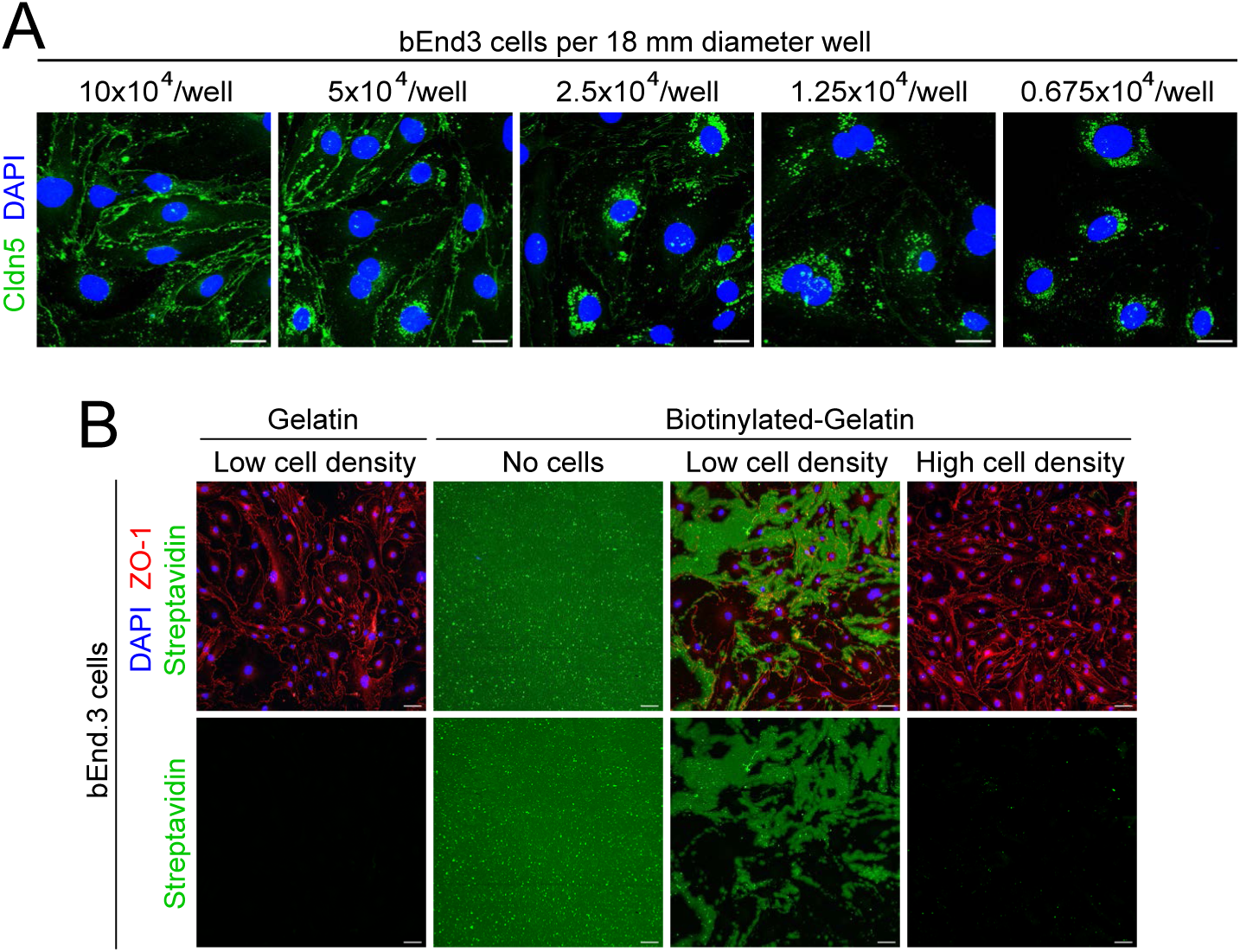
The streptavidin leak assay requires a confluent monolayer of bEnd.3 cells. (A) Among bEnd.3 cells, junctional localization of Cldn5 requires cell-cell contact as shown by immunostaining of cells grown at different densities. Scale bars, 20 µm. (B) bEnd.3 cells were grown at either low or high density on biotinylated gelatin-coated coverslips, incubated with fluorescent streptavidin, and then fixed and immunostained for ZO-1. In regions where the bEnd.3 cells had not formed a contiguous network of tight junctions, the streptavidin gained access to the underlying biotinylated gelatin. Scale bars, 20 µm.

**Figure 5 – figure supplement 1.**
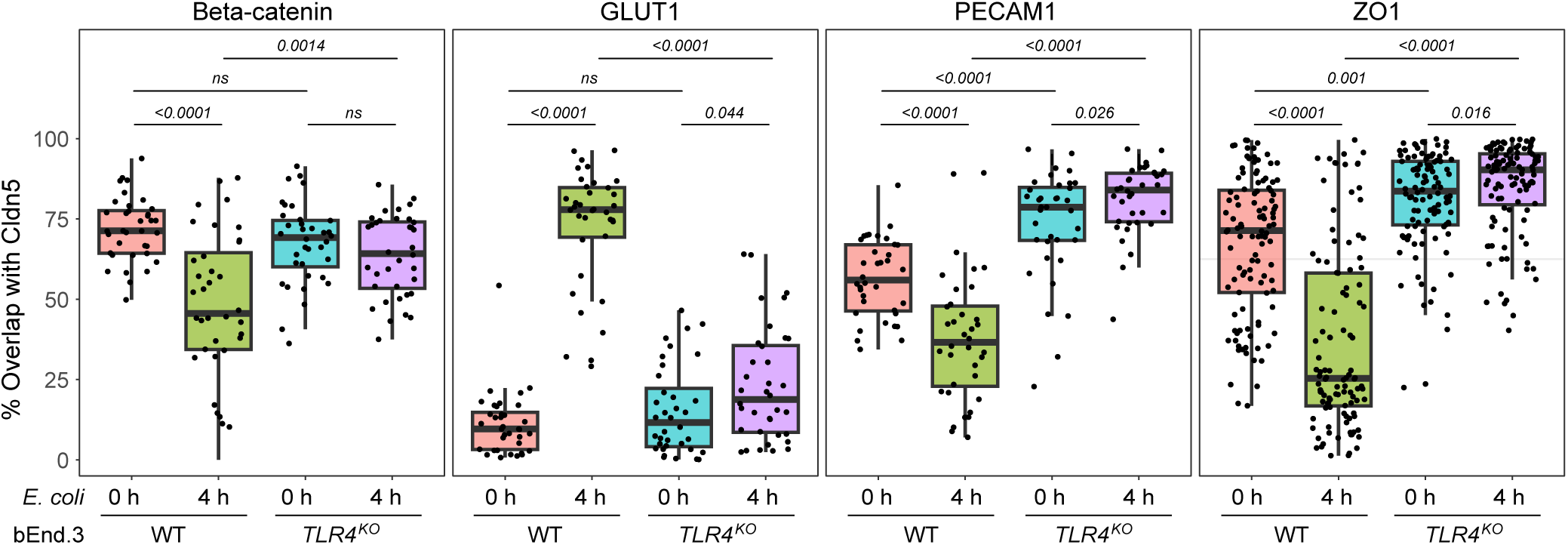
Cldn5 dynamics in bEnd.3 cells following exposure to *E. coli*. The overlap between Cldn5 and β-catenin, GLUT1, PECAM1, and ZO-1, in WT and *Tlr4^KO^* bEnd.3 cells, either not exposed to *E. coli* (control) or exposed to *E. coli* for 4 hours. Shown here is a more granular quantification of the experiment shown in Figure 5A-D, with each datapoint representing a 100 µm x 100 µm region of interest (ROI). Box plots show median, interquartile range, and all individual data points. Each data point represents one image from representative images in three biological replicates. Statistical comparisons (p-values) were calculated with the Wilcoxon rank-sum test.

**Figure 7 – figure supplement 1.**
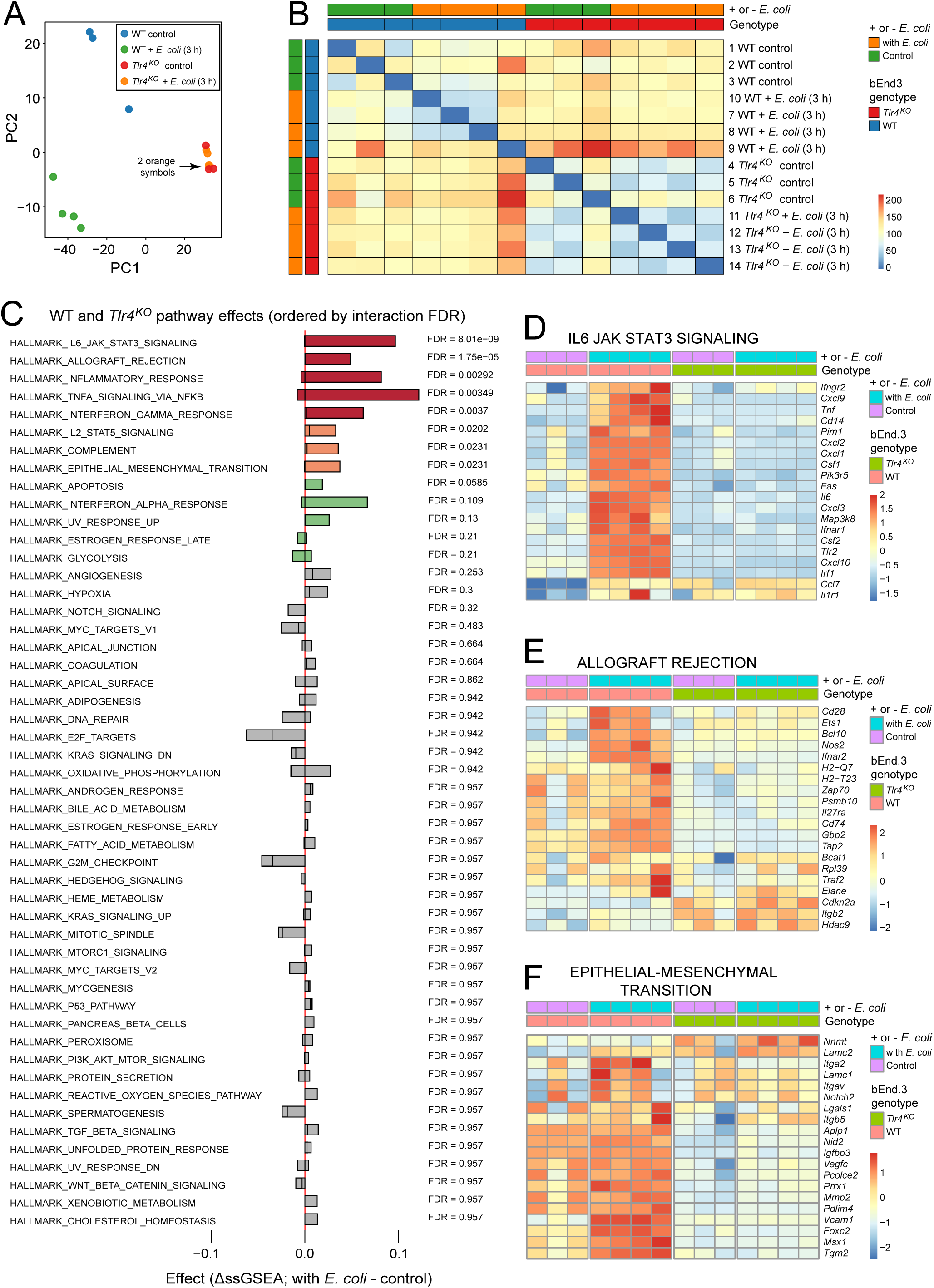
Transcriptome analysis of WT and *Tlr4^KO^* bEnd.3 cells, with or without exposure to *E. coli*. (A) Principal component analysis (PCA) showing the relatedness of the 14 bEnd.3 RNAseq samples. Two of the four orange symbols are nearly superimposed. (B) Pearson correlations for the 14 bEnd.3 RNAseq samples. (C) Pathway-level differences between WT and *Tlr4^KO^* bEnd.3 cells under control and *E. coli*- exposed conditions. Red, false discovery rate (FDR) less than 0.01. Orange, FDR between 0.01 and 0.05. Green, FDR between 0.05 and 0.25. Grey, FDR greater than 0.25. (D-F) Individual transcript abundances in the 14 bEnd.3 RNAseq samples for three GSEA categories. The transcriptome responses to infection are minimal in *Tlr4^KO^* bEnd.3 cells.

## References

Abram CL, Roberge GL, Hu Y, Lowell CA. 2014. Comparative analysis of the efficiency and specificity of myeloid-Cre deleting strains using ROSA-EYFP reporter mice. J Immunol Methods 408:89–100.

Banks WA, Gray AM, Erickson MA, Salameh TS, Damodarasamy M, Sheibani N, Meabon JS, Wing EE, Morofuji Y, Cook DG, Reed MJ. 2015. Lipopolysaccharide-induced blood-brain barrier disruption: roles of cyclooxygenase, oxidative stress, neuroinflammation, and elements of the neurovascular unit. J Neuroinflammation 12:223.

Bernstein NJ, Fong NL, Lam I, Roy MA, Hendrickson DG, Kelley DR. 2020. Solo: Doublet Identification in Single-Cell RNA-Seq via Semi-Supervised Deep Learning. Cell Syst 11:95–101.e5.

Brouwer MC, Tunkel AR, van de Beek D. 2010. Epidemiology, diagnosis, and antimicrobial treatment of acute bacterial meningitis. Clin Microbiol Rev 23:467–492.

Bui TM, Wiesolek HL, Sumagin R. 2020. ICAM-1: A master regulator of cellular responses in inflammation, injury resolution, and tumorigenesis. J Leukoc Biol 108:787–799.

Bundy LM, Rajnik M, Noor A. 2023. Neonatal Meningitis. In StatPearls. Treasure Island, FL (USA): StatPearls Publishing Company,

Casanova JL, Abel L, Quintana-Murci L. 2011. Human TLRs and IL-1Rs in host defense: natural insights from evolutionary, epidemiological, and clinical genetics. Annu Rev Immunol 29:447–491.

Clark IC, Fontanez KM, Meltzer RH, Xue Y, Hayford C, May-Zhang A, D’Amato C, Osman A, Zhang JQ, Hettige P, Ishibashi JSA, Delley CL, Weisgerber DW, Replogle JM, Jost M, Phong KT, Kennedy VE, Peretz CAC, Kim EA, Song S, Karlon W, Weissman JS, Smith CC, Gartner ZJ, Abate AR. 2023. Microfluidics-free single-cell genomics with templated emulsification. Nat Biotechnol 41:1557–1566.

Clausen BE, Burkhardt C, Reith W, Renkawitz R, Förster I. 1999. Conditional gene targeting in macrophages and granulocytes using LysMcre mice. Transgenic Res 8:265–277.

Coisne C, Engelhardt B. 2011. Tight junctions in brain barriers during central nervous system inflammation. Antioxid Redox Signal 15:1285–1303.

Cottarelli A, Jamoul D, Tuohy MC, Shahriar S, Glendinning M, Prochilo G, Edinger AL, Arac A, Agalliu D. 2025. Rab7a is required to degrade select blood-brain barrier junctional proteins after ischemic stroke. Acta Neuropathol Commun 13:203.

Doran KS, Fulde M, Gratz N, Kim BJ, Nau R, Prasadarao N, Schubert-Unkmeir A, Tuomanen EI, Valentin-Weigand P. 2016. Host-pathogen interactions in bacterial meningitis. Acta Neuropathol 131:185–209.

Dubrovskyi O, Birukova AA, Birukov KG. 2013. Measurement of local permeability at subcellular level in cell models of agonist- and ventilator-induced lung injury. Lab Invest 93:254–263.

Edmond K, Clark A, Korczak VS, Sanderson C, Griffiths UK, Rudan I. 2010. Global and regional risk of disabling sequelae from bacterial meningitis: a systematic review and meta- analysis. Lancet Infect Dis 10:317–28.

Engelhardt B, Sorokin L. 2009. The blood-brain and the blood-cerebrospinal fluid barriers: function and dysfunction. Semin Immunopathol 31:497–511.

Fleming SJ, Chaffin MD, Arduini A, Akkad AD, Banks E, Marioni JC, Philippakis AA, Ellinor PT, Babadi M. 2023. Unsupervised removal of systematic background noise from droplet-based single-cell experiments using CellBender. Nat Methods 20:1323–1335.

Franklin BS, Latz E, Schmidt FI. 2018. The intra- and extracellular functions of ASC specks. Immunol Rev 281:74–87.

Freitas RHCN, Fraga CAM. 2018. NF-κB-IKKβ Pathway as a Target for Drug Development: Realities, Challenges and Perspectives. Curr Drug Targets 19:1933–1942.

Gadina M, Johnson C, Schwartz D, Bonelli M, Hasni S, Kanno Y, Changelian P, Laurence A, O’Shea JJ. 2018. Translational and clinical advances in JAK-STAT biology: The present and future of jakinibs. J Leukoc Biol 104:499–514.

Gao Y, Hu F. 2024. Predictive role of PAR and LAR in refractory suppurative meningitis in infants. BMC Pediatr 24:462.

Garcia-Gallardo A, Campbell M. 2025. Understanding the Blood-Brain Barrier: From Physiology to Pathology. Adv Exp Med Biol 1477:1–33.

GBD 2016 Neurology Collaborators. 2019. Global, regional, and national burden of neurological disorders, 1990-2016: a systematic analysis for the Global Burden of Disease Study 2016. Lancet Neurol 18:459–480.

Garges HP, Moody MA, Cotten CM, Smith PB, Tiffany KF, Lenfestey R, Li JS, Fowler VG Jr, Benjamin DK Jr. 2006. Neonatal meningitis: what is the correlation among cerebrospinal fluid cultures, blood cultures, and cerebrospinal fluid parameters? Pediatrics 117:1094–1100.

Guarnera A, Moltoni G, Dellepiane F, Lucignani G, Rossi-Espagnet MC, Campi F, Auriti C, Longo D. 2024. Bacterial Meningoencephalitis in Newborns. Biomedicines 12:2490.

Hafemeister C, Satija R. 2019. Normalization and variance stabilization of single-cell RNA-seq data using regularized negative binomial regression. Genome Biol 20:296.

Hashimoto Y, Campbell M, Tachibana K, Okada Y, Kondoh M. 2021. Claudin-5: A Pharmacological Target to Modify the Permeability of the Blood-Brain Barrier. Biol Pharm Bull 44:1380–1390.

Hashimoto Y, Greene C, Munnich A, Campbell M. 2023. The CLDN5 gene at the blood-brain barrier in health and disease. Fluids Barriers CNS 20:22.

Huang X, Hussain B, Chang J. 2021. Peripheral inflammation and blood-brain barrier disruption: effects and mechanisms. CNS Neurosci Ther 27:36–47.

Huang X, Wei P, Fang C, Yu M, Yang S, Qiu L, Wang Y, Xu A, Hoo RLC, Chang J. 2024. Compromised endothelial Wnt/β-catenin signaling mediates the blood-brain barrier disruption and leads to neuroinflammation in endotoxemia. J Neuroinflammation 21:265.

Hulton CH, Costa EA, Shah NS, Quintanal-Villalonga A, Heller G, de Stanchina E, Rudin CM, Poirier JT. 2020. Direct genome editing of patient-derived xenografts using CRISPR-Cas9 enables rapid in vivo functional genomics. Nat Cancer 1:359–369.

Jia K, Du Y, Lin W, Cao X, Zhang J, Peng L, Li Z, Fang R. 2025. Dual role of TLR4 in bacterial meningitis through regulating endothelial pyroptosis and inflammatory response during extraintestinal pathogenic Escherichia coli infection. Front Immunol 16:1581696.

Jia K, Du Y, Cao X, Shen X, Ran J, Lu Y, Peng L, Li Z, Fang R. 2026. Meningitic Escherichia coli disrupts the blood-brain barrier through pyroptosis and tight junction degradation and NLRP6 deficiency aggravates infection outcomes. Microbiol Res 303:128388.

Johnson I, Spence MTZ. 2010. Probes for Endocytosis, receptors, and ion channels. In The Molecular Probes Handbook: A Guide to Fluorescent Probes and Labeling Technologies. (Life Technologies), pp. 739–772.

Ju, F., Ran, Y., Zhu, L., Cheng, X., Gao, H., Xi, X., Yang, Z., and Zhang, S. 2018. Increased BBB Permeability Enhances Activation of Microglia and Exacerbates Loss of Dendritic Spines After Transient Global Cerebral Ischemia. Front Cell Neurosci 12, 236.

Ju F, Ran Y, Zhu L, Cheng X, Gao H, Xi X, Yang Z, Zhang S. 2018. Increased BBB Permeability Enhances Activation of Microglia and Exacerbates Loss of Dendritic Spines After Transient Global Cerebral Ischemia. Front Cell Neurosci 12:236.

Kawai T, Akira S. 2007. TLR signaling. Semin Immunol 19:24–32.

Kim KS. 2003. Pathogenesis of bacterial meningitis: from bacteraemia to neuronal injury. Nat Rev Neurosci 4:376–385.

Korotkevich G, Sukhov V, Sergushichev A. 2019. Fast gene set enrichment analysis. bioRxiv. DOI: 10.1101/060012

Krishnan S, Chen S, Turcatel G, Arditi M, Prasadarao NV. 2013. Regulation of Toll-like receptor 2 interaction with Ecgp96 controls Escherichia coli K1 invasion of brain endothelial cells. Cell Microbiol 15:63–81.

MacDonald E, Savage B, Zech T. 2020. Connecting the dots: combined control of endocytic recycling and degradation. Biochem Soc Trans 48:2377–2386.

Mapunda JA, Pareja J, Vladymyrov M, Bouillet E, Hélie P, Pleskač P, Barcos S, Andrae J, Vestweber D, McDonald DM, Betsholtz C, Deutsch U, Proulx ST, Engelhardt B. 2023. VE- cadherin in arachnoid and pia mater cells serves as a suitable landmark for in vivo imaging of CNS immune surveillance and inflammation. Nat Commun 14:5837.

Mastorakos P, McGavern D. 2019. The anatomy and immunology of vasculature in the central nervous system. Sci Immunol 4:eaav0492.

McKinsey GL, Lizama CO, Keown-Lang AE, Niu A, Santander N, Larpthaveesarp A, Chee E, Gonzalez FF, Arnold TD. 2020. A new genetic strategy for targeting microglia in development and disease. Elife 9:e54590.

Medzhitov R. 2001. Toll-like receptors and innate immunity. Nat Rev Immunol 1:135–145.

Melville J. 2022. Uwot: the uniform manifold approximation and projection (UMAP) method for Dimensionality reduction. 0.1.14. R Package. https://cran.r-project.org/web/packages/uwot/index.html

Mo A, Mukamel EA, Davis FP, Luo C, Henry GL, Picard S, Urich MA, Nery JR, Sejnowski TJ, Lister R, Eddy SR, Ecker JR, Nathans J. 2015. Epigenomic Signatures of Neuronal Diversity in the Mammalian Brain. Neuron 86:1369–1384.

Montesano R, Pepper MS, Möhle-Steinlein U, Risau W, Wagner EF, Orci L. 1990. Increased proteolytic activity is responsible for the aberrant morphogenetic behavior of endothelial cells expressing the middle T oncogene. Cell 62:435–445.

Monvoisin A, Alva JA, Hofmann JJ, Zovein AC, Lane TF, Iruela-Arispe ML. 2006. VE-cadherin- CreERT2 transgenic mouse: a model for inducible recombination in the endothelium. Dev Dyn 235:3413–3422.

Morita K, Sasaki H, Furuse M, Tsukita S. 1999. Endothelial claudin: claudin-5/TMVCF constitutes tight junction strands in endothelial cells. J Cell Biol 147:185–194.

Nagyoszi P, Wilhelm I, Farkas AE, Fazakas C, Dung NT, Haskó J, Krizbai IA. 2010. Expression and regulation of toll-like receptors in cerebral endothelial cells. Neurochem Int 57:556–564.

Nitta T, Hata M, Gotoh S, Seo Y, Sasaki H, Hashimoto N, Furuse M, Tsukita S. 2003. Size- selective loosening of the blood-brain barrier in claudin-5-deficient mice. J Cell Biol 161:653–660.

Ouchenir L, Renaud C, Khan S, Bitnun A, Boisvert AA, McDonald J, Bowes J, Brophy J, Barton M, Ting J, Roberts A, Hawkes M, Robinson JL. 2017. The Epidemiology, Management, and Outcomes of Bacterial Meningitis in Infants. Pediatrics 140:e20170476.

Pålsson-McDermott EM, O’Neill LA. 2004. Signal transduction by the lipopolysaccharide receptor, Toll-like receptor-4. Immunology 113:153–162.

Pietilä R, Del Gaudio F, He L, Vázquez-Liébanas E, Vanlandewijck M, Muhl L, Mocci G, Bjørnholm KD, Lindblad C, Fletcher-Sandersjöö A, Svensson M, Thelin EP, Liu J, van Voorden AJ, Torres M, Antila S, Xin L, Karlström H, Storm-Mathisen J, Bergersen LH, Moggio A, Hansson EM, Ulvmar MH, Nilsson P, Mäkinen T, Andaloussi Mäe M, Alitalo K, Proulx ST, Engelhardt B, McDonald DM, Lendahl U, Andrae J, Betsholtz C. 2023. Molecular anatomy of adult mouse leptomeninges. Neuron 111:3745–3764.e7.

RStudio Team. 2020. RStudio: Integrated Development Environment for R. RStudio. PBC, Boston, MA. https://rstudio.com

Sanjana NE, Shalem O, Zhang F. 2014. Improved vectors and genome-wide libraries for CRISPR screening. Nat Methods 11:783–784.

Saunders NR, Liddelow SA, Dziegielewska KM. 2012. Barrier mechanisms in the developing brain. Front Pharmacol 3:46.

Schindelin J, Arganda-Carreras I, Frise E, Kaynig V, Longair M, Pietzsch T, Preibisch S, Rueden C, Saalfeld S, Schmid B, Tinevez JY, White DJ, Hartenstein V, Eliceiri K, Tomancak P, Cardona A. 2012. Fiji: an open-source platform for biological-image analysis. Nat Methods 9:676–682.

Scott CC, Vacca F, Gruenberg J. 2014. Endosome maturation, transport and functions. Semin Cell Dev Biol 31:2–10.

Sester DP, Thygesen SJ, Sagulenko V, Vajjhala PR, Cridland JA, Vitak N, Chen KW, Osborne GW, Schroder K, Stacey KJ. 2015. A novel flow cytometric method to assess inflammasome formation. J Immunol 194:455–462.

Shearer LJ, Petersen NO. 2019. Distribution and Co-localization of endosome markers in cells. Heliyon 5:e02375.

Sikorski EE, Hallmann R, Berg EL, Butcher EC. 1993. The Peyer’s patch high endothelial receptor for lymphocytes, the mucosal vascular addressin, is induced on a murine endothelial cell line by tumor necrosis factor-alpha and IL-1. J Immunol 151:5239–5250.

Singh V, Kaur R, Kumari P, Pasricha C, Singh R. 2023. ICAM-1 and VCAM-1: Gatekeepers in various inflammatory and cardiovascular disorders. Clin Chim Acta 548:117487.

Smith NM, Giacci MK, Gough A, Bailey C, McGonigle T, Black AMB, Clarke TO, Bartlett CA, Swaminathan Iyer K, Dunlop SA, Fitzgerald M. 2018. Inflammation and blood-brain barrier breach remote from the primary injury following neurotrauma. J Neuroinflammation 15:201.

Smyth LCD, Xu D, Okar SV, Dykstra T, Rustenhoven J, Papadopoulos Z, Bhasiin K, Kim MW, Drieu A, Mamuladze T, Blackburn S, Gu X, Gaitán MI, Nair G, Storck SE, Du S, White MA, Bayguinov P, Smirnov I, Dikranian K, Reich DS, Kipnis J. 2024. Identification of direct connections between the dura and the brain. Nature 627:165–173.

Sordet O, Rébé C, Plenchette S, Zermati Y, Hermine O, Vainchenker W, Garrido C, Solary E, Dubrez-Daloz L. 2002. Specific involvement of caspases in the differentiation of monocytes into macrophages. Blood 100:4446–4453.

Santambrogio L, Potolicchio I, Fessler SP, Wong SH, Raposo G, Strominger JL. 2005. Involvement of caspase-cleaved and intact adaptor protein 1 complex in endosomal remodeling in maturing dendritic cells. Nat Immunol 6:1020–1028.

Stamatovic SM, Dimitrijevic OB, Keep RF, Andjelkovic AV. 2006. Protein kinase Calpha-RhoA cross-talk in CCL2-induced alterations in brain endothelial permeability. J Biol Chem 281:8379–8388.

Stamatovic SM, Keep RF, Wang MM, Jankovic I, Andjelkovic AV. 2009. Caveolae-mediated internalization of occludin and claudin-5 during CCL2-induced tight junction remodeling in brain endothelial cells. J Biol Chem 284:19053–19066.

Stamatovic SM, Sladojevic N, Keep RF, Andjelkovic AV. 2012. Relocalization of junctional adhesion molecule A during inflammatory stimulation of brain endothelial cells. Mol Cell Biol 32:3414–3427.

Stamatovic SM, Johnson AM, Sladojevic N, Keep RF, Andjelkovic AV. 2017. Endocytosis of tight junction proteins and the regulation of degradation and recycling. Ann NY Acad Sci 1397:54–65.

Stirling, D.R., Swain-Bowden, M.J., Lucas, A.M., Carpenter, A.E., Cimini, B.A., and Goodman, A. 2021. CellProfiler 4: improvements in speed, utility and usability. BMC Bioinformatics 22: ARTN 433.

Subramanian A, Tamayo P, Mootha VK, Mukherjee S, Ebert BL, Gillette MA, Paulovich A, Pomeroy SL, Golub TR, Lander ES, Mesirov JP. 2005. Gene set enrichment analysis: a knowledge-based approach for interpreting genome-wide expression profiles. Proc Natl Acad Sci U S A 102:15545–15550.

Sukumaran SK, Prasadarao NV. 2003. Escherichia coli K1 invasion increases human brain microvascular endothelial cell monolayer permeability by disassembling vascular-endothelial cadherins at tight junctions. J Infect Dis 188:1295–309.

Too LK, Yau B, Baxter AG, McGregor IS, Hunt NH. 2019. Double deficiency of toll-like receptors 2 and 4 alters long-term neurological sequelae in mice cured of pneumococcal meningitis. Sci Rep 9:16189.

Vallières L, Rivest S. 1997. Regulation of the genes encoding interleukin-6, its receptor, and gp130 in the rat brain in response to the immune activator lipopolysaccharide and the proinflammatory cytokine interleukin-1beta. J Neurochem 69:1668–1683.

van Well GT, Sanders MS, Ouburg S, Kumar V, van Furth AM, Morré SA. 2013. Single nucleotide polymorphisms in pathogen recognition receptor genes are associated with susceptibility to meningococcal meningitis in a pediatric cohort. PLoS One 8:e64252.

Wang J, Rattner A, Nathans J. 2023. Bacterial meningitis in the early postnatal mouse studied at single-cell resolution. Elife 12:e86130.

Wang X, Zhang Y, Chen X, Fu K, Cui J, Wu J, Sun Y, Ren J, Xue F, Dai J, Tang F. 2025a. RIPK1 kinase drove brain microvascular endothelial cells death and blood-brain barrier disruption in neonatal Escherichia coli meningitis. Nat Commun 16:7309.

Wang Y, Cho C, Williams J, Smallwood PM, Zhang C, Junge HJ, Nathans J. 2018. Interplay of the Norrin and Wnt7a/Wnt7b signaling systems in blood-brain barrier and blood-retina barrier development and maintenance. Proc Natl Acad Sci U S A 115:E11827–E11836.

Wang Y, Sabbagh MF, Gu X, Rattner A, Williams J, Nathans J. 2019. Beta-catenin signaling regulates barrier-specific gene expression in circumventricular organ and ocular vasculatures. Elife 8:e43257.

Wang Y, Rattner A, Li Z, Smallwood PM, Nathans J. 2025b. Vascular endothelial-specific loss of TGF-beta signaling as a model for choroidal neovascularization and central nervous system vascular inflammation. Elife 14:RP107018.

Wei C, Jiang W, Wang R, Zhong H, He H, Gao X, Zhong S, Yu F, Guo Q, Zhang L, Schiffelers LDJ, Zhou B, Trepel M, Schmidt FI, Luo M, Shao F. 2024. Brain endothelial GSDMD activation mediates inflammatory BBB breakdown. Nature 629:893–900.

Wickham H. 2009. Ggplot2: Elegant Graphics for Data Analysis. New York: Springer-Verlag.

Wickham H. 2017. tidyverse: Easily Install and Load the ‘Tidyverse’. 1.2.1. R Package. https://CRAN.R-project.org/package=tidyverse

Williams RL, Courtneidge SA, Wagner EF. 1988. Embryonic lethalities and endothelial tumors in chimeric mice expressing polyoma virus middle T oncogene. Cell 52:121–31.

Xu M, Hu L, Huang H, Wang L, Tan J, Zhang Y, Chen C, Zhang X, Huang L. 2019. Etiology and Clinical Features of Full-Term Neonatal Bacterial Meningitis: A Multicenter Retrospective Cohort Study. Front Pediatr 7:31.

Yang R, Wang J, Wang F, Zhang H, Tan C, Chen H, Wang X. 2023. Blood-Brain Barrier Integrity Damage in Bacterial Meningitis: The Underlying Link, Mechanisms, and Therapeutic Targets. Int J Mol Sci 24:2852.

Yang R, Chen J, Qu X, Liu H, Wang X, Tan C, Chen H, Wang X. 2024. Interleukin-22 Contributes to Blood-Brain Barrier Disruption via STAT3/VEGFA Activation in Escherichia coli Meningitis. ACS Infect Dis 10:988–999.

Zaffaroni L, Peri F. 2018. Recent advances on Toll-like receptor 4 modulation: new therapeutic perspectives. Future Med Chem 10:461–476.

Zhu N, Liu W, Prakash A, Zhang C, Kim KS. 2020. Targeting E. coli invasion of the blood-brain barrier for investigating the pathogenesis and therapeutic development of E. coli meningitis. Cell Microbiol 22:e13231.

