## Supplemental File 1 for "TLR4 signaling drives tissue inflammation, Claudin-5 internalization, and vascular barrier breakdown in a mouse model of neonatal meningitis"

**Supplemental File 1. snRNAseq statistics**

| Sample (batch) | genotype | condition | number of nuclei | mean number of transcripts per nucleus | mean number of genes per nucleus |
|---|---|---|---|---|---|
| PS3 (2) | *Tlr4*$^{MKO}$ | infected | 8890 | 1004.26 | 500.83 |
| PS12 (5) | *Tlr4*$^{MKO}$ | infected | 9687 | 928.43 | 459.64 |
| PS17 (6) | *Tlr4*$^{MKO}$ | infected | 6225 | 646.25 | 349.69 |
| PS14 (5) | *Tlr4*$^{MKO}$ | uninfected | 9356 | 894.39 | 455.57 |
| PS18 (6) | *Tlr4*$^{MKO}$ | uninfected | 9441 | 777.10 | 406.87 |
| PS1 (1) | *Tlr4*$^{ECKO}$ | infected | 7615 | 1410.12 | 665.22 |
| PS11 (5) | *Tlr4*$^{ECKO}$ | infected | 7687 | 649.76 | 364.03 |
| PS15 (6) | *Tlr4*$^{ECKO}$ | infected | 9142 | 708.67 | 426.62 |
| PS9 (4) | *Tlr4*$^{ECKO}$ | uninfected | 8248 | 927.29 | 467.43 |
| PS10 (4) | *Tlr4*$^{ECKO}$ | uninfected | 7549 | 814.97 | 432.43 |
| PS2 (1) | *Tlr4*$^{CKO/-}$ (=control) | infected | 7964 | 1969.53 | 1055.53 |
| PS6 (3) | *Tlr4*$^{CKO/-}$ (=control) | infected | 8133 | 986.80 | 527.68 |
| PS13 (5) | *Tlr4*$^{CKO/-}$ (=control) | infected | 8458 | 650.66 | 359.79 |
| PS4 (2) | *Tlr4*$^{CKO/-}$ (=control) | uninfected | 10953 | 880.44 | 480.63 |
| PS8 (3) | *Tlr4*$^{CKO/-}$ (=control) | uninfected | 8853 | 830.12 | 417.12 |
| Average | | | 8547 | 935.00 | 489.60 |
| total | | | 128,201 | | |
