## Supplemental File 2 for "TLR4 signaling drives tissue inflammation, Claudin-5 internalization, and vascular barrier breakdown in a mouse model of neonatal meningitis"

**Supplementary File 2. Number of mice and numbers of images per experiment**

### Number of mice per genotype for immunostaining analysis

| Group | ASC | CD206 | Casp3 | Cldn5 | ICAM1 | Streptavidin | ZO1 | Iba1 | RFP-*E. coli* | Sun1-GFP | R26-tdT-2A-GFP |
|---|---|---|---|---|---|---|---|---|---|---|---|
| WT, Uninfected | 2 | 2 | 2 | 3 | 2 | 2 | 2 | 2 | 2 | 0 | 0 |
| WT, Infected | 6 | 7 | 6 | 8 | 7 | 5 | 4 | 6 | 6 | 0 | 0 |
| $TLR4^{VEcko}$, Uninfected | 1 | 1 | 2 | 2 | 2 | 1 | 1 | 2 | 2 | 0 | 0 |
| $TLR4^{VEcko}$, Infected | 4 | 4 | 3 | 10 | 6 | 5 | 7 | 9 | 9 | 0 | 0 |
| $TLR4^{Mcko}$, Uninfected | 0 | 0 | 2 | 2 | 1 | 1 | 0 | 0 | 0 | 0 | 0 |
| $TLR4^{Mcko}$, Infected | 4 | 3 | 3 | 6 | 5 | 2 | 2 | 0 | 0 | 0 | 0 |
| Cdh5CreER/+ Control | 0 | 0 | 0 | 0 | 0 | 0 | 0 | 0 | 0 | 3 | 3 |
| Lyz2cre/+ Control | 0 | 0 | 0 | 0 | 0 | 0 | 0 | 0 | 0 | 3 | 3 |

### Number of images per genotype for immunostaining analysis

| Group | ASC | CD206 | Casp3 | Cldn5 | ICAM1 | Streptavidin | ZO1 | Iba1 | RFP-*E. coli* | Sun1-GFP | R26-tdT-2A-GFP |
|---|---|---|---|---|---|---|---|---|---|---|---|
| WT, Uninfected | 8 | 8 | 3 | 51 | 18 | 16 | 7 | 4 | 4 | 0 | 0 |
| WT, Infected | 20 | 32 | 22 | 120 | 43 | 34 | 9 | 12 | 12 | 0 | 0 |
| $TLR4^{VEcko}$, Uninfected | 4 | 6 | 4 | 21 | 7 | 7 | 2 | 4 | 4 | 0 | 0 |
| $TLR4^{VEcko}$, Infected | 19 | 11 | 19 | 74 | 21 | 28 | 13 | 18 | 18 | 0 | 0 |
| $TLR4^{Mcko}$, Uninfected | 0 | 0 | 2 | 8 | 2 | 4 | 0 | 0 | 0 | 0 | 0 |
| $TLR4^{Mcko}$, Infected | 8 | 6 | 4 | 37 | 13 | 8 | 6 | 0 | 0 | 0 | 0 |
| Cdh5CreER/+ Control | 0 | 0 | 0 | 0 | 0 | 0 | 0 | 0 | 0 | 30 | 30 |
| Lyz2cre/+ Control | 0 | 0 | 0 | 0 | 0 | 0 | 0 | 0 | 0 | 3 | 3 |
